# Trans-omic analysis reveals opposite metabolic dysregulation between feeding and fasting in liver associated with obesity

**DOI:** 10.1101/2023.11.15.567316

**Authors:** Yunfan Bai, Keigo Morita, Toshiya Kokaji, Atsushi Hatano, Satoshi Ohno, Riku Egami, Yifei Pan, Dongzi Li, Katsuyuki Yugi, Saori Uematsu, Hiroshi Inoue, Yuka Inaba, Yutaka Suzuki, Masaki Matsumoto, Masatomo Takahashi, Izumi Yoshihiro, Takeshi Bamba, Akiyoshi Hirayama, Tomoyoshi Soga, Shinya Kuroda

## Abstract

Dysregulation of liver metabolism associated with obesity during feeding and fasting leads to the breakdown of metabolic homeostasis. However, the underlying mechanism remains unknown. Here, we measured multi-omics data in the liver of wild-type and leptin-deficient obese (*ob*/*ob*) mice at *ad libitum* feeding, and constructed a differential regulatory trans-omic network of metabolic reactions. We compared the trans-omic network at feeding with that at 16 h-fasting constructed in our previous study. Intermediate metabolites in glycolytic and nucleotide metabolism decreased in *ob*/*ob* mice at feeding but increased at fasting. Allosteric regulation reversely shifted between feeding and fasting, generally showing activation at feeding while inhibition at fasting in *ob*/*ob* mice. Transcriptional regulation was similar between feeding and fasting, generally showing inhibiting transcription factor regulations, activating enzyme protein regulations in *ob*/*ob* mice. The opposite metabolic dysregulation between feeding and fasting characterizes breakdown of metabolic homeostasis associated with obesity.

## INTRODUCTION

The liver is a central metabolic workhouse that tightly governs whole-body metabolic homeostasis^1–3^. By regulating different metabolic pathways in response to feeding or fasting, liver provides fuel sources for the whole body and maintains stability of metabolite concentrations. At feeding, glycolysis in liver predominantly provides energy for the whole body, given that glucose is abundant. At fasting, the liver absorbs alanine, lactate, and glycerol from the blood and transforms them into glucose by gluconeogenesis. Fatty acid oxidation provides energy with accumulated ketone bodies in hepatic mitochondria^4^. Allosteric regulation is a flexible and dynamic mechanism to modulate enzyme protein activity^5–7^. Many studies have shown different metabolic pathways in the liver are regulated by transcriptional mechanism and allosteric regulators^8–11^. For example, transcription factor, sterol regulatory element binding protein 1c (SREBP-1c) and carbohydrate response element binding protein (ChREBP) are responsible for activation of glycolytic and fatty acid biosynthesis enzymes under insulin and glucose stimulation^12^. The rate-limiting enzymes in glycolysis and gluconeogenesis pathway, such as Pfk1, G6pase, and Fbpase1 are allosterically regulated by metabolites like ATP, ADP, citrate, F26BP^13^. However, a systematic overview of liver metabolism during feeding and fasting from upstream (insulin signals, TFs, protein kinases) to downstream (mRNAs, proteins, phosphoproteins, metabolites, and lipids) using multi-omics data has yet to be conducted and analyzed.

Obesity and over-feeding derange hepatic metabolism^14–16^. Not only derangement of fasting hepatic metabolism but also of feeding plays an important role in the pathogenesis of lifestyle diseases. Postprandial hyperglycemia and hypertriglyceridemia caused by dysregulated hepatic metabolism in obese subjects are closely related to onset of Type 2 Diabetes. Thus, study on metabolism in obese individuals taking a natural amount of food, *i*.*e*., *ad libitum* feeding, provides a prototype to understand how feeding affects hepatic metabolism in obesity. Based on this, we could reveal how metabolism adjusting between feeding and fasting is dysregulated in obesity, which is a critical feature of metabolic homeostasis. Although some studies reported how metabolism changes in the liver of obesity at fasting or high-fat diet by using omics data^17,18^, how liver metabolism differently responds to feeding and fasting associated with obesity has not directly been compared and analyzed yet.

Allosteric regulator, serving as an activator or inhibitor depending on the target enzyme, binds or disturbs a distal site in a protein and caused its conformational change. Some databases such as BRaunschweig ENzyme DAtabase (BRENDA) provide detailed information on metabolites and small molecules function as activators or inhibitors interacting with enzyme proteins, which is widely used in allostery analysis^19–21^. Constructing metabolic networks using simultaneously measured multi-omics data is an efficient way to understand the global picture of dysregulation of metabolism associated with obesity. The core of metabolic network, metabolic reactions, can be regulated by the following events: phosphorylation of enzyme proteins via protein kinases^22–24^; change in amount of enzyme proteins, which TFs controlled through the regulation of gene expression; allosteric regulations via metabolites or lipids; substrate or product interaction. Insulin and other metabolic hormones mediate downstream metabolic events, including TFs and protein kinases activity^5,25–29^. Previously, we proposed the concept of “trans-omic”, a global biochemical network for revealing molecular interactions and regulations^30–33^. We constructed a differential regulatory trans-omic network of metabolic reactions in the liver of wild-type (WT) and *ob*/*ob* mice at 16 h-fasting^34^. The trans-omic network showed more activating enzyme protein regulations, and more inhibiting allosteric regulations in *ob*/*ob* mice at fasting on many metabolic pathways, such as glycolysis/gluconeogenesis, fatty acid degradation. In this study, we constructed differential regulatory trans-omic network at feeding, and took this as a prototype to investigate the breakdown of liver metabolic homeostasis adjusting between feeding and fasting associated with obesity.

In this study, we measured multi-omics data and constructed a differential regulatory trans-omic network of metabolic reactions in the liver between WT and *ob*/*ob* mice at *ad libitum* feeding. We compared the trans-omic network at *ad libitum* feeding with that constructed at 16 h-fasting in our previous study^34^. We found opposite metabolic dysregulation on metabolites and allosteric regulations between *ad libitum* feeding and 16 h-fasting in the liver associated with obesity. Intermediate metabolites in carbohydrate pathways (glycolysis/gluconeogenesis, the TCA cycle), triphosphates in nucleotide pathways decreased in *ob*/*ob* mice at *ad libitum* feeding while increased at 16 h-fasting. Allosteric regulations were influenced by feeding and fasting, generally showing activation in *ob*/*ob* mice at *ad libitum* feeding while inhibition at 16 h-fasting. Transcriptional regulation (from TFs to enzyme proteins to metabolic reactions) was similar between *ad libitum* feeding and 16 h-fasting. This study captures the global picture of the dysregulation of metabolic homeostasis adjusting between feeding and fasting in the liver associated with obesity.

## RESULTS

### Overview of the approach in this study

We measured transcriptome (mRNAs), proteome (proteins), phosphoproteome (phosphoproteins), insulin signaling molecules (phosphoproteins), metabolome (metabolites), and lipidome (lipids). Based on these multi-omics data and bioinformatics databases, we constructed a differential regulatory trans-omic network of metabolic reactions in the liver of WT and *ob*/*ob* mice at *ad libitum* feeding, and compared it with that at 16 h-fasting, which we have previously constructed^34^ (Figure 1 and Figure S1). We used the liver and blood of 10-week-old male WT and *ob*/*ob* mice at *ad libitum* feeding (n = 5). Glucose and insulin concentrations in the blood of *ob*/*ob* mice were significantly higher than those of WT mice (p < 0.001), indicating that *ob*/*ob* mice had hyperglycemia and hyperinsulinemia (Figure S1A). The analysis in this study included the following five steps. In Step 1, we identified differentially expressed or phosphorylated molecules that significantly increased (red nodes) or decreased (blue nodes) in *ob*/*ob* mice compared with WT mice in each omic layer. In Step 2, we identified the differential regulations of gene expression regulated by potential transcription factors (TFs) using the ChIP-Atlas database^35^, differential protein phosphorylation mediated by potential protein kinases using the Netphorest database^36,37^, metabolic reactions regulated by enzyme, substrate or product using the Kyoto Encyclopedia of Genes and Genomes (KEGG) database^38^, and allosteric regulations using the BRaunschweig ENzyme DAtabase (BRENDA) databases^19^. Differential regulations were classified into two groups, activating regulation in *ob*/*ob* mice (red arrows), and inhibiting regulation in *ob*/*ob* mice (blue arrows). In Step 3, we constructed the differential regulatory trans-omic network of metabolic reactions in the liver by integrating differentially expressed or phosphorylated molecules and differential regulations (Figure S1D). In Step 4, we condensed the trans-omic network and extracted specific metabolic pathways from the trans-omic network that highlighted metabolic dysregulation in *ob*/*ob* mice. In Step 5, we compared the differential regulatory trans-omic network of *ad libitum* feeding with that at 16 h-fasting^34^.

**Figure 1.**
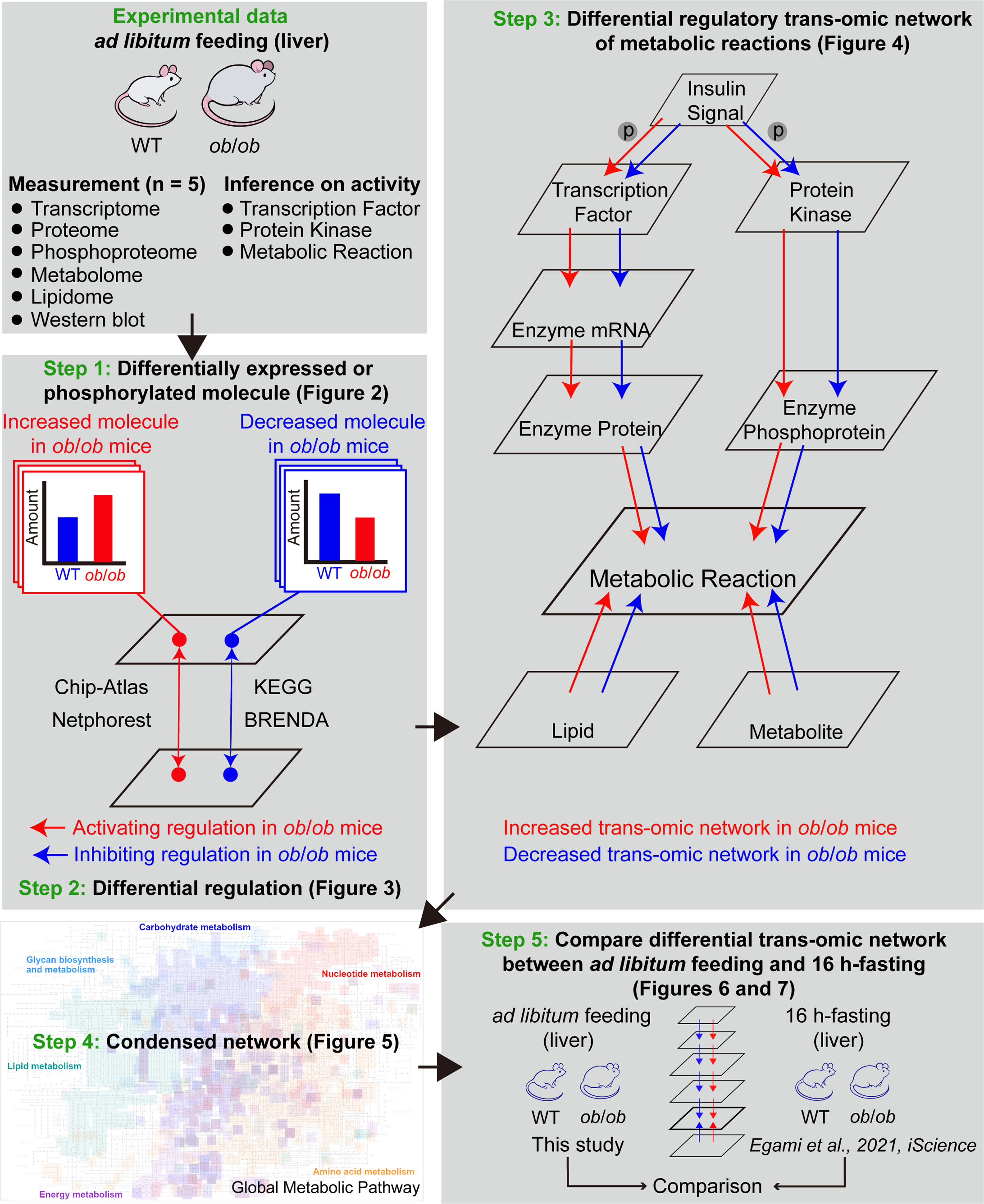
Overview of the strategy for constructing the differential regulatory trans-omic network of metabolic reactions from the liver of WT and *ob*/*ob* mice at *ad libitum* feeding. We measured the multi-omics data in the liver of WT and *ob*/*ob* mice at *ad libitum* feeding (n = 5 mice per genotype), identified differentially expressed or phosphorylated molecules (Step 1), differential regulation (Step 2), constructed a differential regulatory trans-omic network (Step 3), made a condensed network (Step 4), and compared it with that at 16 h-fasting, which we have previously constructed^34^ (Step 5). The letter “p” in the circle in Step 3 indicates the regulation by protein phosphorylation. TF, mRNA, and protein kinase names are italicized. Phosphoprotein names are shown with prefix “p”. Red, increased molecules or activating regulations in *ob*/*ob* mice; blue, decreased molecules or inhibiting regulations in *ob*/*ob* mice. See also Figure S1 for the details of Steps 1, 2, and 3.

### Identification of differentially expressed molecules in *ob*/*ob* mice at *ad libitum* feeding

We identified molecules (mRNAs, proteins, phosphoproteins, metabolites, lipids, TFs, and protein kinases) that differentially expressed or phosphorylated between WT and *ob*/*ob* mice in the liver at *ad libitum* feeding (Figure 2).

**Figure 2.**
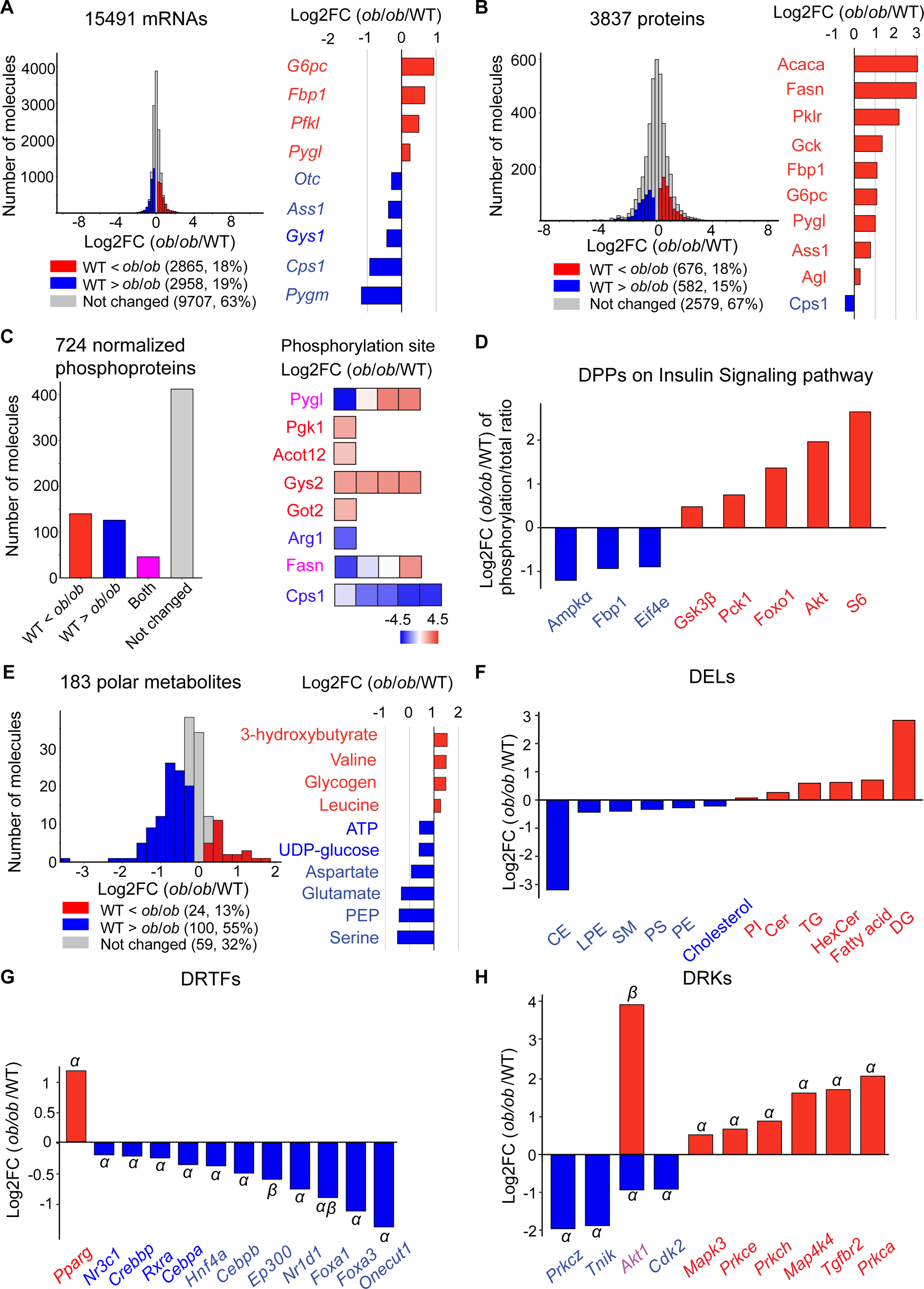
Identification of differentially expressed or phosphorylated molecules in the liver between WT and *ob*/*ob* mice at *ad libitum* feeding. (A) Histogram of log2 fold changes (FCs) of mRNA expression values and the log2 FCs of the indicated DEGs (right). See also Table S1 and Figure S2A. (B) Histogram of log2 FCs of protein expression values (left) and the log2 FCs of the indicated DEPs (right). See also Table S2 and Figure S2A. (C) Bar plot of the increased, decreased, and both increased and decreased DPPs (left) and the log2 FC of phosphorylation level in each phosphorylation site (square) of the indicated DPP, color indicates log2 FC value (right). We calculated the log2 FCs of phosphorylation sites first. If all sites increased or decreased in one phosphoprotein (q < 0.1), this phosphoprotein is identified as the increased or decreased DPP in *ob*/*ob* mice. If some phosphorylation sites increased while others decreased, this phosphoprotein is identified as the increased and decreased DPP in *ob*/*ob* mice (magenta). See also Table S3 and Figures S2B and S2C. (D) The log2 FCs of DPPs on insulin signaling pathway. See also Tables S3 and S4, and Figures S2D and S2E. (E) Histogram of log2 FCs of amounts of polar metabolites (left) and the log2 FCs of indicated DEMs (right). See also Table S5 and Figures S2F and S2G. (F) The log2 FCs of DELs. See also Table S6 and Figure S2H. (G) The log2 FCs of mRNA expression or protein phosphorylation values of inferred DETFs or DPTFs. *α*, DETF; *β*, DPTF. *Foxa1* was inferred as both decreased DETF and DPTF. The mean of log2 FC of mRNA expression and protein phosphorylation values indicated the log2 FC. See also Table S7. (H) The log2 FCs of mRNA expression or protein phosphorylation values of inferred DEKs or DPKs. *α*, DEK; *β*, DPK. *Akt1* was inferred as the decreased DEK and increased DPK. Both log2 FC of mRNA expression and protein phosphorylation values were shown. See also Table S8. For each molecule, the significance of difference between WT and *ob*/*ob* mice (p value) was tested by two-tailed Welch’s t-test, q value was calculated by Storey procedure. Red, increased molecules in *ob*/*ob* mice; blue, decreased molecules in *ob*/*ob* mice; magenta, some phosphoproteins contain increased and decreased phosphorylation sites in *ob*/*ob* mice; gray, unchanged molecules between WT and *ob*/*ob* mice. Log2 FC of each detected molecule was calculated by dividing the expression value of *ob*/*ob* mice to WT mice. We identified increased molecules in *ob*/*ob* mice (log2 FC>0 and q < 0.1) and decreased molecules in *ob*/*ob* mice (log2 FC<0 and q < 0.1). Data are shown as the mean of mice replicates (n = 5 mice per genotype). Note that error bars are not provided because the FC of averaged values of WT and *ob*/*ob* mice were used to calculate log2 FC between WT and *ob*/*ob* mice.

For the transcriptome, we measured the expression of 15,491 mRNAs using RNA sequencing (Figure 2A; Table S1). We identified mRNAs that showed significantly different expression in *ob*/*ob* mice compared with WT mice as differentially expressed genes (DEGs). We identified 2,865 (18%) increased and 2,958 (19%) decreased DEGs in *ob*/*ob* mice compared to WT mice (Figure 2A). We observed the increased DEGs involved in glycolysis/gluconeogenesis (glucose-6-phosphatase [*G6pc*], fructose-bisphosphatase 1 [*Fbp1*]), and the decreased DEGs involved in arginine synthesis (urea cycle) (carbamoyl-phosphate synthase 1 [*Cps1*], ornithine transcarbamylase [*Otc*]).

For the proteome, we quantitatively measured the expression of 3,837 proteins using the liver lysate of stable isotope labeling by amino acids in cell culture (SILAC)-mouse as an internal control (Figure 2A; Table S2). We identified proteins that showed significantly different expression in *ob*/*ob* mice compared with WT mice as differentially expressed proteins (DEPs). We identified 676 (18%) increased and 582 (15%) decreased DEPs in *ob*/*ob* mice compared to WT mice (Figure 2B). The increased DEPs included the metabolic enzymes in glycolysis/gluconeogenesis (pyruvate kinase liver [Pklr], glucokinase [Gck], Fbp1, G6pc) and fatty acid synthesis (acetyl-CoA carboxylase alpha [Acaca], fatty acid synthase [Fasn]). Cps1, which is involved in arginine synthesis, decreased in *ob/ob* mice.

For the phosphoproteome, we used the corresponding protein levels to normalize phosphoprotein levels at each phosphorylation site by dividing the total amount of proteins^39^. After normalization, we obtained 2,434 phosphorylation sites (Figure S2B). We identified proteins that showed significantly different protein phosphorylation at least one phosphorylation site in *ob*/*ob* mice compared with WT mice as differentially phosphorylated proteins (DPPs). We identified 140 (19%) increased, 126 (18%) decreased, and 46 (6%) both increased and decreased DPPs in *ob*/*ob* mice compared to WT mice based on the criterion (Figures 2C, S2B, and S2C; Table S3). DPPs related to carbohydrate metabolism (phosphorylated Pygl [pPygl], phosphorylated phosphoglycerate kinase 1 [pPgk1], and phosphorylated glycogen synthase 2 [pGys2]) contained increased differentially phosphorylated phosphorylation sites (DPSs) (Figure 2C). We identified 15 DPPs on the insulin signaling pathway by western blot analysis and phosphoproteome data (Figures 2D, S2D and S2E; Table S4), including increased phosphorylated glycogen synthase 3 beta (pGsk3ꞵ), phosphorylated forkhead box protein O1 (pFoxo1); and decreased phosphorylated AMP-activated protein kinase alpha (pAmpkα) and pFbp1 in *ob*/*ob* mice.

For the metabolome, we quantified the amounts of 183 polar metabolites using capillary electrophoresis mass spectrometry (CE-MS) (Figure 2E; Table S5). We identified polar metabolites that significantly changed in *ob*/*ob* mice compared with WT mice as differentially expressed metabolites (DEMs). We identified 24 (13%) increased and 100 (55%) decreased DEMs in *ob*/*ob* mice compared to WT mice (Figure 2E). The increased DEMs included ketone body such as 3-hydroxybutyrate. The decreased DEMs included UDP-glucose, cofactors ATP, phosphoenolpyruvate (PEP), and serine (Figures S2F and S2G).

For the lipidome, we quantified the amounts of 17 lipid classes using supercritical fluid chromatography tandem mass spectrometry (SFC-MS/MS) and liquid chromatography tandem mass spectrometry (LC-MS/MS) (Figure 2F; Table S6). We identified lipid classes that significantly changed in *ob*/*ob* mice compared with WT mice as differentially expressed lipids (DELs). We identified six increased and six decreased DELs in *ob*/*ob* mice compared to WT mice. Most energy storage lipids including diacylglycerol (DG), fatty acid, and triacylglycerol (TG) increased in *ob*/*ob* mice. Most membrane component lipids including lysophosphatidylethanolamine (LPE), phosphatidylethanolamine (PE), and phosphatidylserine (PS) decreased in *ob*/*ob* mice.

To reveal the regulatory mechanism of the expression of DEGs, we inferred the TFs that potentially regulate the DEGs using ChIP experimental data obtained from the ChIP-Atlas database (STAR Methods)^35^. We identified those included in DEGs or DPPs as differentially expressed TFs (DETFs) or differentially phosphorylated TFs (DPTFs). We integrated them as differentially regulated TFs (DRTFs). We identified one increased DRTF, peroxisome proliferator activated receptor gamma (*Pparg*), and 22 decreased DRTFs (Table S7). Some of the decreased DRTFs are related to obesity, such as nuclear receptor subfamily 3 group C member 1 (*Nr3c1*), CCAAT enhancer binding protein alpha (*Cebpa*), hepatocyte nuclear factor 4 alpha (*Hnf4a*), and *Foxa1*^40,41^ (Figure 2G). We inferred protein kinases that potentially phosphorylate DPPs by the kinase-substrate relationship through the amino acid sequences of phosphorylated peptides using the Netphorest database (STAR Methods)^36,37^. We identified those also included in DEGs or DPPs as differentially expressed kinases (DEKs) or differentially phosphorylated kinases (DPKs). In further, we defined them as differentially regulated kinases (DRKs). We identified 10 increased DRKs, and 8 decreased DRKs (Table S8). Among them, *Akt1*, protein kinase C zeta (*Prkcz*), and mitogen-activated protein kinase 3 (*Mapk3*) were key signaling molecules in the insulin signaling pathway (Figure 2H).

We identified the changed molecules in each omic layer in *ob*/*ob* mice, highlighted the metabolic pathways they involved in, and prepared nodes for constructing differential regulatory trans-omic network of metabolic reactions at *ad libitum* feeding.

### Identification of differential regulations in *ob*/*ob* mice at *ad libitum* feeding

We identified differential regulations that were connected by regulating (source) and regulated (target) differentially expressed or phosphorylated molecules between WT and *ob*/*ob* mice (Figure 3). Differential regulations from the DEGs to DEPs had the same changed directions (Figures 3A and S1D) (*e*.*g*., the increased DEGs to the increased DEPs, or the decreased DEGs to the decreased DEPs). The increased DEPs, 339 out of 676 (50%), corresponded to the increased DEGs. The decreased DEPs, 317 out of 582 (54%), corresponded to the decreased DEGs. These results suggest that half of the increased or decreased DEPs were regulated by gene expression. We identified differential regulations from proteins or phosphoproteins to metabolic reactions based on the reactions that DEPs or DPPs functioned as metabolic enzymes recorded by the KEGG database (Figures 3B and 3C). We found that 293 (43%) increased DEPs and 160 (27%) decreased DEPs were metabolic enzymes, 42 (30%) increased DPPs, 43 (34%) decreased DPPs, and 13 (28%) both increased and decreased DPPs were metabolic enzymes.

**Figure 3.**
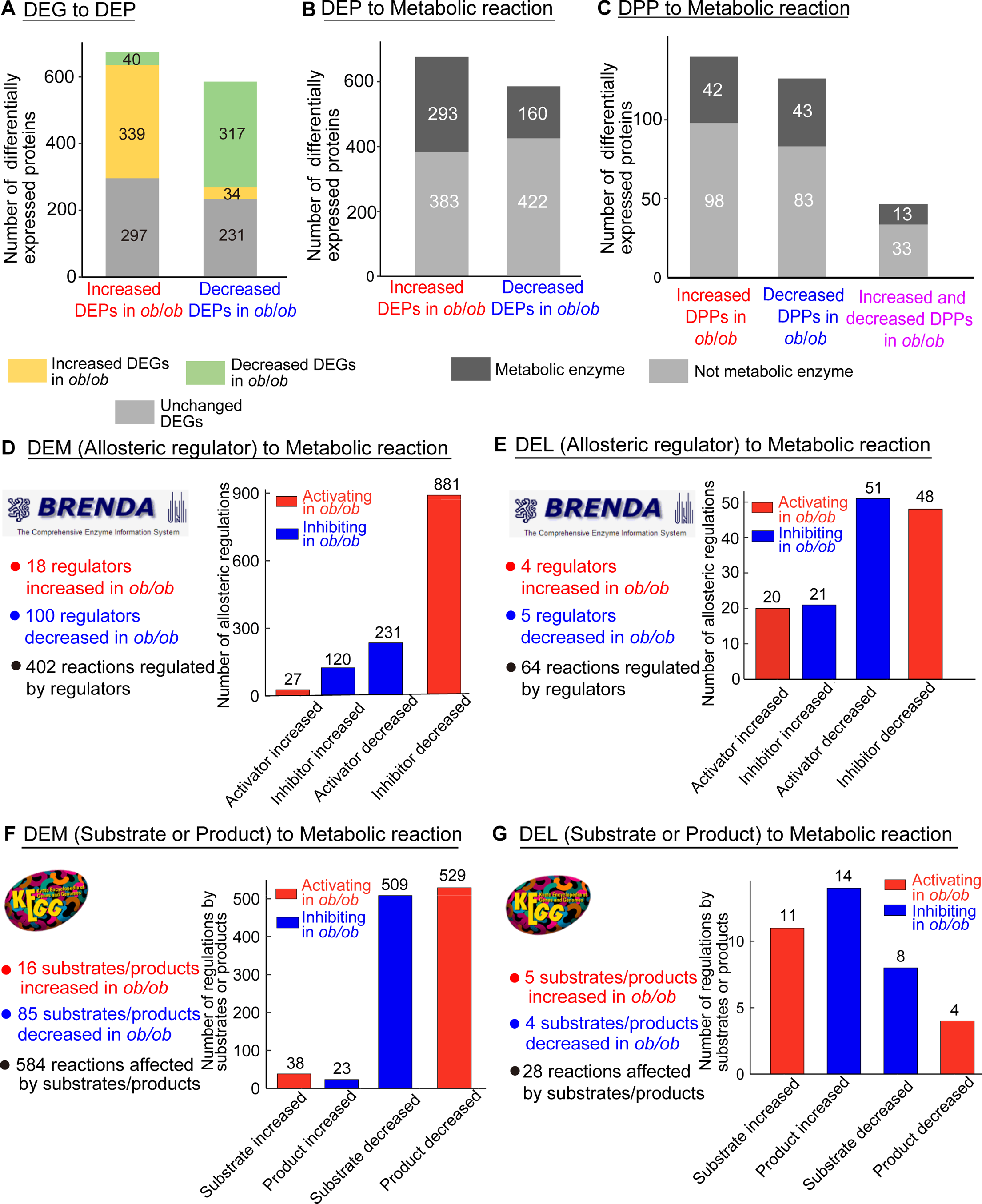
Identification of differential regulations in the liver between WT and *ob*/*ob* mice at *ad libitum* feeding. (A) The number of the increased and decreased DEPs encoded by the increased, decreased, or unchanged DEGs in *ob*/*ob* mice. The IDs of proteins and mRNAs were unified by Ensembl ID. (B) The number of metabolic enzymes and not metabolic enzymes among the increased and decreased DEPs in *ob*/*ob* mice. The IDs of proteins were unified by Entrez ID. (C) The number of metabolic enzymes and not metabolic enzymes among the increased, decreased, both increased and decreased DPPs in *ob*/*ob* mice. The IDs of phosphoproteins were unified by Entrez ID. (D) The number of the increased and decreased DEMs in *ob*/*ob* mice as allosteric regulators, and the number of regulated metabolic enzymes by allostery (left). Bar plot of activating and inhibiting regulations in *ob*/*ob* mice by allosteric regulators (right) using the BRENDA database. (E) The number of the increased and decreased DELs in *ob*/*ob* mice as allosteric regulators, and the number of regulated metabolic enzymes by allostery (left). Bar plot of activating and inhibiting regulations in *ob*/*ob* mice by allosteric regulators (right) using the BRENDA database. (F) The number of the increased and decreased DEMs in *ob*/*ob* mice as substrates or products, and the number of regulated metabolic enzymes by substrates or products (left). Bar plot of activating and inhibiting regulations in *ob*/*ob* mice affected by substrates or products (right) using the KEGG database. (G) The number of the increased and decreased DELs in *ob*/*ob* mice as substrates or products, and the number of regulated metabolic enzymes by substrates or products (left). Bar plot of activating and inhibiting regulations in *ob*/*ob* mice affected by substrates or products (right) using the KEGG database. When the regulating molecule increased in *ob*/*ob* mice was an activator, or that decreased in *ob*/*ob* mice was an inhibitor, then this regulation is activating regulation. When the regulating molecule decreased in *ob*/*ob* mice was an activator, or that increased in *ob*/*ob* mice was an inhibitor, then this regulation is inhibiting regulation. See also Figures S1D and S3A.

Differential regulation of metabolic reactions depends not only on amount and phosphorylation of metabolic enzymes but also on metabolites and lipids through allosteric regulations, or changes in the amount of substrates or products. We identified differential allosteric regulations of metabolic reaction by DEMs and DELs using the BRENDA database (Figures 3D and 3E; STAR Methods; Supplemental file). DEMs or DELs were classified as activators or inhibitors by their regulatory effects on metabolic enzymes in each reaction by using the BRENDA database^19^. Combined with the changed direction of activator or inhibitor in *ob*/*ob* mice, we had four conditions: activator increased led to activating regulation; activator decreased led to inhibiting regulation; inhibitor increased led to inhibiting regulation; inhibitor decreased led to activating regulation (Figure S1D). For allosteric regulations regulated by DEMs, 70% of allosteric regulations were activating allosteric regulations by decreased DEMs that functioned as inhibitors in *ob*/*ob* mice, including ATP, ADP, and GTP (Figures 3D and S3A). For allosteric regulations regulated by DELs, the numbers of activating and inhibiting regulations in *ob*/*ob* mice were almost the same (Figure 3E). PS, PE, and PI were the main lipid allosteric regulators that regulated metabolic reactions (Figure S3A). We identified metabolic reactions mediated by DEMs or DELs that functioned as substrates and products using the KEGG database (Figures 3F, 3G, and S3A; Supplemental file). For metabolic reactions affected by DEMs as substrates or products, 584 metabolic reactions were affected by 16 increased and 85 decreased substrates/products in *ob*/*ob* mice (Figure 3F). The numbers of activating and inhibiting regulations were almost the same (529 activating regulations and 509 inhibiting regulations). For metabolic reactions affected by DELs as substrates or products, 28 metabolic reactions were affected by 5 increased and 4 decreased substrates/products in *ob*/*ob* mice (Figure 3G).

We examined the number of DEPs and DPPs functioned as metabolic enzymes, and found predominant activating allosteric regulations induced by decreased inhibitors in *ob*/*ob* mice at *ad libitum* feeding. We prepared edges for constructing differential regulatory trans-omic network of metabolic reactions at *ad libitum* feeding.

### Construction of a differential regulatory trans-omic network of metabolic reactions at *ad libitum* feeding

We constructed a differential regulatory trans-omic network of metabolic reactions by integrating differentially expressed or phosphorylated molecules and differential regulations (Figure 4; STAR Methods). We selected DEGs, DEPs, and DPPs which functioned as metabolic enzymes, and defined them as Enzyme mRNA, Enzyme Protein, and Enzyme Phosphoprotein. The differential regulatory trans-omic network consisted of three components: nodes, edges, and layers (Figures 4A and S1B-D; Supplemental file). Nodes indicated differentially expressed or phosphorylated molecules. Edges indicated differential regulations. Layers indicated a group of nodes belonging to the same omic data or the same category. There is no regulation between nodes within the same layer, but there is regulation between nodes from different layers. From enzyme mRNA layer to enzyme protein layer, we only considered the molecules that show consistent changes, *i*.*e*., both increased in *ob*/*ob* mice, or both decreased in *ob*/*ob* mice. We observed generally activating regulations from Enzyme mRNA through Enzyme Protein to Metabolic Reaction, and inhibiting regulations from TF to Enzyme mRNA in *ob*/*ob* mice. In addition, metabolites as allosteric regulators induced more activating than inhibiting allosteric regulations. More increased Protein Kinases induced more activating than inhibiting regulations from Protein Kinase to Enzyme Phosphoprotein. In total, 527 Metabolic Reactions were defined as the opposite Metabolic Reactions (Table S9). The numbers of other activating and inhibiting regulations were almost equivalent. There was no regulation from insulin signaling molecules to TF and Protein Kinase. Decreased mRNAs in *ob*/*ob* mice are likely to be intensively regulated by TFs, while increased mRNAs are regulated by other mechanisms, such as DNA methylation.

**Figure 4.**
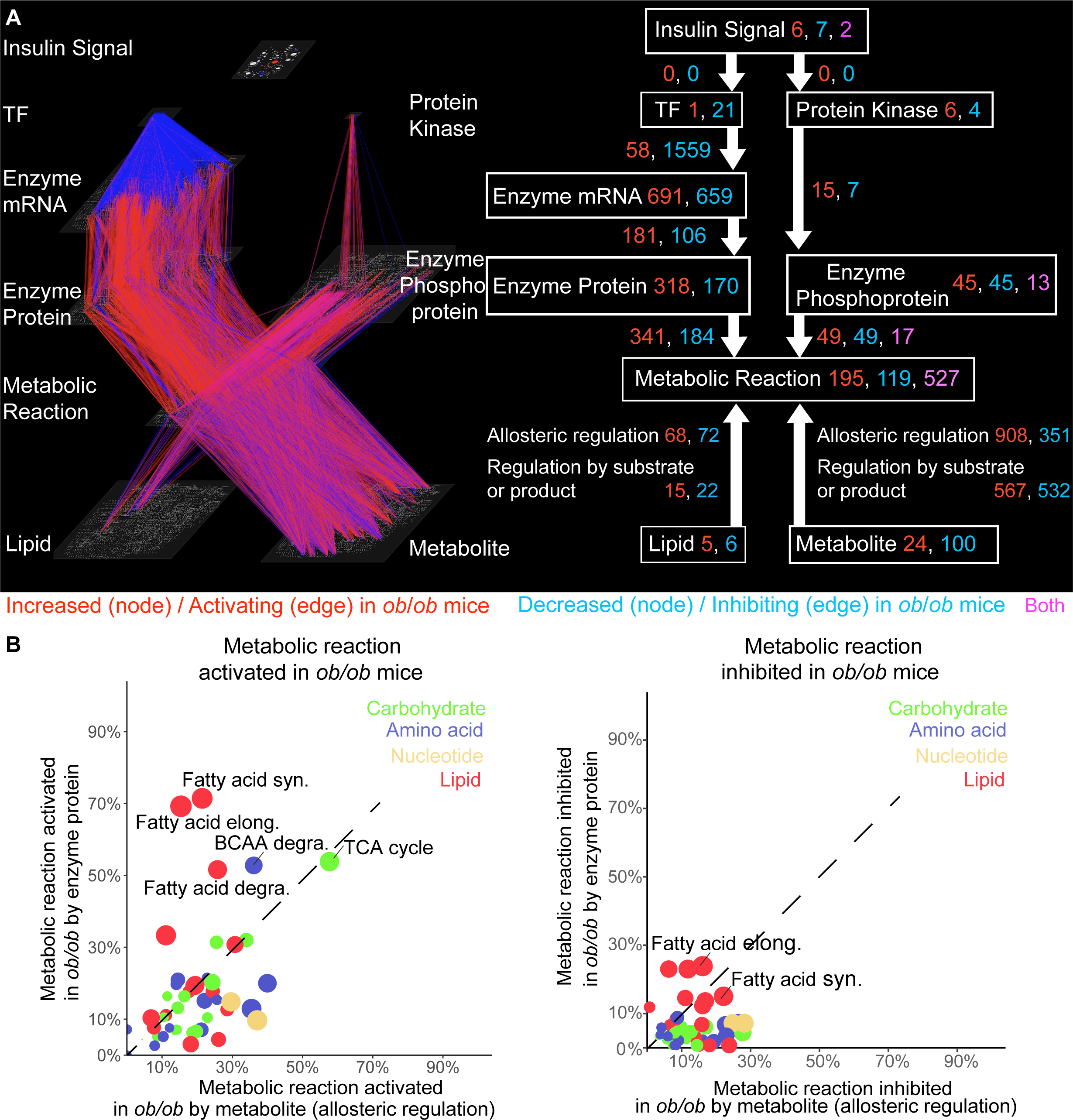
Construction of the differential regulatory trans-omic network of metabolic reactions in the liver between WT and *ob*/*ob* mice at *ad libitum* feeding. (A) The trans-omic network for the differentially expressed or phosphorylated molecules and differential regulations involved in metabolic pathways. Red, increased molecules or activating regulations in *ob*/*ob* mice; blue, decreased molecules or inhibiting regulations in *ob*/*ob* mice; magenta, both increased (activating) and decreased (inhibiting) molecules (regulations) in *ob*/*ob* mice. The detailed numbers of nodes and edges were shown on the right panel. Because some metabolic reactions were both activated and inhibited by different molecules, we defined such metabolic reactions as opposite metabolic reactions (magenta). See also Figure S1D. (B) For each metabolic pathway (circle), the percentage of differentially regulated metabolic reactions mediated by either metabolite as allosteric regulator (*x*-axis) or enzyme protein (*y*-axis). Left, activated metabolic reactions in *ob*/*ob* mice; right, inhibited metabolic reactions in *ob*/*ob* mice. The sizes of dots indicate the sum of percentages of differentially regulated metabolic reactions mediated by metabolites (allosteric regulators) and enzyme proteins. The colors of the dots indicate the classes of metabolism recorded by the KEGG database. See also the percentage of differentially regulated metabolic reactions mediated by metabolites as substrates and products, or enzyme proteins for each metabolic pathway in Figure S3B.

We further examined the regulations of Carbohydrates, Amino acids, Nucleotides, and Lipid metabolism using the KEGG database. We separately extracted the activated or inhibited metabolic reactions from each metabolic pathway. We calculated the percentage of dysregulated metabolic reactions in each metabolic pathway by using the number of dysregulated metabolic reactions divided by the number of total metabolic reactions (Figures 4B, S3B and S3C). Several Lipid metabolisms including fatty acid synthesis, elongation, and degradation, were regulated by enzyme proteins rather than by metabolites, whereas some Amino acids, Nucleotides, and Carbohydrates metabolism were mainly regulated by metabolites (Figures 4B, S3B, and S3C).

### Construction of activating and inhibiting condensed networks at *ad libitum* feeding

To highlight which metabolic pathways are activated or inhibited, we condensed the differential regulatory trans-omic network into the activating and inhibiting condensed networks according to previously reported procedures^33^ (Figure 5; STAR Methods). First, we grouped the metabolic reactions in each metabolic pathway of the differential regulatory trans-omic network into one single node, metabolic pathway node, by using the KEGG database. Through this step, we created a new layer, the Metabolic Pathway layer, instead of the Metabolic Reaction layer. We used the regulations from molecules to metabolic pathways to replace the regulations from molecules to metabolic reactions. Second, we removed less critical molecules and metabolic pathways by setting a threshold on the number of regulations as simplification. By extracting the activating and inhibiting regulations and related molecules, respectively, we further separated the condensed network into activating (Figure 5A) and inhibiting (Figure 5B) condensed networks. The activating and inhibiting condensed networks revealed an overview of the dysregulation of metabolic pathways in *ob*/*ob* mice (Supplemental file). There were more TF regulations to Enzyme mRNA in the inhibiting condensed network (286 regulations) than those in the activating condensed network (27 regulations). There were more Enzyme Protein regulations, Enzyme Phosphoprotein regulations, and allosteric regulations by Metabolites to Metabolic Pathway in the activating condensed network (363 Enzyme Protein regulations, 68 Enzyme Phosphoprotein regulations, and 1355 allosteric regulations by Metabolites) than those in the inhibiting condensed network (69 Enzyme Protein regulations, 43 Enzyme Phosphoprotein regulations, and 373 allosteric regulations by Metabolites). More Metabolic Pathways are activated (33 Metabolic Pathways) than inhibited (17 Metabolic Pathways) including Lipid, Amino acid, and Carbohydrate metabolism. ATP, ADP, NADH, NADPH, NAD^+^, and CoA functioned as allosteric regulators, causing large number of allosteric regulations, as we discussed in Figure 3D and Figure S3A. In further, we extracted molecules and regulations in specific metabolic pathways from differential regulatory trans-omic network at *ad libitum* feeding. We found the dysregulations of these metabolic pathways in the liver of *ob*/*ob* mice at *ad libitum* feeding (Figure S4).

**Figure 5.**
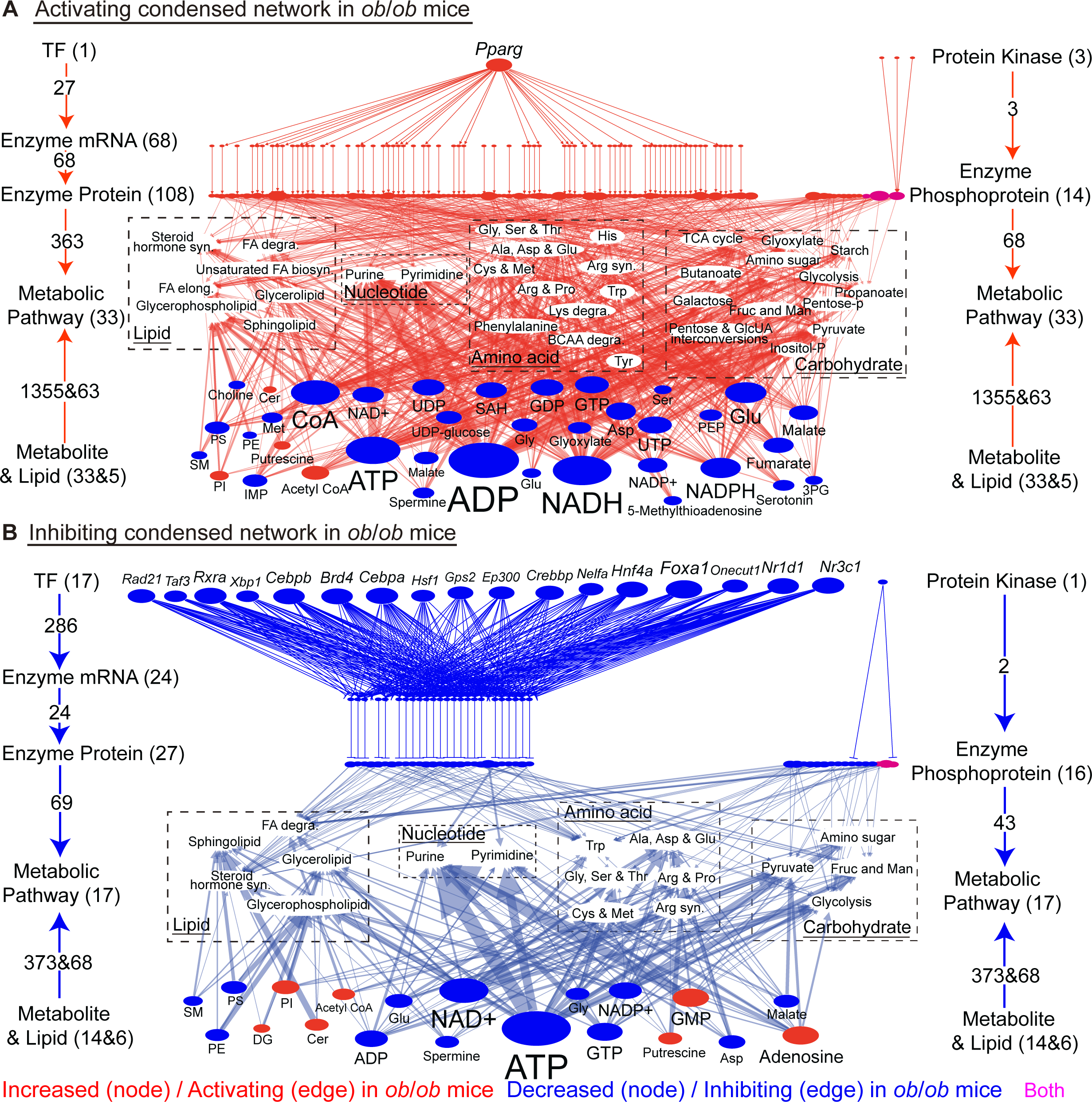
Construction of activating and inhibiting condensed networks in *ob*/*ob* mice at *ad libitum* feeding. (A) Activating condensed network containing activating regulations, differentially expressed or phosphorylated molecules connecting with activating regulations, and activated metabolic pathways in *ob*/*ob* mice. (B) Inhibiting condensed network containing inhibiting regulations, differentially expressed or phosphorylated molecules connecting with inhibiting regulations, and inhibited metabolic pathways in *ob*/*ob* mice. Metabolic pathways were shown in white ovals. Dashed boxes enclose metabolic pathways belonging to Lipid, Nucleotide, Amino acid, and Carbohydrate metabolism. The size of node and width of edge indicate the number of differentially expressed or phosphorylated molecules and differential regulations. Red, increased molecules or activating regulations in *ob*/*ob* mice; blue, decreased molecules or inhibiting regulations in *ob*/*ob* mice; magenta, the DPPs contain both increased and decreased phosphorylation sites in *ob*/*ob* mice.

Condensed network highlighted the conclusions drawn in trans-omic network: TF regulations generally show inhibiting, while enzyme protein, enzyme phosphoprotein, allosteric regulations generally show activating in *ob*/*ob* mice compared to WT mice at *ad libitum* feeding. Activating enzyme protein regulations may not be regulated by TFs in *ob*/*ob* mice, but regulated by other mechanisms, such as DNA methylation. Most activating allosteric regulations are caused by decreased metabolites in *ob*/*ob* mice, indicating a flexible way of modulation.

### Comparison of the trans-omic networks between *ad libitum* feeding and 16 h-fasting

To reveal the difference in metabolic dysfunctions in the liver of *ob*/*ob* mice between *ad libitum* feeding and 16 h-fasting, we compared the differential regulatory trans-omic network of metabolic reactions at *ad libitum* feeding in this study with that at 16 h-fasting in our previous study^34^ (Figures 6, 7, S5 and S6).

**Figure 6.**
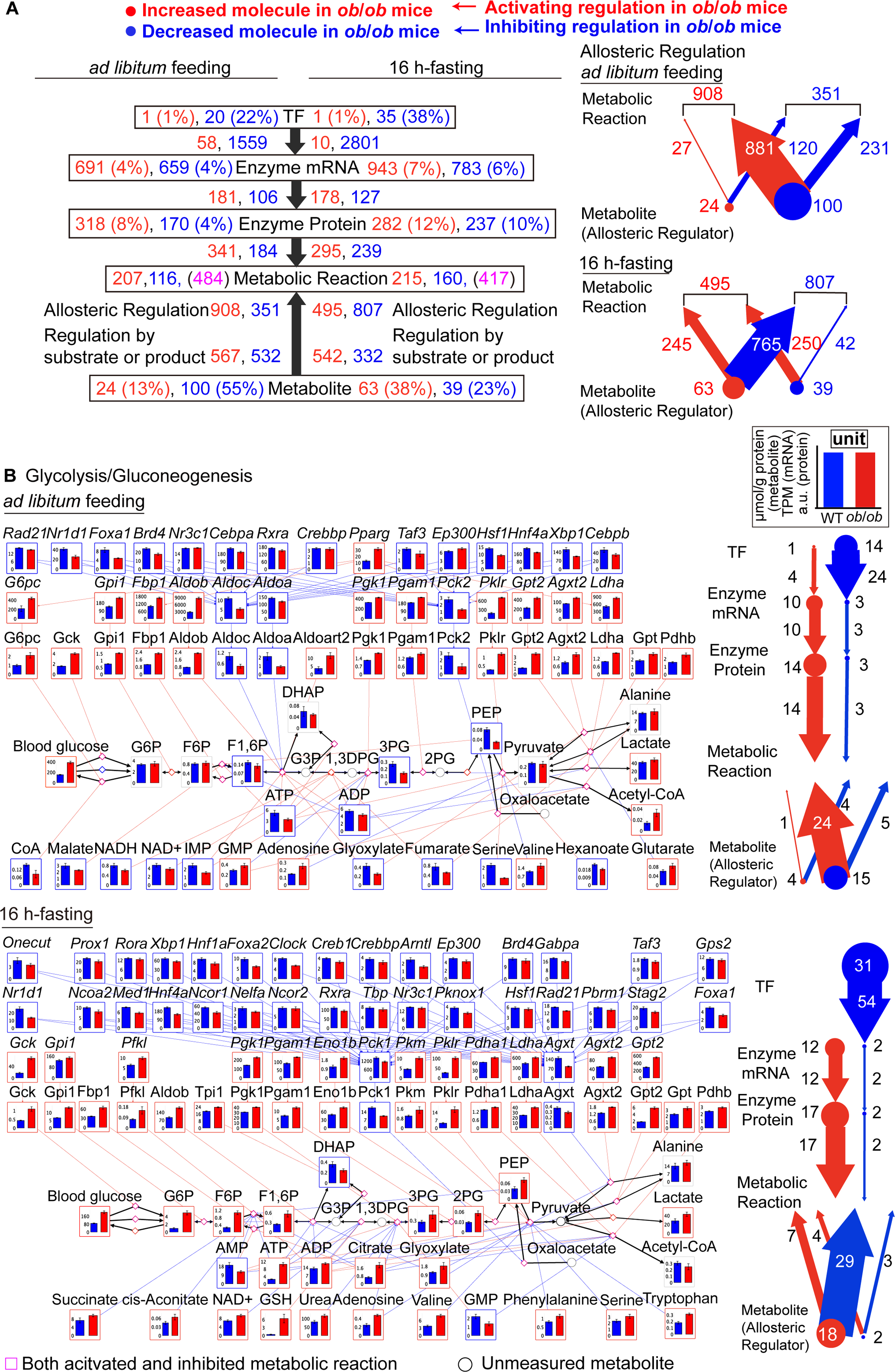
Comparison of the differential regulatory trans-omic network and glycolysis/gluconeogenesis pathway between *ad libitum* feeding and 16 h-fasting. (A) Differential regulatory trans-omic network including differentially expressed molecules and differential regulations at *ad libitum* feeding and 16 h-fasting (left)^34^. The percentages are calculated by dividing the total number of measured molecules or background molecules. The numbers of activating and inhibiting Allosteric Regulation (edge) regulated by Metabolite (Allosteric Regulator) (node) in *ob*/*ob* mice (right). The size of node and width of edge indicate the number of Metabolite (Allosteric Regulator) and Allosteric Regulation, we show the numbers around nodes and edges. See also Figures S5 and S6. (B) Metabolic reactions, molecules, and regulations in Glycolysis/Gluconeogenesis pathway (mmu00010) extracted from the differential regulatory trans-omic network at *ad libitum* feeding (top-left) and 16 h-fasting (bottom-left)^34^. Bar plots are shown on the corresponding nodes as the means and standard error of the mean of replicates in WT and *ob*/*ob* mice. For metabolites as substrates or products in Glycolysis/Gluconeogenesis pathway, we showed bar plots for all measured molecules. The colors of frame and edge indicate the changed direction. Diamond nodes indicate metabolic reactions. Black edges indicate the metabolic reactions between substrates and products. Black circles indicate the metabolites that were not measured. From metabolite to metabolic reaction, only allosteric regulations are colored. We counted the numbers of differentially expressed TF, Enzyme mRNA, Enzyme Protein, Metabolite, Metabolite (Allosteric Regulator) and differential regulations in Glycolysis/Gluconeogenesis pathway at *ad libitum* feeding (top-right) and 16 h-fasting (bottom-right). We used the size of node and the width of edge to indicate them. For those with more than 30 or less than 2, we used one size respectively (>30, 30, <2, 2) in case too big or too small nodes and edges. Specific numbers were shown around nodes and edges. Note that the plotting scales are different from Figure 6A (right) and Figure 6B (right). See Figure S7 for other metabolic pathways. Red, increased molecules or activating regulations in *ob*/*ob* mice; blue, decreased molecules or inhibiting regulations in *ob*/*ob* mice; gray, unchanged molecules between WT and *ob*/*ob* mice; magenta, metabolic reaction is both activated and inhibited in *ob*/*ob* mice.

**Figure 7.**
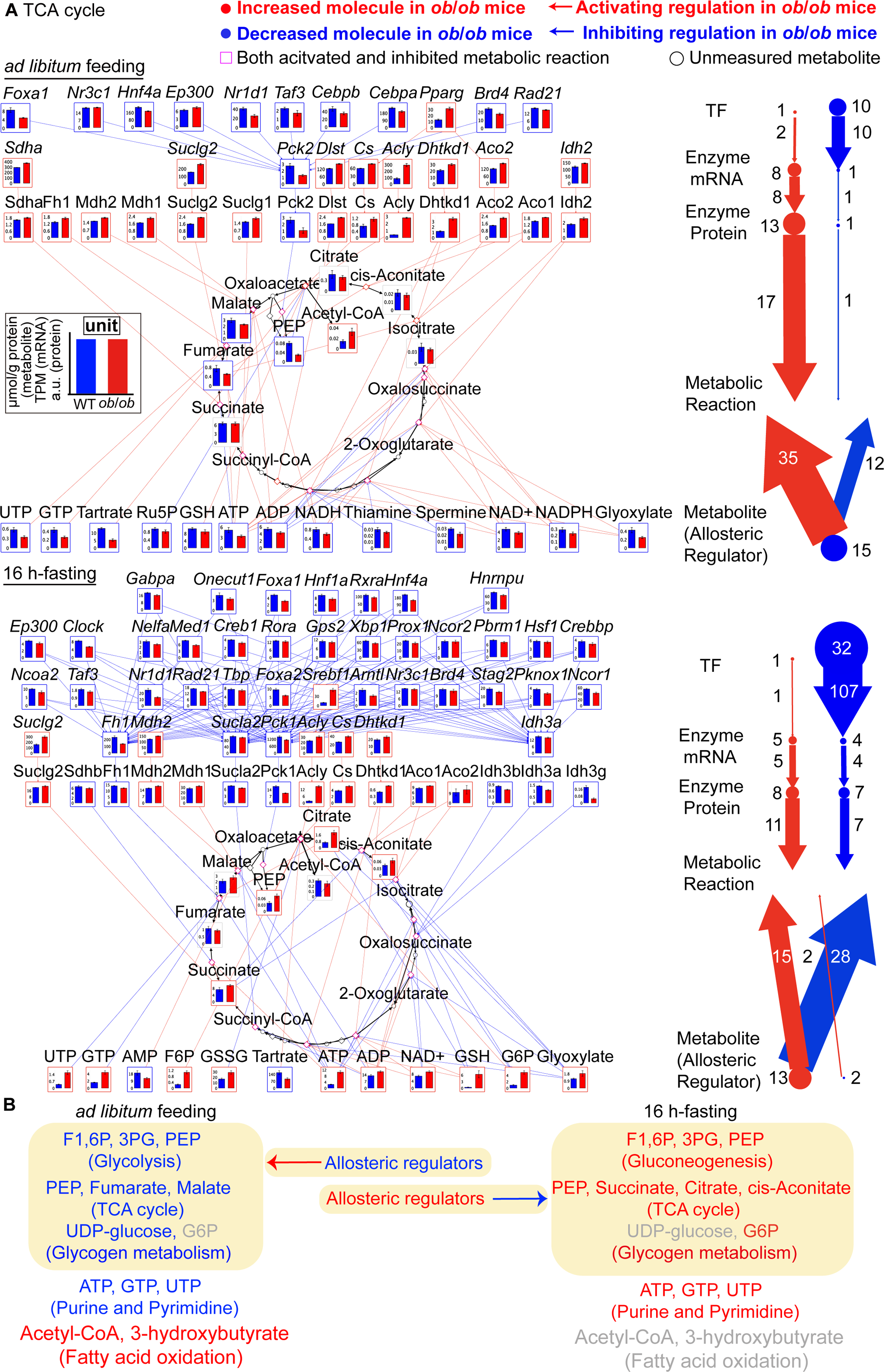
Comparison of TCA cycle between *ad libitum* feeding and 16 h-fasting. (A) Metabolic reactions, molecules, and regulations in TCA cycle (mmu00020) extracted from the differential regulatory trans-omic network at *ad libitum* feeding (top left) and 16 h-fasting (bottom left), and the numbers of differentially expressed molecules (nodes) and differential regulations (edges) in TCA cycle at *ad libitum* feeding (top right) and 16 h-fasting (bottom right)^34^. Other descriptions are the same as Figure 6B. (B) Summary of the opposite changes in the metabolites and allosteric regulations between *ad libitum* feeding (left) and 16 h-fasting (right)^34^. Metabolic pathway names related to metabolites are in parenthesis. Arrow, allosteric regulation. Yellow area, the metabolic pathways regulated by allosteric regulators. See also Figure S7. Red, increased molecules or activating regulations in *ob*/*ob* mice; blue, decreased molecules or inhibiting regulations in *ob*/*ob* mice; magenta, both activated and inhibited metabolic reactions in *ob*/*ob* mice; gray, unchanged molecules between WT and *ob*/*ob* mice.

The differential regulatory trans-omic networks at *ad libitum* feeding in our present study and at 16 h-fasting in our previous study were slightly different in methods of TF, metabolic enzyme, and allosteric regulation inference. For direct comparisons, we modified the trans-omic network at 16 h-fasting by renewing TF inference using the ChIP-Atlas database, and by renewing metabolic enzyme and allosteric regulation inference using the same version of BRENDA and KEGG database in this study (see STAR Methods). The main conclusions were the same in trans-omic network at 16 h-fasting before and after modification, except TF regulations, which generally show activating inferred by the TRANSFAC database^34^ while generally show inhibiting inferred by the ChIP-Atlas database (Figure 6A). We also removed the Protein Kinase layer, Enzyme Phosphoprotein layer, and Lipid layer from the trans-omic network at *ad libitum* feeding, which were not present in trans-omic network at 16 h-fasting^34^. In addition, we removed the Insulin Signal layer from both trans-omic networks for which measured by western blot and phosphoproteome at *ad libitum* feeding, while measured by only western blot at 16 h-fasting.

Overall, we compared TF, Enzyme mRNA, Enzyme Protein, and Metabolite layers in trans-omic networks between *ad libitum* feeding and 16 h-fasting (Figure 6A). In further, we compared the metabolic pathways extracted from trans-omic networks, including glycolysis/gluconeogenesis, TCA cycle, purine and pyrimidine metabolism, glycogen metabolism, fatty acid synthesis and degradation, arginine synthesis between *ad libitum* feeding and 16 h-fasting (Figures 6B, 7A, and S7). We counted the numbers of differentially expressed TFs, Enzyme mRNAs, Enzyme Proteins, Metabolites (Allosteric Regulators), and differential regulations between *ad libitum* feeding and 16 h-fasting and show them on the right panel (Figures 6B, 7A, and S7; Supplemental file). We drew three main conclusions as follows.

### Intermediate metabolites in glycolytic metabolism and triphosphates in nucleotide metabolism decreased in *ob*/*ob* mice at *ad libitum* feeding but increased at 16 h-fasting

In glycolysis/gluconeogenesis pathway, the intermediate metabolites fructose 1,6-bisphosphate (F1,6P), 3-phosphoglyceric acid (3PG), and PEP decreased in *ob*/*ob* mice at *ad libitum* feeding, whereas they increased in *ob*/*ob* mice at 16 h-fasting (Figure 6B). In the TCA cycle, the intermediate metabolites, PEP, fumarate, and malate, decreased in *ob*/*ob* mice only at *ad libitum* feeding. By contrast, citrate, cis-aconitate, and succinate increased only at 16 h-fasting (Figure 7A). In purine metabolism, ATP, ADP, and GTP, which are the important energy-providing nucleotides, cofactors, and allosteric regulators, decreased in *ob*/*ob* mice at *ad libitum* feeding but increased at 16 h-fasting (Figure S7A). In pyrimidine metabolism, UDP and UTP decreased at *ad libitum* feeding while increased at 16 h-fasting (Figure S7B). In glycogen metabolism, glycogen increased in *ob*/*ob* mice at both *ad libitum* feeding and 16 h-fasting (Figure S7C). However, UDP-glucose, the substrate of glycogen, decreased at *ad libitum* feeding and did not change at 16 h-fasting.

### Allosteric regulation reversely shifted between feeding and fasting

For allosteric regulations, we observed generally activating allosteric regulations at *ad libitum* feeding, while generally inhibiting allosteric regulations at 16 h-fasting in *ob*/*ob* mice (Figure 6A). Among them, 100 decreased metabolites caused 881 activating allosteric regulations at *ad libitum* feeding, whereas 63 increased metabolites caused 765 inhibiting allosteric regulations at 16 h-fasting. In glycolysis/gluconeogenesis pathway, 15 decreased allosteric regulators caused 24 activating allosteric regulations in *ob*/*ob* mice at *ad libitum* feeding, whereas 18 increased allosteric regulators caused 29 inhibiting allosteric regulations at 16 h-fasting (Figure 6B). In TCA cycle, all 15 decreased allosteric regulators caused 35 activating allosteric regulations and 12 inhibiting allosteric regulations at *ad libitum* feeding, whereas 13 increased allosteric regulators caused 15 activating allosteric regulations and 28 inhibiting allosteric regulations at 16 h-fasting (Figure 7A). Similarly, in glycogen metabolism (Figure S7C) and arginine synthesis (Figure S7F), we observed generally activating allosteric regulations at *ad libitum* feeding while generally inhibiting allosteric regulations at 16 h-fasting in *ob*/*ob* mice.

We observed predominant activating allosteric regulations caused by decreased allosteric regulators in *ob*/*ob* mice at *ad libitum* feeding, while predominant inhibiting allosteric regulations caused by increased allosteric regulators in *ob*/*ob* mice at 16 h-fasting. The opposite allosteric regulation pattern in glycolysis/gluconeogenesis, TCA cycle, glycogen metabolism, and arginine synthesis was same as the overall allosteric regulation pattern (Figure 6A).

### Transcriptional regulation does not shift between *ad libitum* feeding and 16 h-fasting

Transcriptional regulation includes the regulation from TF to Enzyme mRNA to Enzyme Protein to Metabolic Reaction. The number of inhibiting TF regulations, activating Enzyme mRNA regulations, activating Enzyme Protein regulations in *ob*/*ob* mice were more than WT mice at both *ad libitum* feeding and 16 h-fasting trans-omic networks (Figure 6A), indicating that there is not clearly shift on transcriptional regulation between *ad libitum* feeding and 16 h-fasting, unlike the allosteric regulation. Similarly, in metabolic pathways, including glycolysis/gluconeogenesis (Figure 6B), TCA cycle (Figure 7A), pyrimidine metabolism (Figure S7B), glycogen metabolism (Figure S7C), fatty acid synthesis (Figure S7D) and degradation (Figure S7E), we did not observe a clear shift on transcriptional regulation between *ad libitum* feeding and 16 h-fasting.

In conclusion, we found the opposite metabolic dysregulation between *ad libitum* feeding and 16 h-fasting associated with obesity, which were shown on intermediate metabolites in carbohydrate and nucleotide metabolism, and allosteric regulations (Figure 7B). At *ad libitum* feeding, we observed the decreased intermediate metabolites in glycolysis and the TCA cycle (Figures 6B and 7A); decreased UDP-glucose in glycogen synthesis (Figure S7C), and decreased nucleotides in purine and pyrimidine metabolism (Figures S7A and S7B). Acetyl-CoA and 3-hydroxybutyrate in fatty acid oxidation increased (Figure S5E) in *ob*/*ob* mice. At 16 h-fasting, we observed the increased intermediate metabolites in the gluconeogenesis pathway and TCA cycle (Figures 6B and 7A), and increased nucleotides in purine and pyrimidine metabolisms (Figure S7A and S7B). Acetyl-CoA and 3-hydroxybutyrate in fatty acid oxidation did not change between WT and *ob*/*ob* mice (Figure S5E). Decreased allosteric regulators in *ob*/*ob* mice led to activating allosteric regulations at *ad libitum* feeding. By contrast, increased allosteric regulators in *ob*/*ob* mice led to inhibiting allosteric regulations at 16 h-fasting. The opposite metabolic dysregulation characterized the breakdown of liver metabolic homeostasis associated with obesity. We did not observe clear shift of transcriptional regulations between *ad libitum* feeding and 16 h-fasting. Some mathematical models were needed to determine the key contributor among enzyme proteins to answer this question^42^.

## DISCUSSION

In this study, we measured multi-omics data in the liver of WT and *ob*/*ob* mice at *ad libitum* feeding, and constructed the differential regulatory trans-omic network of metabolic reactions. We compared the trans-omic network between *ad libitum* feeding with that of 16 h-fasting^34^. Our study provided new insights into the differential shifts and common patterns in liver metabolism in response to feeding and fasting in *ob*/*ob* mice. We found the opposite metabolic dysregulation between *ad libitum* feeding and 16 h-fasting associated with obesity. Intermediate metabolites in carbohydrate (glycolysis/gluconeogenesis, TCA cycle, glycogen metabolism) and nucleotide pathways (purine and pyrimidine metabolism) decreased at *ad libitum* feeding but increased at 16 h-fasting (Figure 7B). In most metabolic pathways, allosteric regulations activating in *ob*/*ob* mice at *ad libitum* feeding while inhibiting in *ob*/*ob* mice at 16 h-fasting. Transcriptional regulation (from TF to Enzyme Protein to Metabolic Reaction) did not clearly shift between two trans-omic networks.

### The opposite dysregulation on energy-related metabolism

The liver has great flexibility in selecting metabolic fuels, glucose, or fatty acid, depending on feeding or fasting. Glucose oxidation is the process that breaks down of glucose to provide energy at feeding, including glycolysis and the TCA cycle. Fatty acid oxidation is the process of breaking down of fatty acid into acetyl-CoA, which predominantly provides energy at fasting. Meanwhile, liver generates glucose by gluconeogenesis at fasting to maintain glucose concentration^43^. The fold change of acetyl-CoA and intermediate metabolites in glucose oxidation could reflect whether glucose oxidation or fatty acid oxidation is dominant in living cells, given that glucose oxidation generates both while fatty acid oxidation generates only acetyl-CoA. In this study, at *ad libitum* feeding, glucose oxidation potentially decreased with decreased intermediate metabolites, but fatty acid oxidation potentially increased with increased acetyl-CoA and 3-hydroxybutyrate, indicating the shift from fatty acid oxidation to glucose oxidation is not sufficient even under enough food supply in *ob*/*ob* mice (Figure 7B). The decreased intermediate metabolites in glycolysis with increased blood glucose concentration may indicate insulin resistance in *ob*/*ob* mice at *ad libitum* feeding. At fasting, gluconeogenesis and TCA cycle potentially increased in *ob*/*ob* mice with increased intermediate metabolites, and without changes in acetyl-CoA and 3-hydroxybutyrate, suggesting that glucose oxidation is the major energy source compared with fatty acid oxidation even under starvation in *ob*/*ob* mice. These phenomena reveal the dysfunction of energy supply in *ob*/*ob* mice. ATP, ADP, GTP, UDP, and UTP, these important energy-providing molecules, cofactors, ribonucleotides, and allosteric regulators decreased at *ad libitum* feeding (high-energy state) but increased at 16 h-fasting (low-energy state) (Figures 7B, S7A, and S7B). These phenomena reveal the dysfunction of energy storage in *ob*/*ob* mice. In conclusion, the liver in *ob*/*ob* mice showed the opposite dysregulation of energy-related metabolism between feeding and fasting.

### Allosteric dysregulation can be the key pathological mechanism associated with obesity

Allosteric regulation of enzyme proteins is an adaptive mechanism to control metabolite concentration through feedback from metabolite to enzyme protein activity. Many conformational states of allosteric proteins exist with different enzymatic activities in the living individual^6^. Allosteric activation of glycogen synthase by allosteric activator G6P is the primary mechanism in the liver^44^ and muscle^45^ instead of phosphorylation of glycogen synthase. Similar to our previous study of 16 h-fasting^33^, some nucleotides, including ATP, ADP, AMP, and NAD, were widely involved in allosteric regulations at *ad libitum* feeding. In this study, we did not observe a clear shift on transcriptional regulation in most metabolic pathways, while we observed activating allosteric regulations in *ob*/*ob* mice at *ad libitum* feeding and inhibiting allosteric regulations in *ob*/*ob* mice at 16 h-fasting. Our previous study has reported the impairment of allosteric regulations in *ob*/*ob* mice during oral glucose administration^33^. Although the reason for this phenomenon remains unclear, one possible explain would be that transcriptional regulation replies on gene expression, which is slow and not easy to shift in response to feeding and fasting. Allosteric regulation by metabolites provides rapid modulation of enzyme activities by directly altering enzyme structures. Therefore, it is reasonable that allosteric regulation is widely used in response to feeding and fasting. These results reveal that compare to transcriptional regulation; the dysregulation of allosteric regulation may be one of the important pathological mechanisms associated with obesity.

### A common scenario on dysregulation of metabolism associated with obesity

Taken together with the differential regulatory trans-omic network constructed at 16 h-fasting^34^, we summarized a common scenario on dysregulation of metabolism associated with obesity in liver. First, reverse dysregulation of allosteric regulation. Second, generally decreased TFs with inhibiting TF regulations in *ob*/*ob* mice. Generally increased enzyme proteins in *ob*/*ob* mice without increased TF regulated. This may indicate that decreased enzyme proteins were regulated by decreased TFs in *ob*/*ob* mice, while increased enzyme proteins may not directly regulated by TFs but by other mechanisms, like DNA methylation. Third, generally activating transcriptional regulation in *ob*/*ob* mice (Figure 6A), which may indicate that metabolic activity is activated by enzyme proteins. Fourth, the generation of ribosome and spliceosome was inhibited in *ob*/*ob* mice (Figures S2A, S5C, and S5D). It is likely to reveal that protein synthesis was suppressed, and protein diversity was reduced (an energy-consuming process) in *ob*/*ob* mice. Some studies have shown that ribosomes are downregulated in the liver and muscle of *ob*/*ob* mice or high-fat diet mice due to essential amino acid deficiency^46–49^ or acute nutrient deprivation^50^, endoplasmic reticulum stress or ribosomal stress^51^. At 16 h-fasting, more differential molecules (DETFs, DEGs, DEPs) and differential enzyme protein regulations were observed than at *ad libitum* feeding (Figure 6A). However, at *ad libitum* feeding, there are more dysregulated metabolic pathways than 16 h-fasting (Figure S6C). These phenomena indicate that fasting could reduce metabolic dysregulation.

### Underlying mechanism of dysregulation of metabolic homeostasis associated with obesity at *ad libitum* feeding

Many multi-omics studies have revealed how obesity changes metabolism and related regulatory mechanism in living cells, organs, and individuals^17,18,33,52,53^. This study has the consistent findings with previous studies that the abundance of proteins related to fatty acid metabolism and peroxisome increased in *ob*/*ob* mice liver after fasting (Tables S1 and S2)^17^, increased mitochondrial fatty acid *β*-oxidation and ketone body formation, increased TG, decreased glycerophospholipids, decreased CE and acyl-Carnitines^18^, increased mRNAs *Pfkl* involved in carbohydrate metabolism in high-fat fed mice liver^53^ (Figures 2A, 2F, and S2H). Kim et al. observed significant downregulations of protein phosphorylation in high-fat fed mice liver, which is not observed in this study^18^.

We observed a significant increase of expression amount of key enzyme or rate-limiting enzyme^54^ in *ob*/*ob* mice in carbohydrate and lipid metabolic pathway, including glycolysis and gluconeogenesis (Gck, Pklr, Fbp1, Pcx, G6pc), TCA cycle (Cs, Aco1, Aco2, Suclg1, Sdha, Fh1, Mdh2, Dhtkd1, Suclg2, Mdh1), glycogen metabolism (Pygl, Agl), fatty acid synthesis (Fasn, Acacb, Acaca), fatty acid degradation (Acsl5, Acads, Ehhadh, Hsd17b10, Hadhb, Acaa1b, Echs1, Hadha, Hadh). We also observed a significant decrease of key enzyme in nucleotide metabolic pathway, including purine metabolism (Gart, Paics, Impdh2, Gmps), pyrimidine metabolism (Cad). Downstream of glycolysis, from PEP to pyruvate, can be activated by increased Pklr (Figure S4A). Upstream of gluconeogenesis, from Oxaloacetate to PEP, can be inhibited by decreased Pck2 and increased pPck1. Downstream of gluconeogenesis, from F1,6P to F6P, can be activated by increased Fbp1 and decreased pFbp1. These dysregulations accelerate the usage and retard the generation of F1,6P, 3PG, and PEP at *ad libitum* feeding. Many of decreased TFs has been reported to be associated with the onset of metabolic syndrome. CCAAT/enhancer-binding protein beta (*Cebpb*) is the principal TF involved in the early steps of adipogenesis^55^. *Hnf4a* mutants were demonstrated to modestly induce the onset of type 2 diabetes mellitus^56^.

In conclusion, our differential regulatory trans-omic network of metabolic reactions integrates multi-omics datasets and bioinformatics databases. It would hopefully provide an effective methodology for research on metabolism diseases beyond obesity. Meanwhile, the comparison of trans-omic networks between *ad libitum* feeding and 16 h-fasting provides key insights into the breakdown of metabolic homeostasis, the opposite metabolic dysregulation. The allosteric dysregulation maybe the key pathological mechanism associated with obesity.

### Limitations of the study

The study founds opposite allosteric regulation in WT and *ob*/*ob* mice between feeding and fasting. Further experimental validation of conformational changes in key enzymes caused by allosteric regulators during feeding and fasting would provide additional support for our conclusion. The opposite metabolic dysregulation observed in this study may not only directly reflect the breakdown of liver metabolic homeostasis but also the dysregulation of inter-organ metabolic cycles as the metabolites synthesized by liver circulate among different organs, including skeletal muscle, adipose tissue, and brain^57^. Systematical investigation of this issue based on the trans-omic networks of inter-organ metabolic cycles is needed in the future^34^. The trans-omic network in this study cannot determine the weight of different regulators and regulations. We will examine the effective contributions of regulators, including the amount of enzyme, phosphorylation, and allosteric regulation on changes in metabolites using mathematical modeling in the future^42^. This study revealed the opposite dysregulation of energy-related metabolism in *ob*/*ob* mice. The calculation of energy charge would be helpful for quantitatively understanding the energy-related dysregulation in *ob*/*ob* mice^58^. The underlying mechanism of metabolic dysregulation between feeding and fasting in liver associated with obesity remains to be further explored by adding epigenetic controls, including DNA methylation, histone modification, non-coding RNAs into trans-omic network. The amounts of food intake may also be different from WT and *ob*/*ob* mice, it would be more reasonable if we take it as one of variables.

## ACKNOWLEDGEMENTS

We thank Maki Ohishi and Ayano Ueno (Keio University) for their expertise and assistance with metabolome analysis using CE-MS. We thank Maiko Goto (Kyushu University) for technical assistance with lipidome analysis using SFC-MS/MS and LC-MS/MS. We thank Shinsuke Uda (Kyushu University) for giving us suggestions on statistical analysis. We also thank our laboratory members for critically reading this manuscript and technical assistance with the experiments. The computational analysis of this work was performed in part with support of the supercomputer system of National Institute of Genetics (NIG), Research Organization of Information and Systems (ROIS). The proteomic and phosphoproteomic measurements were performed in Kazusa DNA Research Institute.

This study was supported by the Japan Society for the Promotion of Science (JSPS) KAKENHI Grant Numbers JP17H06300, JP17H06299, JP18H03979, JP21H04759, CREST, the Japan Science and Technology Agency (JST) (JPMJCR2123), and by The Uehara Memorial Foundation. Y.B. receives funding from China Scholarship Council Grant Number 202008050067. K.M. receives funding from a Grant-in-Aid for Early-Career Scientists (JP21K15342). T.K. receives funding from JSPS KAKENHI (JP21K16349). S.O. receives funding from a Grant-in-Aid for Early-Career Scientists (JP21K14467). K.Y. receives funding from JSPS KAKENHI Grant Number JP18H05431, and CREST, JST (JPMJCR22N5). S.U. receives funding from JSPS KAKENHI Grant Number JP19J22134. Y.Inaba receives funding from the Japan Agency for Medical Research and Development (AMED) through the Practical Research Project for Life-Style related Diseases including Cardiovascular Diseases and Diabetes Mellitus Grant Number JP21ek0210156. Y.S. receives funding from JSPS KAKENHI Grant Number JP17H06306. M.T. receives funding from JSPS KAKENHI Grant Number JP20K15101. Y.Izumi receives funding from JSPS KAKENHI Grant Number JP22H01883. T.B. receives funding from JSPS KAKENHI Grant Number JP17H06304 and JP17H06299. T.S. receives funding from AMED under Grant Number JP21zf0127001.

## AUTHOR CONTRIBUTIONS

Y.B. and S.K. conceived and supervised the project; K.M. and A. Hatano designed and performed the animal experiments, sample preparation of omics measurements, enzymatic assays, and western blot measurements; A. Hirayama and T.S. performed metabolomic analysis using CE-MS; Y.S. performed transcriptomic analysis using RNA-seq; M.T., Y.Izumi, and T.B. performed lipidomic analysis using SFC-MS/MS and LC-MS/MS; A. Hatano and M.M. performed sample preparation of proteomic and phosphoproteomic analysis; Y.B., T.K., S.O., R.E., Y.P., D.L., K.Y., and S.U. performed trans-omic analysis; writing group consisted of Y.B., S.O., H.I, Y.Inaba, and S.K. All authors read and approved the final manuscript.

## STAR METHODS

### RESOURCE AVAILABILITY

#### Lead Contact

For any additional information and requests regarding resources and reagents, please contact the lead contact, Shinya Kuroda.

#### Materials Availability

The study did not generate any new material.

#### Data and Code Availability

- RNA-seq data measured in this study has been deposited in the DNA DataBank of Japan Sequence Read Archive (DRA) (www.ddbj.nig.ac.jp/) with accession number DRA015836. All other data generated in this study are uploaded into Tables S1–S9.
- The code used for the analysis in this paper is available online at Zenodo
- Any additional information required to reanalyze the data reported in this paper is available from the lead contact upon request.

## EXPERIMENTAL MODEL AND SUBJECT DETAILS

### Mouse studies

All procedures that involved animal experiments were approved by the University of Tokyo Animal Ethics Committee (Tokyo, Japan). We purchased 10-week-old male C57BL/6 WT and *ob*/*ob* mice from Japan SLC Inc (Hamamatsu, Japan). Just after the arrival, mice were allowed free access to water and standard rodent chow. Mice were euthanized by cervical dislocation (clock time: 16:00-20:00, GMT +9) following to the measurement of glucose levels. Livers were removed and frozen in liquid nitrogen, in further pulverized with dry ice to a fine powder with a Wonder Blender (WB-1, OSAKA CHEMICAL CO., LTD) or ShakeMaster NEO ver1.0 (BMS-M10N21, BMS) and separated into tubes for omic analysis (transcriptomics, proteomics, phosphoproteomics, metabolomics, and lipidomics), glycogen assay, and Western Blot analysis. Five replicates were used for WT and *ob*/*ob* mice. We used the same samples of liver in all experiments except Western Blot analysis.

## METHOD DETAILS

### Transcriptomics analysis

Total RNA extraction from liver samples collected at *ad libitum* feeding was performed by using the RNeasy Mini Kit (QIAGEN, Germantown, MD, USA) and QIAshredder (QIAGEN) according to the manufacturer’s protocol. The extracted RNA was assessed for quantity using Nanodrop (Thermo Fisher Scientific, Waltham, MA, USA) and the quality was evaluated by using the 2100 Bioanalyzer (Agilent Technologies, Santa Clara, CA, USA). The TruSeq Stranded mRNA Kit (Illumina, San Diego, CA, USA) was used to generate complementary DNA (cDNA) libraries. On the Illumina Novaseq6000 Platform (Illumina), 150 base pair (bp) paired-end sequencing of the resulting cDNAs was performed. To align sequences using the STAR software package (v.2.5.3a)^59^,, the mouse reference genome was obtained from the Ensembl database (GRCm38/mm10, Ensembl release 97)^60,61^. Htseq-count tool (v. 0.9.1) was used to count how many reads mapped to each genomic feature. From aligned sequences, transcript models (Ensembl release 97) were assembled and gene expression levels were quantified using the RSEM tool (v.1.3.0). Gene expression level was represented as Transcripts Per Kilobase Million (TPM).

### Proteomics analysis

The protein solutions prepared by metabolomics analysis were used. Proteins from frozen liver powder of Lys(6)-SILAC mice (Silantes) were extracted with methanol/chloroform extraction. Pre-chilled methanol, chloroform, and water were added to the frozen liver powder. The mixture was placed on ice for 30 minutes and centrifuged to remove the separated aqueous phase. Methanol was added to the organic and intermediate phases and centrifuged to precipitate the proteins. After washing with 80% methanol, protein pellet was resolved in 1%SDS and 50 mM Tris pH8.8. By calibrating to 1 mg/mL, bicinchoninic acid (BCA) assay (Thermo Fisher Scientific) was used to quantify the protein concentration of the lysates. Then, 50 µg of non-labeled and labeled protein extracts were combined in another tube for peptide preparation. We blocked cysteine residues using 2 mM Tris (2-carboxyethyl) phosphine hydrochloride (TCEP) (Thermo Fisher Scientific) at 37 °C for 30 min followed by alkylation with 10 mM 2-iodoacetamide (IAA) at room temperature for 30 min. By ultrasonically treating the pellet three times for 30 s with intervals of 30 s with a Bioruptor (Diagenode), the proteins precipitated with acetone for 3 h at −30 °C were dispersed in 50 mM triethylammonium bicarbonate. Lysyl endopeptidase (Wako, Osaka, Japan) was added to the protein suspension and digested at 37 °C for 16 h. Prior to MS analysis, the resulting peptides were purified using C18-StageTip purification after centrifuging at 15,000 *g* for 15 min at 4°C^62^. Before MS analysis, peptides were resolved in 50 µL 3% ACN-0.1% foromic acid.

Samples were analyzed on the Q Exactive HF-X mass spectrometer (Thermo Fisher Scientific) instrument equipped with the UltiMate 3000 RSLCnano LC system (Thermo Fisher Scientific) with a nano-electrospray source and a column oven set at 50°C (AMR Inc., Tokyo, Japan). Peptides were directly injected onto a 75 μm × 25 cm PicoFrit emitter (New Objective, Woburn, MA, USA) packed in-house with C18 core-shell particles (CAPCELL CORE MP 2.7 μm, 160 Å material; Osaka Soda Co., Ltd., Osaka, Japan) with a linear gradient of 7–42% B for 76 min, 42–80% B for 5 min, and 80% B for 9 min at a flow rate of 100 nL/min, where A is 0.1% formic acid and B is 0.1% formic acid and 80% acetonitrile. The data-independent acquisition (DIA) mode was used to conduct MS data acquisition. In the DIA mode for quantification, MS1 spectra were collected in the range of 423–865 m/z at 60,000 resolution to set an automatic gain control (AGC) target of 3 × 10^6^. MS2 spectra were collected in the range of >200 m/z at 30,000 resolution to set an AGC target of 3 × 10^6^. The isolation width was set to 8 m/z with stepped normalized collision energies of 20% and 23%. Isolation window patterns in 428– 860 m/z were used as window placements optimized by Skyline 4.1^63^. By using the above measurements and gas-phase fractionation method, the pool of all samples was analyzed in the DIA mode for a spectral library. In the gas-phase fractionation method, we used six MS ranges (426–502, 498–574, 570–646, 642–718, 714–790, and 786–862 m/z), and each was measured by overlapping DIA^64,65^. AGC target of 3 × 10^6^ was set by collecting MS1 spectra at 120,000 resolution. AGC target of 3 × 10^6^ was set by collecting MS2 spectra in the range of >200 m/z at 60,000 resolution. With stepped normalized collision energies of 20 and 23, the isolation width was set to 2 m/z. Window placements optimized by Skyline 4.1 were set as isolation window patterns in the ranges of 428–500, 500–572, 572–644, 644–716, 716–788, and 788–860 m/z.

With Scaffold DIA (Proteome Software Inc., Portland, OR, USA), the spectral library was built by searching MS data in the library against the mouse Ensembl release 97 database. The setting parameters were as follows: experimental data search enzyme, LysC/P; maximum missed cleavage sites, 1; precursor mass tolerance, 6 ppm; fragment mass tolerance, 8 ppm; and static modification, cysteine carbamidomethylation. The peptide identification threshold was a peptide false discovery rate (FDR) < 1%. For identification of peptides derived from Lys(6)-SILAC mice, a library of identified peptides with a mass increase 6.0201s29 Da on lysine was added to the library obtained above. DIA-NN (version 1.8) was used to process all raw data with library search mode using the parameters below: reaanotation, Mus_musculus.GRCm38.97.pep.all.fa; modification, UniMod188 (6.020129), and peak-translation. By using the output from DIA-NN software, we calculated the SILAC ratios of all assigned fragment ion pairs, and then the median of the SILAC ratios of each protein was used for protein quantification.

### Phosphoproteomics analysis

The protein solution prepared in metabolome analysis was used. Enrichment of phosphopeptides was conducted with some modifications as previously described ^66^. Briefly, TCEP was used to reduce cysteine residues in the protein samples from the liver tissues prepared above, after that alkylation of the thiol groups with IAA. By addition of ice-cold acetone, proteins were precipitated and then digested with trypsin in 100 mM ammonium bicarbonate. To enrich the phosphopeptides, the resulting peptides (1 mg) were subjected to Fe^3+^-IMAC. All samples were analyzed with Orbitrap Exploris 480 mass spectrometer (Thermo Fisher Scientific) equipped with the UltiMate 3000 RSLCnano LC system (Thermo Fisher Scientific) via a nano-electrospray source with a column oven set at 60°C (AMR Inc.). Peptides were directly injected onto a 75 μm × 120 mm column (Nikkyo Technos Co.,Ltd., Tokyo, Japan) with a linear gradient of 5–32% B for 70 min and 32–65% B for 10 min at a flow rate of 200 nL/min, where A is 0.1% formic acid and B is 0.1% formic acid and 80% acetonitrile.

MS data acquisition was conducted in overlapping DIA mode^64,65^. In the DIA mode for quantification, to set an AGC target of 3 × 10^6^, MS1 spectra were collected in the range of 390–1010 m/z at 30,000 resolution. To set an AGC target of 3 × 10^6^, MS2 spectra were collected in the range of >200 m/z at 30,000 resolution, and stepped normalized collision energies of 22%, 26%, and 30%. Window placements optimized by Skyline v4.1 were used with overlapping window patterns in 400-1000 m/z and the isolation width for MS2 set to 10 m/z^63^. In the DDA-MS mode for a spectral library, the pool of all samples was analyzed with the gas-phase fractionation method. The MS ranges of 395–555, 545–705, 695–1005 and 390–1010 m/z were respectively used and measured in DDA mode. To set an AGC target of 3 × 10^6^, MS1 spectra were collected at 120,000 resolution. In the MS ranges of 395–555, 545–705, 695–1005 m/z, the 20 most intense ions with charge states of 2^+^ to 5^+^ that exceeded 5.0 × 10^3^ were fragmented by higher collisional dissociation (HCD) with normalized collision energies of 22%, 26%, and 30%, and MS2 spectra were collected in the range of >200 m/z at 60,000 resolution to set an AGC target of 5 × 10^5^. In the MS ranges of 390–1010 m/z, the 40 most intense ions with charge states of 2^+^ to 5^+^ that exceeded 5.0 × 10^3^ were fragmented by HCD with normalized collision energies of 22%, 26%, and 30%, and MS2 spectra were collected in the range of >200 m/z m/z at 30,000 resolution to set an AGC target of 2 × 10^5^.

By searching MS data in the library against the mouse Ensembl database (GRCm38/mm10, Ensembl release 97) using Proteome Discoverer v2.3 (Thermo Fisher Scientific), the spectral library was generated. The setting parameters were as follows: Fragmentation, HCD; Precursor Tolerance, 8 ppm; Fragment Tolerance, 0.02 Da; Digestion Enzyme, Trypsin; Max Missed Cleavages, 1; Variable Modification, Phospho [S, t, Y]; Fixed Modification, Carbamidomethylation [C]; Site Probability Threshold, 75; Peptide FDR, < 1%.

### Metabolomics analysis

Metabolomics analysis was conducted as previously described^33,34^. By adding methanol:chloroform:water (2.5:2.5:1) extraction, total metabolites and proteins were extracted from the liver. To equalize peak intensities of mass spectrometry (MS) among runs, about 40 mg of frozen liver tissue was suspended with 500 μL precooled methanol with internal standards [20 μM L-methionine sulfone (Wako), 2-Morpholinoethanesulfonic acid (Dojindo), and D-Camphor-10-sulfonic acid (Wako)] to normalize. The mixture was kept in an ice-water bath, followed by the addition of 500 μL of chloroform, and 200 μL of water. Then the mixture was centrifuged at 4,600 g for 15 min at 4°C. The aqueous layer was filtered through a 5 kDa cutoff filter (Human Metabolome Technologies) to remove protein contamination. Finally, 320 μL filtrate was dried and stored at -80°C until analysis. Before MS analysis, samples were dissolved in 50 μL water containing reference compounds [200 μM each of trimesate (Wako) and 3-aminopyrrolidine (Sigma-Aldrich)]. Proteins were precipitated again by adding 800 μL of precooled methanol to the interphase and organic layers. The mixture was centrifuged at 12,000 g, 4°C, for 15 min. The resulting pellet was washed with 1 mL of ice-cold 80% (v/v) methanol and resuspended in 1 mL of sample buffer containing 1% SDS and 50 mM Tris-Cl pH8.8. The mixture was kept in an ice-water bath and homogenized with ultrasonication. The concentration of total protein was quantified by BCA assay and used to normalize metabolite concentration among samples. The resulting protein solution was used for proteomics analysis and phosphoproteomics analysis.

All CE-MS experiments were conducted based on the Agilent 1600 Capillary Electrophoresis system (Agilent Technologies), an Agilent 6230 TOF LC/MS system, an Agilent 1200 series isocratic pump, a G1603A Agilent CE-MS Adapter Kit, and a G1607A Agilent CE Electrospray Ionization (ESI)-MS Sprayer Kit. In order to analyze cationic compounds, a fused silica capillary (50 μm internal Diameter (i.d.) × 100 cm) was loaded with 1 M formic acid as the electrolyte. The sheath liquid was methanol/water (50% v/v) containing 0.01 μM hexakis (2,2-difluoroethoxy) phosphazene with flow rate set at 10 μL/min. Masses reference standards ([13C isotopic ion of a protonated methanol dimer (2CH3OH+H)]+, m/z 66.0631) and ([hexakis(2,2-difluoroethoxy) phosphazene +H]+, m/z 622.0290) were employed to automatically recalibrate each acquired spectrum. Electrospray ionization (ESI) positive ion mode with capillary voltage set as 4 kV was used for data acquisition. Metabolites were identified by comparing m/z values and relative migration times to the metabolite standards. By comparing peak regions to calibration curves generated using internal standardization techniques with methionine sulfone, quantification was carried out. The other conditions remained the same as reported before^67^. For analyzing anionic metabolites, 50 mM ammonium acetate solution (pH 8.5) as the electrolyte with a COSMO (+) (chemically coated with cationic polymer) capillary (50 μm i.d. x 105 cm) (Nacalai Tesque, Kyoto, Japan) were employed. The sheath liquid was methanol/5 mM ammonium acetate (50% v/v) containing 0.01 μM hexakis(2,2-difluoroethoxy) phosphazene with flow rate set at 10 μL/min. Masses reference standards ([13C isotopic ion of deprotonated acetate dimer (2CH3COOH-H)]-, m/z 120.0384) and ([hexakis (2,2-difluoroethoxy) phosphazene +deprotonated acetate (CH3COOH-H)]-, m/z 680.0355) were employed to automatically recalibrate each acquired spectrum. Data acquisition was performed in electrospray ionization (ESI) negative mode with capillary voltage set as 3.5 kV. D-Camphor-10-sulfonic acid was used as the internal standards for anion analysis. The other conditions were identical to those described previously^68^. The generated raw data were analyzed by using our proprietary software^69^.

### Lipidomics analysis

For lipid extraction, liver samples were prepared by the Bligh and Dyer’s method with minor modifications^70^.. Briefly, by adding 1 mL of cold methanol, the lipids and their related metabolites were extracted from frozen and crushed liver tissues (∼50 mg). The samples were thoroughly mixed for 1 min and sonicated for 5 min, followed by centrifuging at 16,000 ×*g* for 5 min at 4°C. The supernatant was collected. Protein concentrations in the pellet were measured with the Pierce™ BCA Protein Assay Kit (Thermo Fisher Scientific). The collected supernatant (180 μL) was mixed with 200 μL chloroform, 150 μL water, 10 μL internal standard (IS) A (Mouse SPLASH Lipidomix Mass Spec Standard, Avanti Polar Lipids Inc., Alabaster, AL, USA) containing 1.0 nmol phosphatidylcholine (PC) 15:0–18:1 (d7), 0.070 nmol phosphatidylethanolamine (PE) 15:0–18:1 (d7), 0.20 nmol phosphatidylserine (PS) 15:0–18:1 (d7), 0.050 nmol phosphatidylglycerol (PG) 15:0–18:1 (d7), 0.20 nmol phosphatidylinositol (PI) 15:0–18:1 (d7), 0.10 nmol phosphatidic acid (PA) 15:0–18:1 (d7), 0.45 nmol lysophosphatidylcholine (LPC) 18:1 (d7), 0.020 nmol lysophosphatidylethanolamine (LPE) 18:1 (d7), 2.5 nmol cholesteryl ester (CE) 18:1 (d7), 0.15 nmol diacylglycerol (DG) 15:0–18:1 (d7), 0.35 nmol triacylglycerol (TG) 15:0–18:1 (d7)–15:0, and 0.20 nmol sphingomyelin (SM) d18:1–18:1 (d9), 10 μL of IS B (Avanti Polar Lipids Inc.) containing 1.0 nmol monoacylglycerol (MG) 18:1 (d7), 0.10 nmol ceramide (Cer) d18:1 (d7)–15:0, 0.10 nmol hexosylceramide (HexCer) d18:1 (d7)–18:1, 1.1 nmol free fatty acid 16:0 (^13^C16), and 3.0 nmol cholesterol (d7), and 10 μL of IS C (Merck, Darmstadt, Germany) containing 0.40 nmol acylcoenzyme A (acyl-CoA) 2:0 (^13^C^2^). By vortexing and centrifugation at 16,000 ×g and 4 °C for 5 min, the aqueous and organic layers were collected.

The aqueous (upper) layer (250 μL) was transferred to a clean tube and analyzed by a LC (Nexera X2 UHPLC System, Shimadzu Co., Kyoto, Japan) with a metal-free peek-coated InertSustain C18 column (2.1 mm i.d. × 150 mm, 3 μm particle size, GL Sciences Inc., Tokyo, Japan) coupled with the Q Exactive, High-Performance Benchtop Quadrupole Orbitrap High-Resolution Tandem mass spectrometer (Thermo Fisher Scientific) (C18-LC/MS) for acyl-CoAs and acyl-carnitines^71^. By evaporating the aqueous layer extracts under a vacuum, the dried extracts were stored at -80°C. Before C18-LC/MS analysis, the dried extracts were reconstituted with 50 μL water. The organic (lower) layer (140 μL) was transferred to a clean tube and analyzed by a supercritical fluid chromatography (SFC, Nexera UC System, Shimadzu Co.) with ACQUITY UPC^2^ HSS C18 column (3.0 mm i.d. × 100 mm, 1.8 μm particle size, Waters, Milford, MA, USA) coupled with a triple quadrupole MS (TQMS, LCMS-8060, Shimadzu Co.) (C18-SFC/MS/MS) for fatty acids, TGs, and CEs^71^ and an SFC (Shimadzu Co.) with a ACQUITY UPC^2^ Torus diethylamine (DEA) (3.0 mm i.d. × 100 mm, 1.7 μm particle size, Waters) coupled with a TQMS (DEA-SFC/MS/MS) for PCs, PEs, PSs, PGs, PIs, PAs, LPCs, LPEs, MGs, DGs, SMs, cholesterol, Cers, and HexCers^71,72^. With a solution of methanol, the liver lipid extract (140 μL) was diluted to a final volume of 200 μL for C18-SFC/MS/MS and DEA-SFC/MS/MS analyses.

### Glycogen assay

Glycogen content was measured as previously described with some modifications^73^. About 20 mg liver sample was digested for 1 h at 95°C in 1 mL of 30% (w/v) potassium hydroxide solution and 50 μL of lysate was collected into another 1.5 mL tube before being neutralized in 15.3 μL of glacial acetic acid. By using BCA assay, the total protein concentration of the liver digest was measured and adjusted to 1 μg of protein per microliter. The Bligh and Dyer method was employed to remove lipids and extract glycogen from liver digest. The liver digest (50 μL) was then mixed with 120 μL ice-cold methanol, 50 μL chloroform, 10 μL 1% (w/v) linear polyacrylamide, and 70 μL of water. The final mixture was centrifuged at 12,000 g to remove the aqueous layer after 30 min ice-incubation. The glycogen was precipitated by adding 200 μL of methanol, centrifuging at 12,000 *g* for 30 min at 4°C, washing with ice-cold 80% (v/v) methanol, and completely drying. Glycogen pellets were suspended in 20 μL of amyloglucosidase (0.1 mg/mL; Sigma-Aldrich) in 50 mM sodium acetate buffer and incubated for 2 hours at 55°C to digest the glycogen. According to the manufacturer’s instructions, the concentration of glucose produced from the glycogen was evaluated by using the Amplex Red Glucose/Glucose Oxidase Assay Kit glucose assay (Thermo Fisher Scientific).

### Insulin assay and blood glucose measurement

The insulin concentration of mice plasma was measured using the LBIS Mouse Insulin ELISA Kit (Wako) according to the manufacturer’s protocol. U-type (633-03411) was used for WT mice and T-type (634-01481) was used for *ob*/*ob* mice. To avoid being out of the standard curb, all of samples were serially diluted before the experiments. The blood glucose level was measured using the ACCU-CHECK Aviva Nano meter system (Roche, Basel, Switzerland).

### Western blot analysis

Total proteins were extracted from the liver with methanol:chloroform:water (2.5:2.5:1) as previously described. About 40 mg liver with ice-cold methanol (500 μl) to achieve a concentration of 100 mg/mL the weight of the liver. The suspension was added into chloroform (400 μL) and water (200 μl), and then centrifugated at 4,600 *g* for 10 min at 4°C. Removing the aqueous and organic phases and added 800 μl of ice-cold methanol to the interphase to precipitate the proteins. The remaining pellet was suspended with 1000 µL lysis buffer (10 mM Tris-HCl [pH 6.8] in 1% SDS) and sonication. The total protein concentration of the resulting supernatant was measured by the BCA assay (Pierce BCA Protein Assay Kit, Thermo, 23227). The following antibodies were purchased from Cell Signaling Technology (Danvers, MA, USA): total Irs1 (#2382), total Erk1/2 (#9102), pErk1/2 (Thr202/Tyr204) (#9101), total eukaryotic translation initiation factor 4e (eIF4e) (#9742), peIF4e (Ser209) (#9741), total Akt (#9272), pAkt (Ser473) (#9271), total S6 (#2217), pS6 (Ser235/Ser236) (#2211), total Gsk3β (#9315), pGsk3β (Ser9) (#9336), total Ampkα (#2532), pAmpkα (Thr172) (#2531), total Foxo1 (#9462), and pFoxo1 (Ser256) (#9461). Antibody against pIrs1 (Tyr612) (09-432) was purchased from Millipore (Burlington, MA, USA). The proteins (20 μg–40 μg) were separated on SDS– polyacrylamide gel electrophoresis (PAGE) and blotted with the antibodies. Proteins were detected using the Immobilon Western Chemiluminescent HRP Substrate (Millipore) or SuperSignal West Pico PLUS Chemiluminescent Substrate (Thermo Fisher Scientific), and quantified using a luminoimage analyzer (Fusion System Solo 7S; M&S Instruments Inc., Osaka, Japan) and Fiji software (ImageJ; National Institutes of Health, Bethesda, MD, USA)^74^. Contrast and brightness adjustment and treatment were performed with Photoshop CS6 (Adobe).

## QUANTIFICATION AND STATISTICAL ANALYSIS

### Identification of differentially expressed or phosphorylated molecules between WT and *ob*/*ob* mice

For molecules measured in different omics, only those detected in more than 70% of replicates in either WT or *ob*/*ob* mice were used for further analyses. The differences in the abundance of molecules between WT and *ob*/*ob* mice were calculated by a statistical test using the following methods. The significance of difference (*p* value) was tested by the two-tailed Welch’s t-test, *q* value was calculated with the Storey procedure, for each protein expression, phosphorylation ratio (phosphorylation/total protein amount) measured by phosphoproteome, polar metabolite amount, and lipid amount. The *q* value of protein phosphorylation ratio (phosphorylation/total protein amount) measured by western blot analysis was calculated with the Benjamini-Hochberg (BH) procedure due to the small numbers of molecules (< 100). For transcriptomics analysis, the significance of difference (*p* value and *q* value) for gene expression was tested by the edgeR package (version 3.32.1) of the R language (version 4.0.3) with default parameters. The genes have sufficiently large counts to be retained in statistical analysis were determined by the function “filterByExpr”. Only the genes that pass the screening will be included in further analysis with their TPM values. The use of 0.05 as *p* value cut-off in scientific research is a controversial topic that has been debated by American Statistical Association^75^. One argument against the 0.05 cut-off is that it is often misused and considered arbitrary as *p* value indicates the degree of data compatibility with the null hypothesis. For clinical diagnosis, a lower *p* value is necessary to be conservative. In this study, we conducted targeted metabolome analysis, measuring 183 polar metabolites. To prevent excessive of false negatives, it is important to adjust our threshold based on the total number of measured molecules. Based on this, molecules with *q* value < 0.1 were defined as differentially expressed or phosphorylated molecules in *ob*/*ob* mice. Fold change (FC) value was computed by using amount of molecule in *ob*/*ob* mice to divide the amount of molecule in WT mice. Differentially expressed or phosphorylated molecules with FC values > 1 were defined as increased molecules in *ob*/*ob* mice, and FC values < 1 were defined as decreased molecules in *ob*/*ob* mice. For phosphoproteins, the ratios of phosphorylation to total amount of molecules were used (normalized phosphorylation site ratio = phosphorylation site value/protein value). One phosphoprotein may cover multiple phosphorylation sites. We defined a phosphoprotein with at least one changed phosphorylation site between WT and *ob*/*ob* mice as DPP. For the phosphoproteins including both increased and decreased phosphorylation sites in *ob*/*ob* mice, we defined them as increased and decreased DPPs in *ob*/*ob* mice.

### Inference of potential TFs for mRNA transcription

The ChIP-Atlas database was employed to estimate the potential TFs. An experimental list was downloaded and only data with mouse genome mm10 annotation was remained. By tracking the targeted experimental IDs, we downloaded experimental data registered in the ChIP-Atlas. We extracted flanking regions from -1000 bp to 1000 bp of the transcription start site in target genes. We excluded experimental subjects that were not liver but kept Foxo1 in all organs, because Foxo1 regulates metabolism in the liver although it was not measured in the liver in all ChIP-atlas datasets. We further set the TFs with any experiment signal larger than 50 as detected TFs.

For inference of differential regulations between regulating DRTFs and regulated DEGs, TF enrichment analysis was performed. The enrichment of TF was determined by the one-tailed Fisher’s exact test, and TFs with *q* value (calculated by Benjamini-Hochberg method) less than 0.1 were defined as significantly enriched. We used genes detected in more than 70% of the replicates in either WT or *ob*/*ob* mice as background. To avoid overestimation, of the enriched TFs with *q* < 0.1, we identified those included in DEGs or in DPPs with the same changed direction as the differentially expressed TFs (DETFs) or differentially phosphorylated TFs (DPTFs). In total, they were defined as DRTFs. The differential regulations from DRTFs to DEGs were determined based on the relationship between the TFs and the expression of DEGs described in the ChIP-Atlas database.

### Inference of potential protein kinases for protein phosphorylation

We predicted protein kinases regulating DPPs using the NetPhorest software. Potential protein kinases for phosphopeptides measured in the phosphoproteome were predicted based on the amino acid sequences of proteins by using online version of NetPhorest for humans with the default parameters. Input data for NetPhorest included DPPs with measured phosphorylation sites. Output data for NetPhorest included possible protein kinases with posterior probabilities of an amino acid residue being recognized by a protein kinase classifier (kinases with similar substrate recognition motifs). Among the candidate classifiers, we restricted “tree” into “KIN”. For each Netphorest Group of protein kinases, we selected the group with Netphorest Score above 0.3 in further analysis. To avoid overestimation, we identified those included in DEGs as the DEKs or in DPPs as DPKs. In total, they were defined as DRKs. The differential regulations from DRKs to DPPs were determined based on relationship between the protein kinases and the phosphorylation of DPPs described in Netphorest database.

### KEGG pathway enrichment analysis

The enrichment of genes or proteins in each pathway were performed by using one-tailed Fisher exact test. The background genes or proteins were set as all genes or proteins measured in each omics with more than 70% of replicates in either WT or *ob*/*ob* mice. The *q* values were calculated by BH procedure. The KEGG pathways with *q* values < 0.05 were defined as significantly enriched.

### Construction of the differential regulatory trans-omic network of metabolic reactions

The differential regulatory trans-omic network of metabolic reactions consisted of differentially expressed or phosphorylated molecules in nine omic layers: insulin signal, TF, protein kinase, enzyme mRNA, enzyme protein, enzyme phosphoprotein, metabolic reaction, metabolite, and lipid layers. Differential regulations connecting the differentially expressed or phosphorylated molecules across omic layers. By extracting a group of molecules from insulin signaling pathway (map04910), including the upstream of Creb from “The phosphatidylinositol 3’-kinase(PI3K)-Akt signaling pathway” (map04151) and “mitogen-activated protein kinase signaling pathway” (mmu04010) in the KEGG database, we created the insulin signaling layer. DPPs in insulin signaling layer are measured by phosphoproteome or western blot analysis. The TF layer consisted of all DRTFs inferred in this study. The protein kinase layer consisted of all DRKs inferred in this study. The enzyme mRNA layer, enzyme protein layer, enzyme phosphoprotein layer, metabolic reaction layer, metabolite layer, and lipid layer consisted of all the genes, proteins, and phosphoproteins of metabolic enzymes, metabolic reactions (based on EC number), metabolites and lipids in the “Metabolic pathways” (map01100) obtained from the KEGG database, including DEGs, DEPs, DPPs, expressed metabolic reactions, DEMs, and DELs respectively. To determine differential regulations between the regulating and regulated differentially expressed or phosphorylated molecules, we used the process that we have reported previously^33,34^. The direction of differential regulations (activating or inhibiting) was determined by the changed directions of the regulating and regulated molecules (increased or decreased). For the regulations from the enzyme mRNA layer to the enzyme protein layer, we determined the direction based on the correspondence between the DEGs encoding DEPs. The differential regulations from the enzyme protein layer and enzyme phosphoprotein layer to the metabolic reaction layer were determined by the corresponding metabolic enzymes using the KEGG database. For allosteric regulators to the metabolic reaction layer, we defined the activating regulations when the molecule as activator increased in *ob*/*ob* mice or that as inhibitor decreased in *ob*/*ob* mice. On the contrary, we defined the inhibiting regulations when the molecule as activator decreased in *ob*/*ob* mice or that as inhibitor increased in *ob*/*ob* mice. The information of activators or inhibitors in each allosteric regulation were obtained from the BRENDA database. We only include the allosteric regulations reported for mammals (Bos taurus, Felis catus, Homo sapiens, “Macaca,” “Mammalia,” “Monkey,” Mus booduga, Mus musculus, Rattus norvegicus, Rattus rattus, Rattus sp., Sus scrofa, “dolphin,” and “hamster”). Regulations by substrate or product were determined by the KEGG database. Because the reversibility of metabolic reactions was not clear, metabolic reactions were regarded to be regulated by both the substrate and product.

### Condensation of the differential regulatory trans-omic network of metabolic reactions into activating and inhibiting condensed networks

Differential regulatory trans-omic network was condensed into activating and inhibiting condensed network in *ob*/*ob* mice. According to the KEGG metabolic pathways, the related metabolic reactions in a specific metabolic pathway were grouped into one single node. We only kept the metabolic pathways with more than 35 regulations and created a pathway layer. We classified the metabolic pathway nodes into four classes, carbohydrate, amino acid, nucleotide, and lipid according to the KEGG database, and used dashed frames to circle them. Next, we extracted enzyme proteins, enzyme phosphoproteins, metabolites and lipids connected with each pathway, enzyme mRNAs, protein kinases, and TFs connected with enzyme proteins, enzyme phosphoproteins, and enzyme mRNAs respectively, and counted their regulation numbers. Finally, we removed the proteins and phosphoproteins with less than 1 regulations, metabolites with less than 10 regulations, lipids with less than 5 regulations, and TFs with less than 10 regulations.

### Extraction of specific metabolic pathways from the differential regulatory trans-omic network of metabolic reactions

Similar with “Condensation of the differential regulatory trans-omic network of metabolic reactions into activating and inhibiting condensed networks”, we downloaded different metabolic pathways from the KEGG database. We gathered metabolic reaction information in each metabolic pathway, and extracted enzyme proteins, enzyme phosphoproteins, metabolites, and lipids connecting with these metabolic reactions from the trans-omic network we constructed. And then, we extracted enzyme mRNAs, protein kinases, and TFs connecting with target enzyme proteins, enzyme phosphoproteins, and enzyme mRNAs respectively. We kept regulation information (edge) the same as trans-omic network. Finally, we added bar plot of each molecule in each metabolic pathway. We are displaying the same data in several individual metabolic networks.

### Comparison of the differential regulatory trans-omic networks of metabolic reactions between *ad libitum* feeding and 16 h-fasting in WT and *ob*/*ob* mice

C57BL/6J mice at *ad libitum* feeding in this study and 16 h-fasting in the previous study were cultured under the same condition^34^. Omic measurements were performed separately in two different batches. To avoid the batch effect, we analyzed trans-omic network at each feeding condition independently by using almost the same analytical method. Some slight differences on analysis methods were described in the following section. We slightly modified the differential regulatory trans-omic network at 16 h-fasting by renewing TF estimation using ChIP-Atlas database, and by renewing allosteric regulations and metabolic information using the same version of BRENDA and KEGG database in this study. We also removed the Protein Kinase layer, Enzyme Phosphoprotein layer, and Lipid layer in the differential regulatory trans-omic network at *ad libitum* feeding of this study, which were not in the differential regulatory trans-omic network at 16 h-fasting. In addition, we removed insulin signal layer in both differential regulatory trans-omic networks. In differential regulatory trans-omic network, the detailed numbers and percentages of nodes and edges were shown. For metabolic pathway comparison, first, we downloaded different metabolic pathways from the KEGG database. Second, we gathered metabolic reaction information in each metabolic pathway, and extracted enzyme proteins and metabolites connecting with these metabolic reactions from the trans-omic network we constructed. Then, we extracted related enzyme mRNAs and TFs connecting with target enzyme proteins. Third, we condensed TFs, enzyme mRNAs, enzyme proteins, metabolites (allosteric regulators) into one node according to their changed directions in *ob*/*ob* mice.

### Implementation

Statistical analyses and trans-omic network analysis were performed using MATLAB 2021a (The Mathworks Inc.), R language (version 4.0.3). Visualization of trans-omic network in Graph Modeling Language (GML) formats was performed using Python (version 2.7) and VANTED (version 2.8.3)^76^.

## EXCEL TABLE TITLES AND LEGENDS

**Table S1. Transcriptomic analysis in the liver of WT and *ob*/*ob* mice at *ad libitum* feeding. Related to Figure 2A.**

**Table S2. Proteomic analysis in the liver of WT and *ob*/*ob* mice at *ad libitum* feeding. Related to Figure 2B.**

**Table S3. Phosphoproteomic analysis in the liver of WT and *ob*/*ob* mice at *ad libitum* feeding. Related to Figures 2C and 2D.**

**Table S4. Western Blot data in the liver of WT and *ob*/*ob* mice at *ad libitum* feeding. Related to Figure 2D.**

**Table S5. Metabolomic analysis in the liver of WT and *ob*/*ob* mice at *ad libitum* feeding. Related to Figure 2E.**

**Table S6. Lipidomic analysis in the liver of WT and *ob*/*ob* mice at *ad libitum* feeding. Related to Figure 2F.**

**Table S7. Transcription factor enrichment analysis in the liver of WT and *ob*/*ob* mice at *ad libitum* feeding. Related to Figure 2G.**

We identified differential regulations from TFs to DEGs based on the connections that DRTFs are regulating DEGs inferred by the ChIP-Atlas database. We identified 352 increased and 36,098 decreased TF to DEG regulations in *ob*/*ob* mice.

**Table S8. Protein kinase inference in the liver of WT and *ob*/*ob* mice at *ad libitum* feeding. Related to Figure 2H.**

We identified differential regulations from protein kinases to DPPs based on the relationship that DRKs are activating DPPs inferred by the Netphorest database. We identified 45 increased and 33 decreased DRK to DPPs regulations in *ob*/*ob* mice.

**Table S9. The numbers of activating regulations and inhibiting regulations on metabolic reactions in *ob*/*ob* mice at *ad libitum* feeding. Related to Figure 4A.**

