## Supplemental Text and Figures for "Trans-omic analysis reveals opposite metabolic dysregulation between feeding and fasting in liver associated with obesity"

**Supplemental Information**

**Supplemental Text**

Identification of DEMs and DELs to metabolic reactions in the liver between WT and *ob*/*ob* mice at *ad libitum* feeding (Figure 3)

A DEM or DEL can be the activator for some enzymes and the inhibitor for other enzymes. Combined with the changed direction in DEM or DEL, we identified the regulation regulated by DEM or DEL as the activating or inhibiting regulation. DEMs and DELs that regulate metabolic reactions as substrates or products evoked the same magnitude of activating and inhibiting regulations (Figures 3F and 3G). Some DEMs, including ATP and ADP, are also involved in substrates or products regulations, with ATP contributing as decreased substrates (inhibiting regulations) and ADP contributing as decreased products (activating regulations) (Figure S3A, bottom left). For allosteric regulations regulated by DEMs, we identified 402 metabolic enzymes regulated by 18 increased DEMs and 100 decreased DEMs that functioned as allosteric regulators (Figure 3D). For metabolic reactions affected by DELs, Cer, PI, DG, and PE were involved in allosteric regulations and regulations affected by the substrate or product (Figure S3A). For allosteric regulations regulated by DELs, we identified 64 metabolic enzymes regulated by 4 increased DELs and 5 decreased DELs, which functioned as allosteric regulators (Figure 3F).

Regulation of metabolic pathways by regulators in differential regulatory trans-omic network of metabolic reactions at *ad libitum* feeding (Figure 4)

Most carbohydrate metabolism, including glycolysis/gluconeogenesis, the TCA cycle; amino acid metabolism, including BCAA degradation; nucleotide metabolism, including purine and pyrimidine metabolism; and some lipid metabolisms, including fatty acid metabolism, were mainly activated by metabolites and enzyme proteins in *ob*/*ob* mice (Figure S3C). Similarly, most carbohydrate metabolism, some Amino acids, lipid metabolism, and two nucleotide metabolisms were inhibited by metabolites. Enzyme phosphoproteins regulated fatty acid degradation and tryptophan metabolism, glycolysis/gluconeogenesis. Lipids mainly regulated lipid and carbohydrate metabolisms. Enzyme proteins mainly inhibited steroid hormone biosynthesis and purine metabolism (Figure S3C). Increased DRTF and decreased DRTFs revealed different regulation patterns on metabolic pathways (Figure S3D). Increased DRTF *Pparg* is associated with glycolysis/gluconeogenesis, BCAA degradation, fatty acid degradation, pyruvate, and steroid hormone biosynthesis. Decreased DRTFs such as *Cebpa*, *Hnf4a*, and *Foxa1*, significantly impacted tryptophan, purine, and steroid hormone biosynthesis. *Cebpa*, *Rxra*, *Brd4*, and *Cebpb* were significantly associated with steroid hormone biosynthesis ($q < 0.01$).

<u>Condensation of the differential regulatory trans-omic network of metabolic reactions into activating and inhibiting condensed networks at *ad libitum* feeding (Figure 5)</u>

We classified metabolic pathways into four classes: Lipid, Nucleotide, Amino acid, and Carbohydrate (dashed boxes). Because of no regulation from insulin signaling molecules to DPTF and DPK, we omitted the Insulin Signal layer in this analysis. The activating condensed network included differentially expressed or phosphorylated molecules (DRTFs, DEGs, DRKs, DEPs, DPPs, DEMs, and DELs) and activated metabolic pathways by activating regulations in *ob*/*ob* mice (Figure 5A). The inhibiting condensed network included differentially expressed or phosphorylated molecules and inhibited metabolic pathways by inhibiting regulations in *ob*/*ob* mice (Figure 5B). Some metabolic pathways were activated and inhibited by molecules at the same time, so they are both shown in activating and inhibiting condensed networks. Some DEMs or DELs oppositely regulate metabolic pathways, *e.g.*, decreased DEMs and DELs as allosteric inhibitors activate metabolic pathways (Figures 3D-3G and S1), which is why some blue nodes are connecting with red edges (Figure 5A), similar to Figure 5B. Only metabolic pathways regulated by more than 35 regulations, proteins and phosphoproteins regulated by more than 1 regulations, metabolites regulated by more than 10 regulations, lipids regulated by more than 5 regulations, and TFs regulated by more than 10 regulations, and mRNAs and protein kinases regulated by at least one regulation, are shown in this graph.

Comparison of the differential regulatory trans-omic networks of metabolic reactions between *ad libitum* feeding and 16 h-fasting in WT and *ob*/*ob* mice (Figures 6 and 7)

*Differential regulatory trans-omic networks comparison*

Overall, the percentages of the decreased TFs, the changed Enzyme mRNAs and Enzyme Proteins, the increased Metabolites in *ob*/*ob* mice were less at *ad libitum* feeding than at 16 h-fasting (Figure 6A). Most of the changed molecules intersected among increased molecules or decreased molecules except metabolites (Figure S6D). For TF regulations to Enzyme mRNA, the numbers of inhibiting regulations in *ob*/*ob* mice were less at *ad libitum* feeding than at 16 h-fasting. We found the dysregulation of enzyme proteins involved in glycolysis/gluconeogenesis, TCA cycle, fatty acid metabolism, and BCAA degradation (Figure S6C), so we further examined the difference in these metabolic pathways in detail (Figures 6B, 7A, and S7).

*Glycolysis/gluconeogenesis comparison*

Lactate increased in *ob*/*ob* mice at both *ad libitum* feeding and 16 h-fasting (Figure 6B). Acetyl-CoA increased only at *ad libitum* feeding. For rate-limiting enzymes specific to glycolysis, Gck and Pklr increased at both *ad libitum* feeding and 16 h-fasting. Pfkl and Pkm increased only at 16 h-fasting. For rate-limiting enzymes specific to gluconeogenesis, Pcx and Fbp1 increased at both *ad libitum* feeding and 16 h-fasting. There were less activating enzyme protein regulations and less inhibiting TF regulations in *ob*/*ob* mice at *ad libitum* feeding than at 16 h-fasting.

### Supplemental Figure

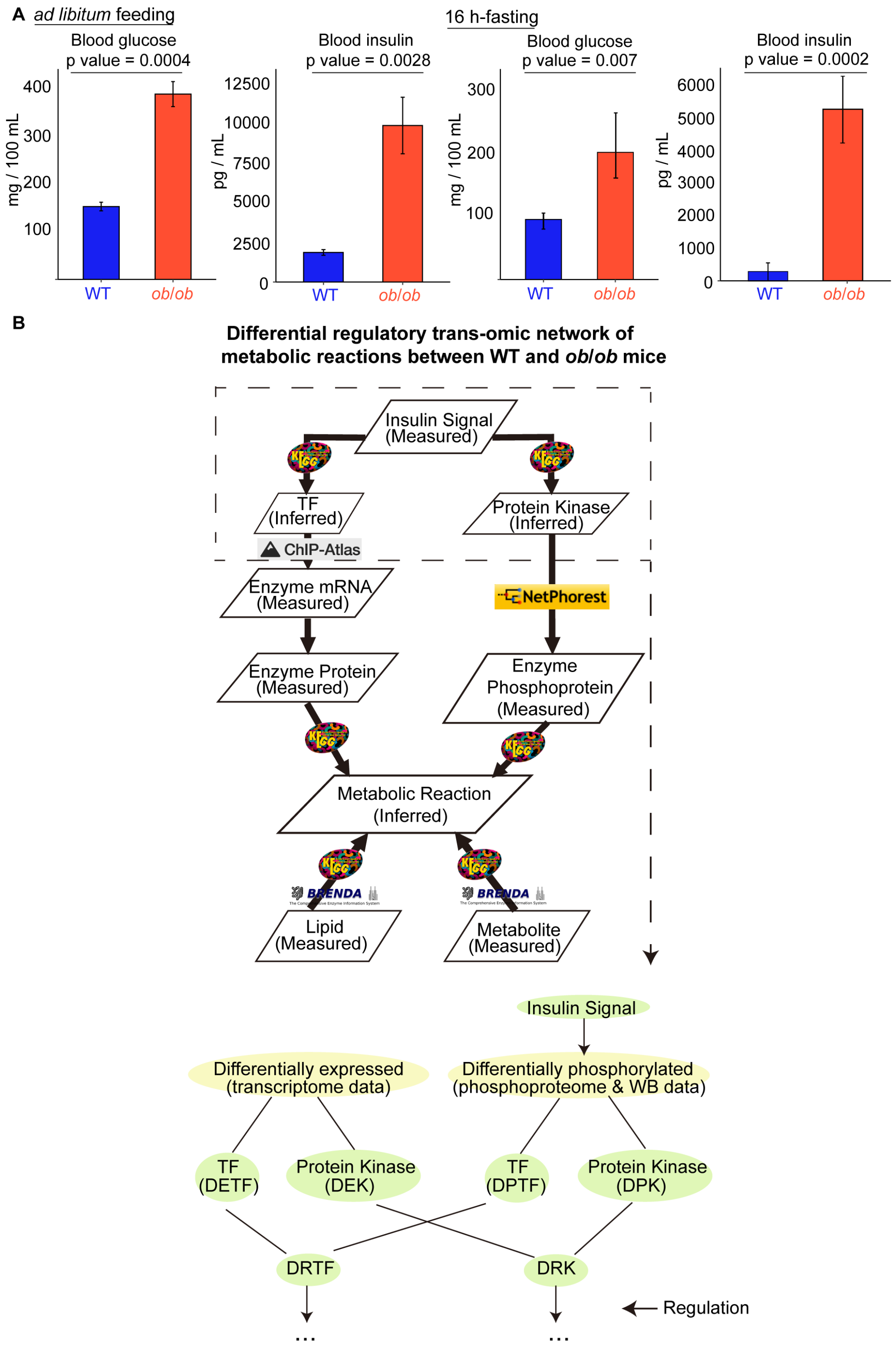

C

**Definition of contents in differential regulatory trans-omic network of metabolic reactions**

**Differentially expressed or phosphorylated molecules**

● **Insulin Signal**
Insulin signaling molecules of differentially phosphorylated proteins (DPPs)

● **Transcription Factor (TF)**
Differentially regulated TFs (DRTFs): Differentially expressed TFs (DETFs) and differentially phosphorylated TFs (DPTFs)

● **Enzyme mRNA**
Differentially expressed genes (DEGs) encoding metabolic enzymes

● **Enzyme Protein**
Differentially expressed proteins (DEPs) encoding metabolic enzymes

● **Protein Kinase**
Differentially regulated kinases (DRKs): Differentially expressed kinases (DEKs) and differentially phosphorylated kinases (DPKs)

● **Enzyme Phosphoprotein**
Differentially phosphorylated proteins (DPPs) encoding metabolic enzymes

● **Metabolic Reaction**
Differentially regulated metabolic reactions

● **Metabolite**
Differentially expressed metabolites

● **Lipid**
Differentially expressed lipids

**Differential regulations**

Regulations between differentially expressed or phosphorylated molecules in adjacent layers

**Using database for identification of differential regulations**

**From Insulin Signal to TF (phosphorylation)**
✓ KEGG

**From Insulin Signal to Protein Kinase**
✓ KEGG

**From TF to Enzyme Protein**
✓ Chip-Atlas

**From Protein Kinase to Enzyme Phosphoprotein**
✓ Netphorest

**From Enzyme Protein to Metabolic Reaction**
✓ KEGG

**From Enzyme Phosphoprotein to Metabolic Reaction**
✓ KEGG

**From Metabolite to Metabolic Reaction**
✓ BRENDA (for Allosteric regulation)
✓ KEGG (for Regulation by substrate or product)

**From Lipid to Metabolic Reaction**
✓ BRENDA (for Allosteric regulation)
✓ KEGG (for Regulation by substrate or product)

D

| | | TF (Phosphorylation) | | Enzyme mRNA | | Enzyme Protein | | Phosphoprotein (site) | | Metabolic Reaction |
|---|---|---|---|---|---|---|---|---|---|---|
| | | Increased | Decreased | Increased | Decreased | Increased | Decreased | Increased | Decreased | |
| Insulin Signal (Activator) | Increased | → | – | | | | | | | |
| | Decreased | – | → | | | | | | | |
| Insulin Signal (Inhibitor) | Increased | – | → | | | | | | | |
| | Decreased | → | – | | | | | | | |
| TF | Activated | | | → | – | | | | | |
| | Inhibited | | | – | → | | | | | |
| Enzyme mRNA | Increased | | | | | → | – | | | |
| | Decreased | | | | | – | → | | | |
| Protein Kinase | Increased | | | | | | | → | – | |
| | Decreased | | | | | | | – | → | |
| Enzyme Protein | Increased | | | | | | | | | → |
| | Decreased | | | | | | | | | → |
| Enzyme Phosphoprotein | Increased | | | | | | | | | → |
| | Decreased | | | | | | | | | → |
| | Both | | | | | | | | | → |
| Metabolite / Lipid (Allosteric activator) | Increased | | | | | | | | | → |
| | Decreased | | | | | | | | | ⊣ |
| Metabolite / Lipid (Allosteric inhibitor) | Increased | | | | | | | | | ⊣ |
| | Decreased | | | | | | | | | → |
| Metabolite / Lipid (Substrate) | Increased | | | | | | | | | → |
| | Decreased | | | | | | | | | ⊣ |
| Metabolite / Lipid (Product) | Increased | | | | | | | | | ⊣ |
| | Decreased | | | | | | | | | → |

**Figure S1**

**Figure S1. Contents of the differential regulatory trans-omic network of metabolic**

**reactions between WT and *ob*/*ob* mice. Related to Figure 1.**

(A) The concentrations of blood glucose and blood insulin of WT and *ob*/*ob* mice at *ad libitum* feeding (left) and 16 h-fasting (right)[1]. Blue, WT mice; red, *ob*/*ob* mice. The means and standard errors of the means (SEMs) of 5 mice were shown.

(B) The schema of constructing the differential regulatory trans-omic network of metabolic reactions between WT and *ob*/*ob* mice (top). The lower panel describes Insulin Signal regulates Differentially phosphorylated TF and Protein Kinase (DPTF and DPK).

(C) The definition of differentially expressed or phosphorylated molecules in each layer (left), differential regulations between adjacent layers and the databases employed in differential regulatory trans-omic network (right).

(D) Classification of the differential regulations (activating or inhibiting in *ob*/*ob* mice) based on the changed direction of differentially expressed or phosphorylated molecules, and characters of insulin signal, metabolite, and lipid (activator or inhibitor, substrate or product). The methodology of constructing differential regulatory trans-omic network of metabolic reactions is based on our previous study[1], and in further add Protein Kinase layer, Enzyme Phosphoprotein layer, and Lipid layer.

**A**

| | Transcriptome | | Proteome | |
|---|---|---|---|---|
| | *ad libitum* feeding | 16 h-fasting | *ad libitum* feeding | 16 h-fasting |
| Increased in *ob/ob* mice | - | SNARE interactions in vesicular transport * | Proteasome * | - |
| | - | DNA replication * | - | - |
| Decreased in *ob/ob* mice | Ribosome *** | Spliceosome ** | Ribosome *** | Ribosome *** |
| | Spliceosome *** | Protein processing in endoplasmic reticulum ** | Spliceosome *** | Spliceosome ** |
| | mRNA surveillance pathway * | | Protein processing in endoplasmic reticulum *** | Protein processing in endoplasmic reticulum ** |
| | | | Protein export * | |
| | | | Ribosome biogenesis * | |

q < 0.001 ***, 0.001 < q < 0.01 **, 0.01 < q < 0.05 *

**B**

Identify DPSs
2434 normalized phosphorylation sites

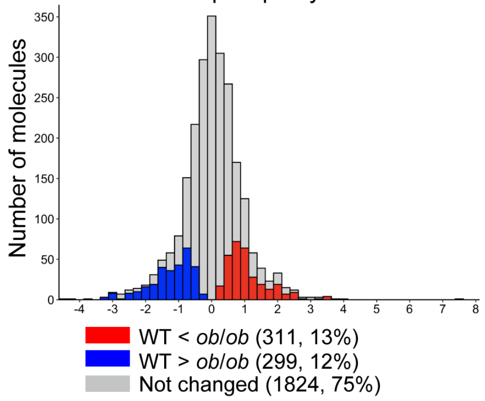


For each phosphoprotein, at least one phosphorylation site is measured

- All sites increased in *ob/ob* mice
  → Increased DPP in *ob/ob* mice 140 (19%)
- All sites decreased in *ob/ob* mice
  → Decreased DPP in *ob/ob* mice 126 (18%)
- Contains both increased and decreased sites in *ob/ob* mice
  → Increased and decreased DPP in *ob/ob* mice 46 (6%)
- None of site changed
  → not chagned phosphoprotein 412 (57%)

**C**

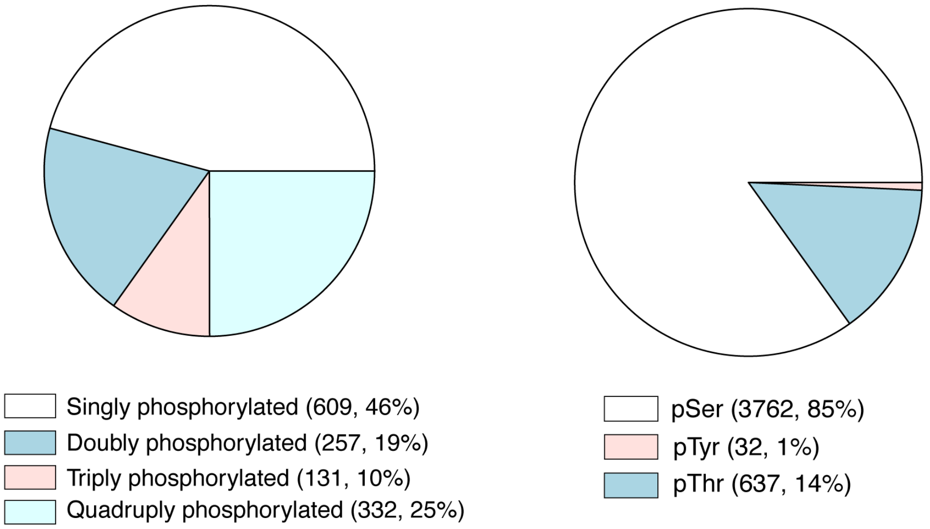


1

D

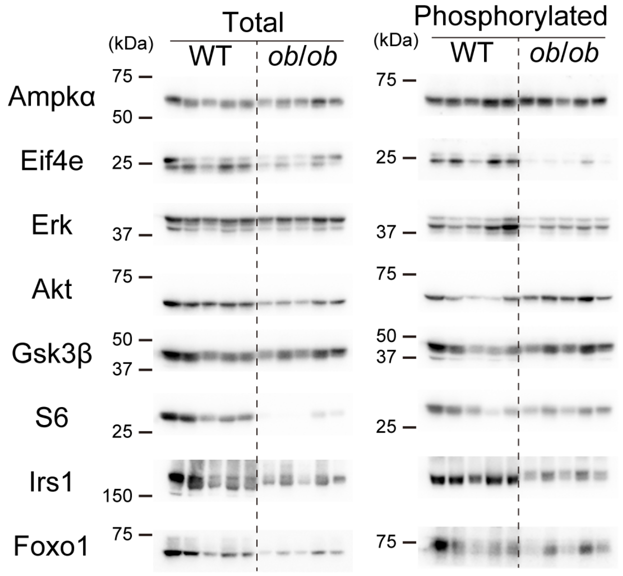

Total
Phosphorylated
(kDa)
WT
*ob/ob*
Ampkα
Eif4e
Erk
Akt
Gsk3β
S6
Irs1
Foxo1
75
50
25
37
150


E

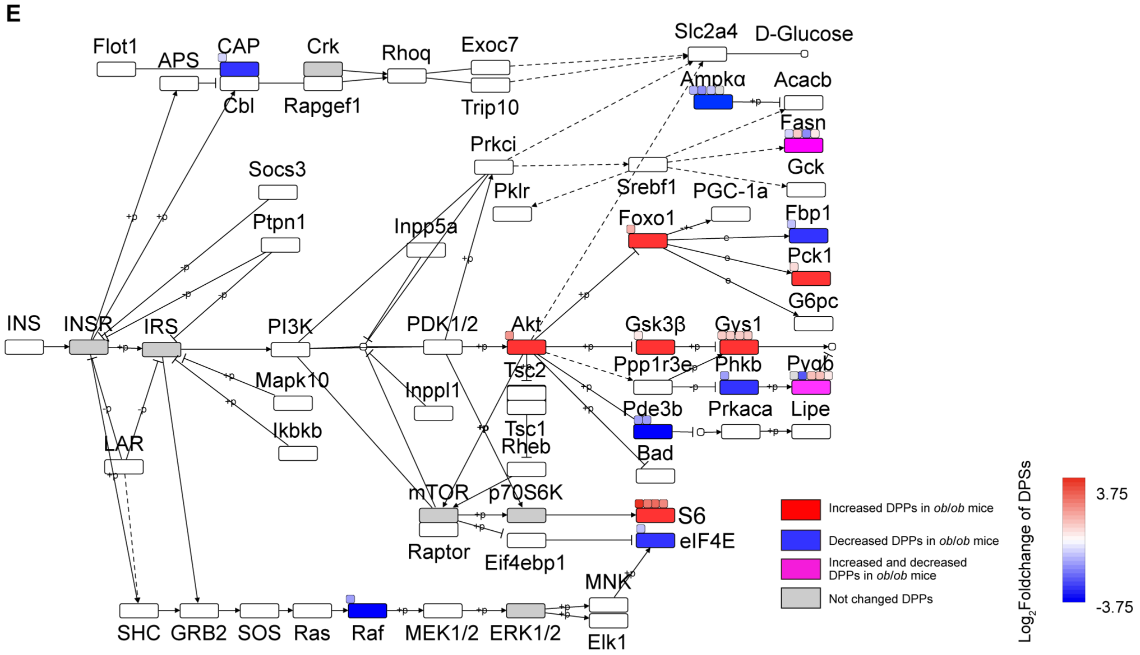

Flot1
APS
CAP
Cbl
Crk
Rapgef1
Rhoq
Exoc7
Trip10
Slc2a4
D-Glucose
Ampkα
Acacb
Fasn
Gck
Prkci
Srebf1
PGC-1a
Pklr
Socs3
Ptpn1
Inpp5a
Foxo1
Fbp1
Pck1
G6pc
INS
INSR
IRS
PI3K
PDK1/2
Akt
Gsk3β
Gys1
Tsc2
Ppp1r3e
Phkb
Pygb
Mapk10
Inppl1
Ikbkb
Tsc1
Rheb
Pde3b
Prkaca
Lipe
Bad
LAR
mTOR
p70S6K
Raptor
S6
eIF4E
Eif4ebp1
MNK
SHC
GRB2
SOS
Ras
Raf
MEK1/2
ERK1/2
Elk1
Increased DPPs in *ob/ob* mice
Decreased DPPs in *ob/ob* mice
Increased and decreased DPPs in *ob/ob* mice
Not changed DPPs
Log2Foldchange of DPSs
3.75
-3.75

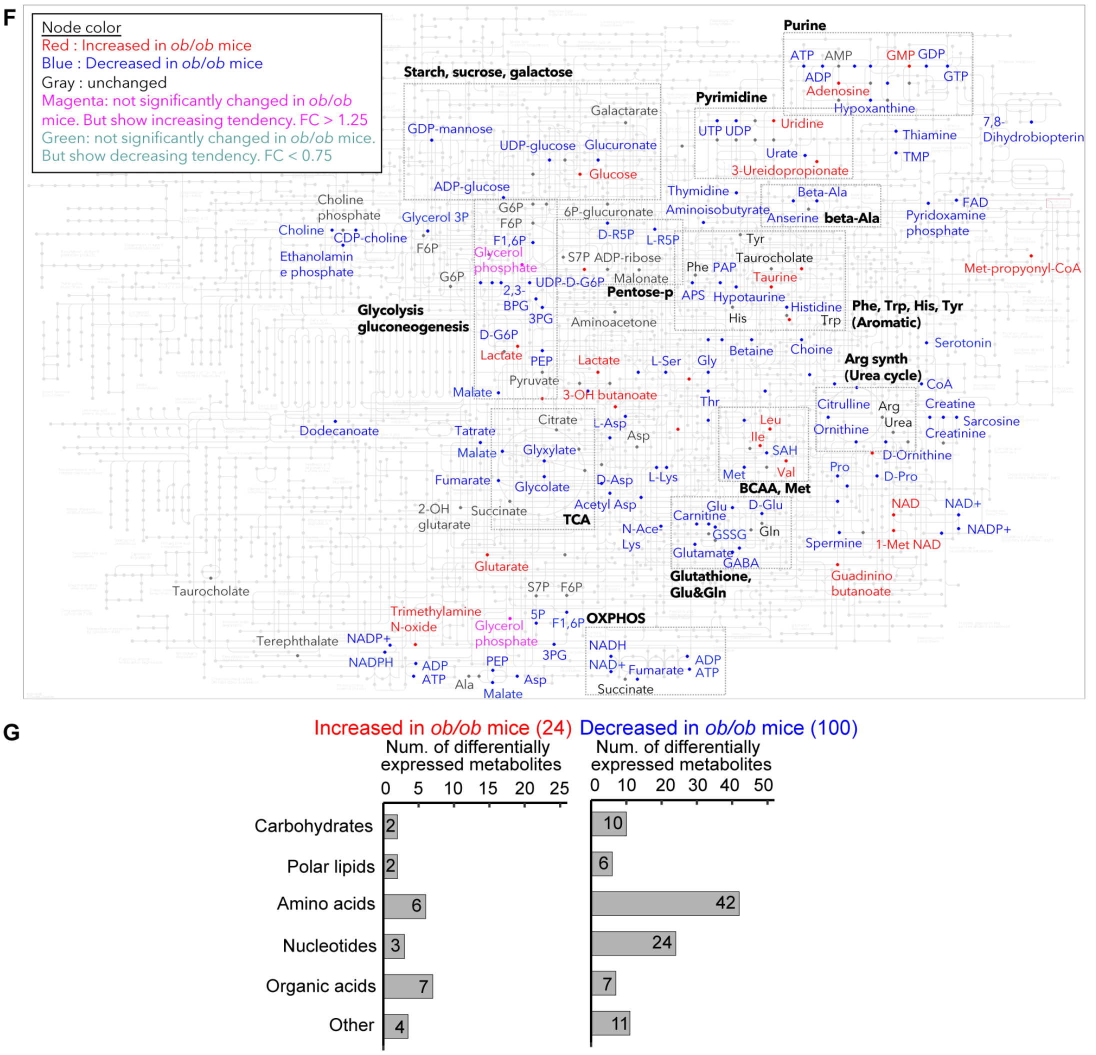


Figure S2

**Figure S2. Single omic analysis at *ad libitum* feeding. Related to Figure 2.**

(A) Summary of KEGG enrichment analysis results belonging to Genetic Information Process in transcriptome and proteome at *ad libitum* feeding and 16 h-fasting. Ribosome and Spliceosome were highlighted in red. The number of stars indicates the q value level of each KEGG pathway. KEGG enrichment analysis results belonging to the Genetic Information Process in transcriptome and proteome at *ad libitum* feeding and 16 h-fasting revealed ribosome and spliceosome pathways decreased on transcriptome and proteome levels, suggesting the amount and diversity of proteins potentially decreased in *ob*/*ob* mice. This result is consistent with the previous studies that the synthesis of ribosomes or related mRNA and protein expression are downregulated in the liver and muscle of *ob*/*ob* mice[2], high-fat diet mice[3] and diabetic rats[4]. The splicing machinery dysregulated in obesity-related metabolic diseases[5,6]. See also Tables S1 and S2. Related to Figures 2A and 2B.

(B) and (C) Characterization of phosphoprotein data. (B) Histogram of log2FC of phosphorylation amount of *ob*/*ob* mice compared to WT mice in phosphorylation site (left). Definition of four phosphoprotein groups and their numbers and percentages (right). (C) Distribution of phosphorylation sites per peptide (left) and amino acid residues per phosphorylation sites (right). Numbers indicate the number of phosphorylation sites or phosphopeptides. Percentage relative to the total phosphorylation sites or phosphopeptides, respectively. See also Table S3. Related to Figure 2C. We measured levels of phosphorylation at 4,120 sites using data-independent acquisition mass spectrometry (Figure 2C and Table S3). We used corresponding protein levels to normalize phosphoprotein levels (normalized phosphorylation amount = phosphorylation amount/protein amount). After normalization, we obtained 2,434 phosphorylation sites, 1,329 phosphopeptides, and 724 phosphoproteins for further analysis (Figure S2B). One phosphoprotein could cover multiple phosphorylation sites. We identified phosphoproteins with at least one changed phosphorylation site in *ob*/*ob* mice compared to WT mice as Differentially Phosphorylated Proteins (DPPs). For those phosphoproteins that included both increased and decreased phosphorylation sites, we defined them as increased and decreased DPPs. We identified 311 (13%) increased and 299 (12%) decreased differentially phosphorylated phosphorylation sites (DPSs). Singly phosphorylated peptides represented 609 (46%) of 1,329 phosphorylation peptides (Figure S2C). The measured phosphorylation sites included 3,762 (85%) phosphoserines (pSer).

(D) Western Blot analysis of the insulin signaling molecules in the liver of WT and *ob*/*ob* mice at *ad libitum* feeding. The phosphoproteins that were not detected in the phosphoproteome but are well-known key insulin signaling proteins (10 proteins), they were measured total amounts and phosphorylation amounts by Western Blot with 5 replicates. The ratio of phosphorylated protein to total protein for each pair of protein was calculated. The unabbreviated name of the molecules can be found in Table S4. Related to Figure 2D.

(E) Differential phosphorylated phosphoproteins in *ob*/*ob* mice at *ad libitum* feeding on "Insulin signaling pathway" (map04910) by the KEGG database. The squares above each rectangle indicate differential phosphorylated phosphorylation sites (DPSs). The color indicates the log2 FC value of each DPS in *ob*/*ob* mice compared to WT mice. Red, increased DPPs in *ob*/*ob* mice; Blue, decreased DPPs in *ob*/*ob* mice; Magenta, both increased and decreased DPPs in *ob*/*ob* mice. Gray, unchanged phosphoproteins. We found that phosphorylation of ribosomal protein S6 (pS6) (Phosphoprotein names are shown with prefix "p") increased (Figure 2D), while many ribosomal proteins, including S6, decreased on transcriptome and proteome level (Tables S1 and S2). pAkt increased, leading to a wide range of disturbances on downstream (Figure S2E). Increase of pFoxo1 suggested its translocation from nucleus to cytosol, resulting in down-regulation of its target genes in glycolysis and gluconeogenesis pathway in *ob*/*ob* mice[7]. Gsk3β activity was inhibited through increased phosphorylation at Ser9. pGsk3β inactivated Gys2[8]. pGys2 at Ser7 caused enzyme inactivation by decreasing affinity for both UDP-glucose and glucose-6-phosphate (G6P)[9]. Phosphorylation of Ampkα increases its kinase activity, leading to phosphorylation of multiple downstream targets and switching off anabolic pathways (e.g., fatty acid and sterol synthesis)[10]. pAmpkα decreased, suggesting that Ampkα does not inhibit anabolic pathways, such as fatty acid synthesis.

(F) Differential expressed metabolites (nodes) in *ob*/*ob* mice on "Metabolic pathways" (map01100) by the KEGG database. Dashed frames roughly show the class of metabolites. Red, increased metabolites in *ob*/*ob* mice; Blue, decreased metabolites in *ob*/*ob* mice; Gray, unchanged metabolites; Magenta, metabolites which not significantly changed in *ob*/*ob* mice but show increasing tendency (FC > 1.25); Green, metabolites which not significantly changed in *ob*/*ob* mice but show decreasing tendency (FC < 0.75).

(G) The DEMs numbers classified by metabolite class using the KEGG database. Among the decreased DEMs, "amino acids, peptides, and analogues" (42) and "nucleosides, nucleotides, and analogues" (24) took the great proportions. Related to Figure 2E.

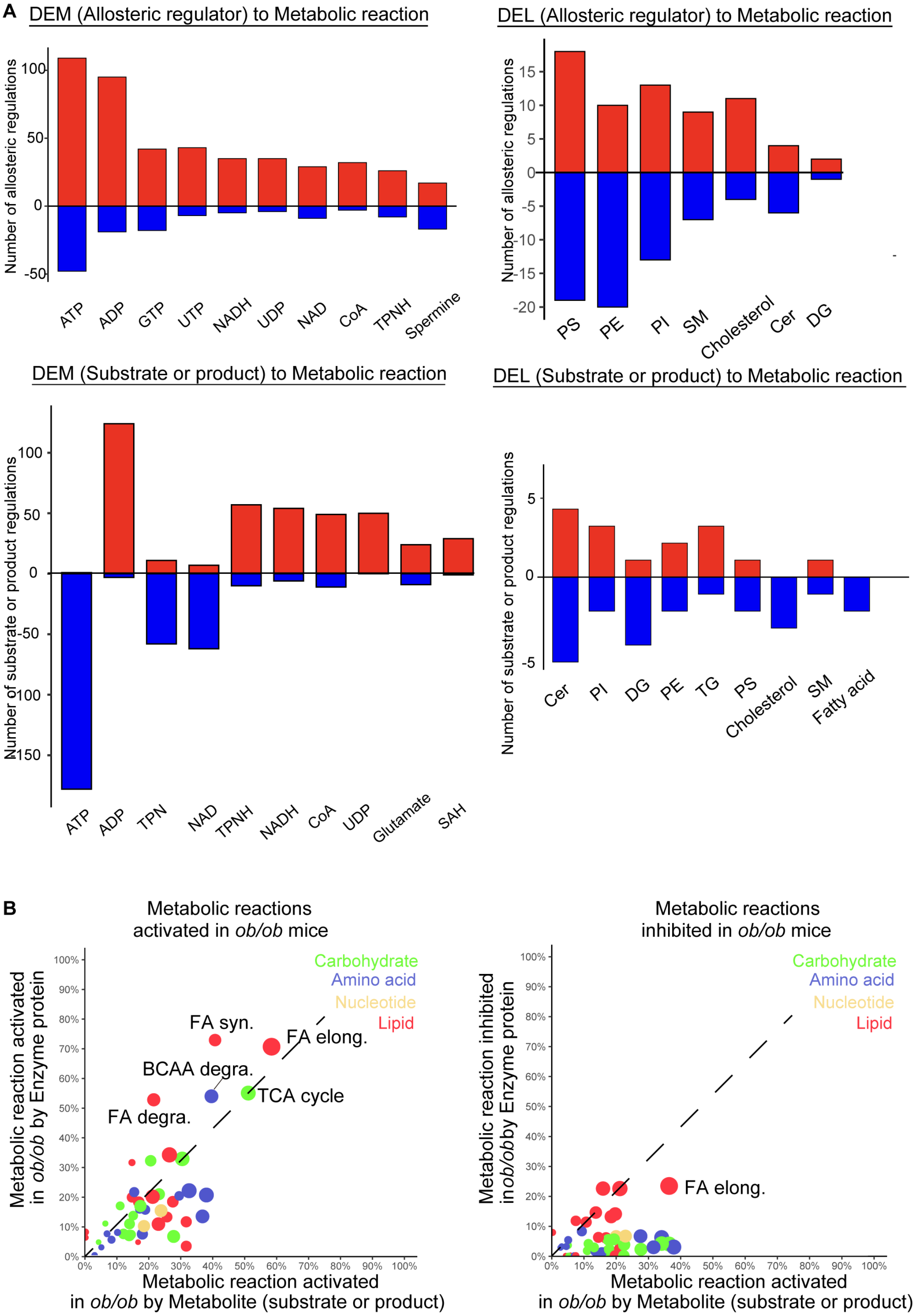
A
DEM (Allosteric regulator) to Metabolic reaction
Number of allosteric regulations
ATP
ADP
GTP
UTP
NADH
UDP
NAD
CoA
TPNH
Spermine
DEL (Allosteric regulator) to Metabolic reaction
PS
PE
PI
SM
Cholesterol
Cer
DG
DEM (Substrate or product) to Metabolic reaction
Number of substrate or product regulations
ATP
ADP
TPN
NAD
TPNH
NADH
CoA
UDP
Glutamate
SAH
DEL (Substrate or product) to Metabolic reaction
Cer
PI
DG
PE
TG
PS
Cholesterol
SM
Fatty acid
B
Metabolic reactions activated in ob/ob mice
Metabolic reaction activated in ob/ob by Enzyme protein
Carbohydrate
Amino acid
Nucleotide
Lipid
FA syn.
FA elong.
BCAA degra.
TCA cycle
FA degra.
Metabolic reaction activated in ob/ob by Metabolite (substrate or product)
Metabolic reactions inhibited in ob/ob mice
Metabolic reaction inhibited in ob/ob by Enzyme protein
FA elong.

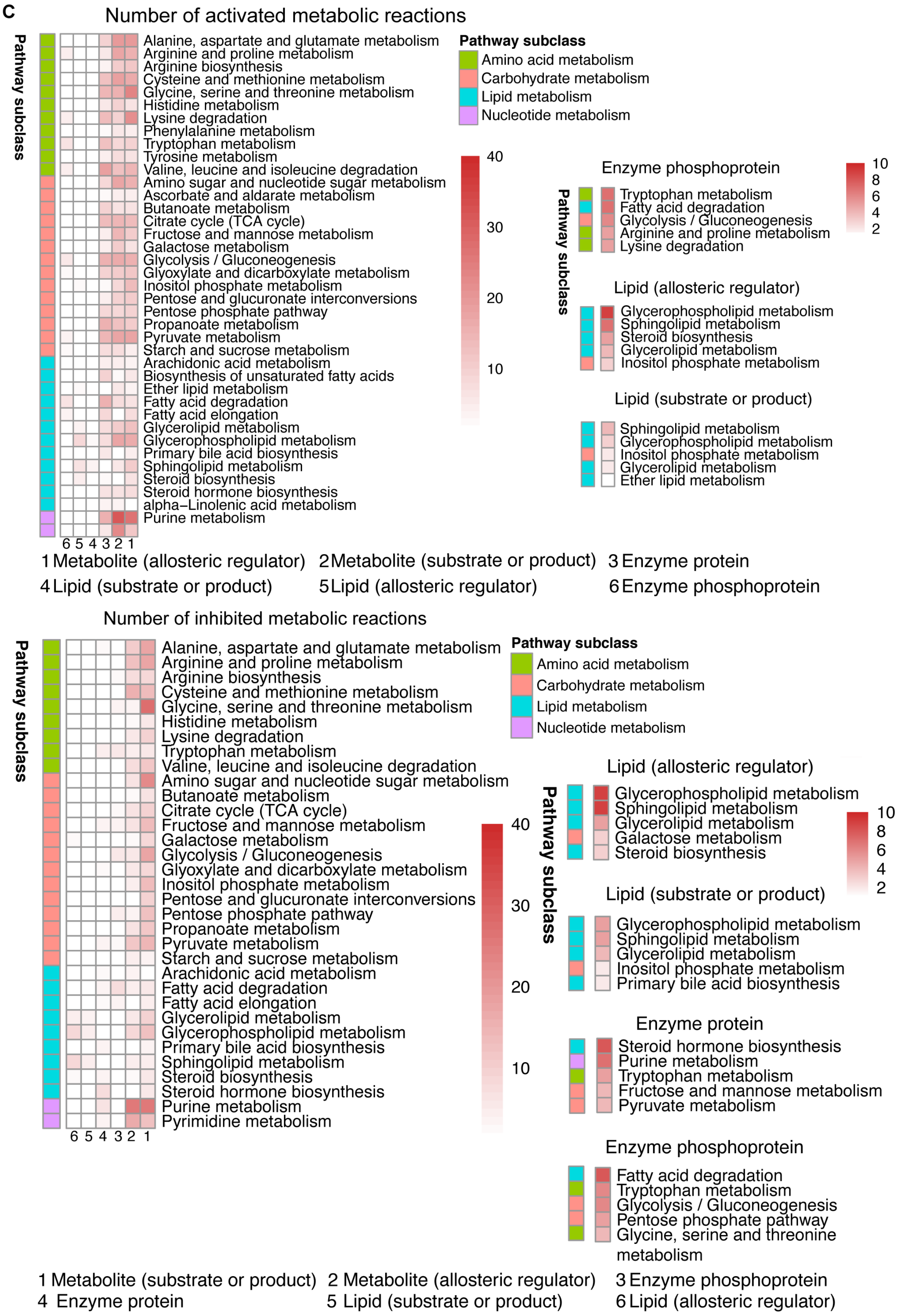
C
Number of activated metabolic reactions
Pathway subclass
Alanine, aspartate and glutamate metabolism
Arginine and proline metabolism
Arginine biosynthesis
Cysteine and methionine metabolism
Glycine, serine and threonine metabolism
Histidine metabolism
Lysine degradation
Phenylalanine metabolism
Tryptophan metabolism
Tyrosine metabolism
Valine, leucine and isoleucine degradation
Amino sugar and nucleotide sugar metabolism
Ascorbate and aldarate metabolism
Butanoate metabolism
Citrate cycle (TCA cycle)
Fructose and mannose metabolism
Galactose metabolism
Glycolysis / Gluconeogenesis
Glyoxylate and dicarboxylate metabolism
Inositol phosphate metabolism
Pentose and glucuronate interconversions
Pentose phosphate pathway
Propanoate metabolism
Pyruvate metabolism
Starch and sucrose metabolism
Arachidonic acid metabolism
Biosynthesis of unsaturated fatty acids
Ether lipid metabolism
Fatty acid degradation
Fatty acid elongation
Glycerolipid metabolism
Glycerophospholipid metabolism
Primary bile acid biosynthesis
Sphingolipid metabolism
Steroid biosynthesis
Steroid hormone biosynthesis
alpha–Linolenic acid metabolism
Purine metabolism
6 5 4 3 2 1
Pathway subclass
Amino acid metabolism
Carbohydrate metabolism
Lipid metabolism
Nucleotide metabolism
40
30
20
10
Enzyme phosphoprotein
Tryptophan metabolism
Fatty acid degradation
Glycolysis / Gluconeogenesis
Arginine and proline metabolism
Lysine degradation
10
8
6
4
2
Lipid (allosteric regulator)
Glycerophospholipid metabolism
Sphingolipid metabolism
Steroid biosynthesis
Glycerolipid metabolism
Inositol phosphate metabolism
Lipid (substrate or product)
Sphingolipid metabolism
Glycerophospholipid metabolism
Inositol phosphate metabolism
Glycerolipid metabolism
Ether lipid metabolism
1 Metabolite (allosteric regulator)
2 Metabolite (substrate or product)
3 Enzyme protein
4 Lipid (substrate or product)
5 Lipid (allosteric regulator)
6 Enzyme phosphoprotein
Number of inhibited metabolic reactions
Pathway subclass
Alanine, aspartate and glutamate metabolism
Arginine and proline metabolism
Arginine biosynthesis
Cysteine and methionine metabolism
Glycine, serine and threonine metabolism
Histidine metabolism
Lysine degradation
Tryptophan metabolism
Valine, leucine and isoleucine degradation
Amino sugar and nucleotide sugar metabolism
Butanoate metabolism
Citrate cycle (TCA cycle)
Fructose and mannose metabolism
Galactose metabolism
Glycolysis / Gluconeogenesis
Glyoxylate and dicarboxylate metabolism
Inositol phosphate metabolism
Pentose and glucuronate interconversions
Pentose phosphate pathway
Propanoate metabolism
Pyruvate metabolism
Starch and sucrose metabolism
Arachidonic acid metabolism
Fatty acid degradation
Fatty acid elongation
Glycerolipid metabolism
Glycerophospholipid metabolism
Primary bile acid biosynthesis
Sphingolipid metabolism
Steroid biosynthesis
Steroid hormone biosynthesis
Purine metabolism
Pyrimidine metabolism
6 5 4 3 2 1
Pathway subclass
Amino acid metabolism
Carbohydrate metabolism
Lipid metabolism
Nucleotide metabolism
40
30
20
10
Lipid (allosteric regulator)
Glycerophospholipid metabolism
Sphingolipid metabolism
Glycerolipid metabolism
Galactose metabolism
Steroid biosynthesis
10
8
6
4
2
Lipid (substrate or product)
Glycerophospholipid metabolism
Sphingolipid metabolism
Glycerolipid metabolism
Inositol phosphate metabolism
Primary bile acid biosynthesis
Enzyme protein
Steroid hormone biosynthesis
Purine metabolism
Tryptophan metabolism
Fructose and mannose metabolism
Pyruvate metabolism
Enzyme phosphoprotein
Fatty acid degradation
Tryptophan metabolism
Glycolysis / Gluconeogenesis
Pentose phosphate pathway
Glycine, serine and threonine metabolism
1 Metabolite (substrate or product)
2 Metabolite (allosteric regulator)
3 Enzyme phosphoprotein
4 Enzyme protein
5 Lipid (substrate or product)
6 Lipid (allosteric regulator)

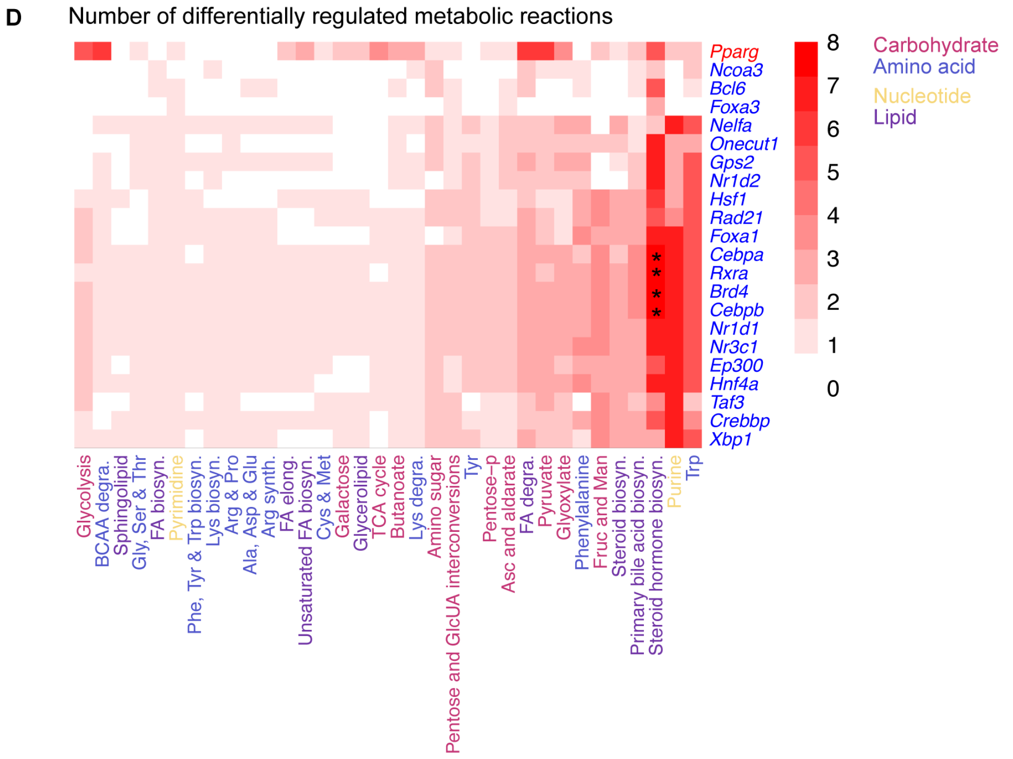


Figure S3

**Figure S3. Differential regulations analysis at *ad libitum* feeding. Related to Figure 3.**

(A) Bar plot of allosteric regulations by DEMs (top-left), allosteric regulations by DELs (top-right), regulations affected by DEMs functioned as substrates or products (bottom-left), regulations affected by DELs functioned as substrates or products (bottom-right). Red, activating regulations in *ob*/*ob* mice; blue, inhibiting regulations in *ob*/*ob* mice.

(B) The number of activated or inhibited metabolic reactions in *ob*/*ob* mice in each metabolic pathway regulated by metabolites as substrate or product. For each metabolic pathway (circle), the percentage of differentially regulated metabolic reactions mediated by either metabolites as substrates or products ($x$-axis) or enzyme proteins ($y$-axis). Left, activated metabolic reactions in *ob*/*ob* mice; right, inhibited metabolic reactions in *ob*/*ob* mice. The size of circles indicates the sum of percentages of differentially regulated metabolic reactions mediated by metabolites (substrate or products) and enzyme proteins. The color of the dots indicates the classes of metabolic pathways by the KEGG database. The dashed line is diagonal line.

(C) The number of activated metabolic reactions (top) and inhibited metabolic reactions (bottom). The top 5 dysregulated metabolic pathways on number scale regulated by enzyme phosphoproteins, lipids (allosteric regulator), lipids (substrate or product), and enzyme protein (only inhibited) were shown on the right panel.

(D) For each metabolic pathway (row), the number of differentially regulated metabolic reactions by Differentially Regulated TFs (DRTFs) (column). The color of different metabolic pathways (row) indicated the classes of metabolic pathways recorded by the KEGG database. The color of DRTFs indicated increased (red) or decreased (blue) DRTFs in *ob*/*ob* mice. The saturation of red in heatmap indicated the number of differentially regulated metabolic reactions. Hierarchical clustering analysis was performed using Euclidean distance and Ward's method. The sign * indicates differentially regulated metabolic reactions in metabolic pathway were significantly associated with DRTFs by one-tailed Fisher's exact test ($q < 0.01$).

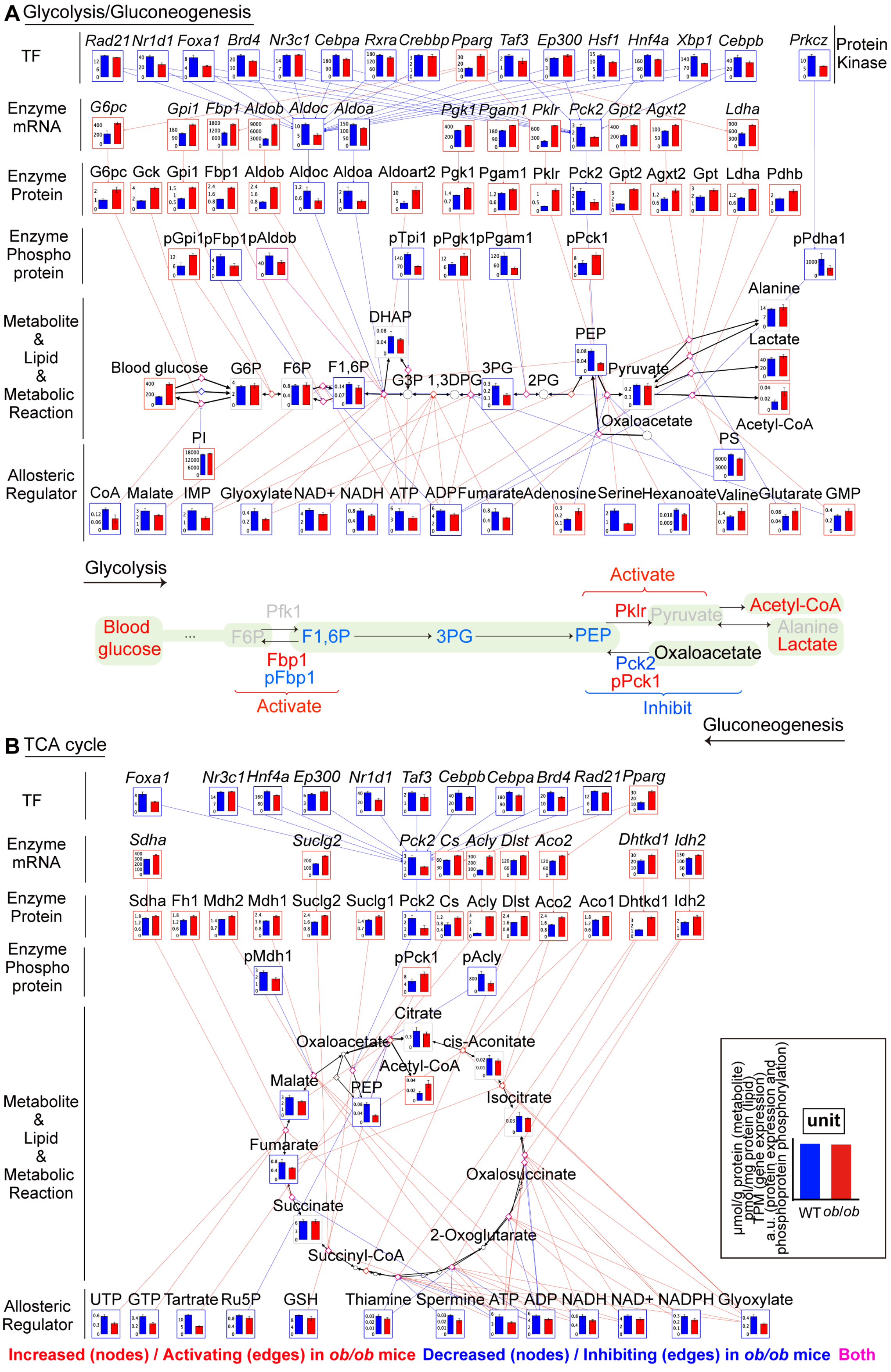
A Glycolysis/Gluconeogenesis
TF
Rad21 Nr1d1 Foxa1 Brd4 Nr3c1 Cebpa Rxra Crebbp Pparg Taf3 Ep300 Hsf1 Hnf4a Xbp1 Cebpb
Prkcz
Protein Kinase
Enzyme mRNA
G6pc Gpi1 Fbp1 Aldob Aldoc Aldoa Pgk1 Pgam1 Pklr Pck2 Gpt2 Agxt2 Ldha
Enzyme Protein
G6pc Gck Gpi1 Fbp1 Aldob Aldoc Aldoa Aldoart2 Pgk1 Pgam1 Pklr Pck2 Gpt2 Agxt2 Gpt Ldha Pdhb
Enzyme Phospho protein
pGpi1 pFbp1 pAldob pTpi1 pPgk1 pPgam1 pPck1 pPdha1
Metabolite & Lipid & Metabolic Reaction
DHAP PEP Alanine Lactate Blood glucose G6P F6P F1,6P G3P 1,3DPG 3PG 2PG Pyruvate Oxaloacetate Acetyl-CoA PS
PI
Allosteric Regulator
CoA Malate IMP Glyoxylate NAD+ NADH ATP ADP Fumarate Adenosine Serine Hexanoate Valine Glutarate GMP
Glycolysis
Activate
Pfk1
Pklr Pyruvate
Acetyl-CoA
Blood glucose
F6P F1,6P 3PG PEP
Alanine
Lactate
Oxaloacetate
Fbp1
pFbp1
Activate
Pck2
pPck1
Inhibit
Gluconeogenesis
B TCA cycle
TF
Foxa1 Nr3c1 Hnf4a Ep300 Nr1d1 Taf3 Cebpb Cebpa Brd4 Rad21 Pparg
Enzyme mRNA
Sdha Suclg2 Pck2 Cs Acly Dlst Aco2 Dhtkd1 Idh2
Enzyme Protein
Sdha Fh1 Mdh2 Mdh1 Suclg2 Suclg1 Pck2 Cs Acly Dlst Aco2 Aco1 Dhtkd1 Idh2
Enzyme Phospho protein
pMdh1 pPck1 pAcly
Metabolite & Lipid & Metabolic Reaction
Citrate Oxaloacetate cis-Aconitate Acetyl-CoA Malate PEP Isocitrate Fumarate Oxalosuccinate Succinate 2-Oxoglutarate Succinyl-CoA
μmol/g protein (metabolite)
pmol/mg protein (lipid)
TPM (gene expression)
a.u. (protein expression and phosphoprotein phosphorylation)
unit
WT ob/ob
Allosteric Regulator
UTP GTP Tartrate Ru5P GSH Thiamine Spermine ATP ADP NADH NAD+ NADPH Glyoxylate
Increased (nodes) / Activating (edges) in ob/ob mice Decreased (nodes) / Inhibiting (edges) in ob/ob mice Both

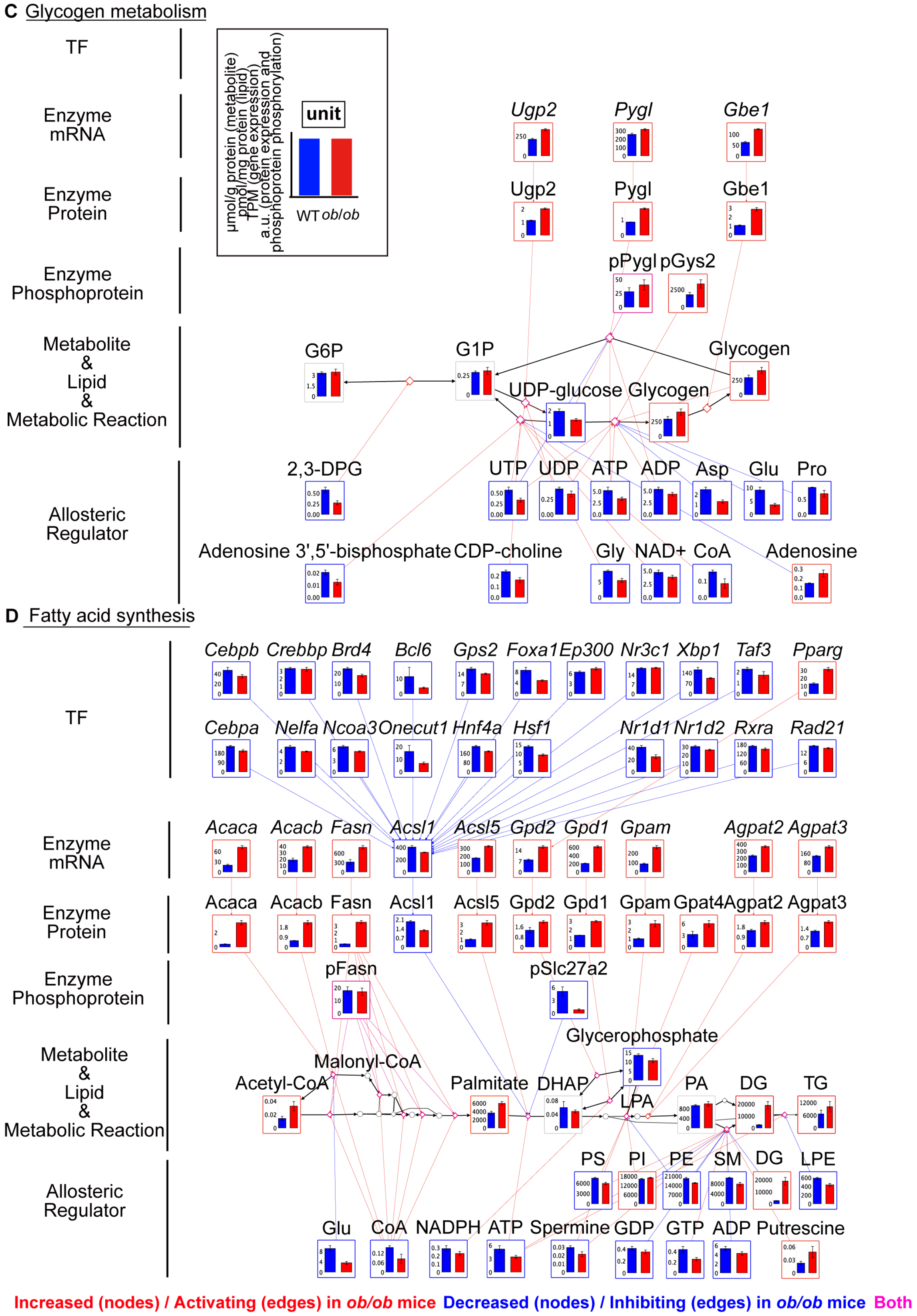
C Glycogen metabolism
TF
Enzyme mRNA
Enzyme Protein
Enzyme Phosphoprotein
Metabolite & Lipid & Metabolic Reaction
Allosteric Regulator
μmol/g protein (metabolite)
pmol/mg protein (lipid)
TPM (gene expression)
a.u. (protein expression and phosphoprotein phosphorylation)
unit
WT ob/ob
Ugp2
Pygl
Gbe1
pPygl
pGys2
G6P
G1P
Glycogen
UDP-glucose
2,3-DPG
UTP
UDP
ATP
ADP
Asp
Glu
Pro
Adenosine 3',5'-bisphosphate
CDP-choline
Gly
NAD+
CoA
Adenosine
D Fatty acid synthesis
Cebpb
Crebbp
Brd4
Bcl6
Gps2
Foxa1
Ep300
Nr3c1
Xbp1
Taf3
Pparg
Cebpa
Nelfa
Ncoa3
Onecut1
Hnf4a
Hsf1
Nr1d1
Nr1d2
Rxra
Rad21
Acaca
Acacb
Fasn
Acsl1
Acsl5
Gpd2
Gpd1
Gpam
Agpat2
Agpat3
Gpat4
pFasn
pSlc27a2
Glycerophosphate
Malonyl-CoA
Acetyl-CoA
Palmitate
DHAP
LPA
PA
DG
TG
PS
PI
PE
SM
LPE
Glu
CoA
NADPH
ATP
Spermine
GDP
GTP
ADP
Putrescine
Increased (nodes) / Activating (edges) in ob/ob mice
Decreased (nodes) / Inhibiting (edges) in ob/ob mice
Both

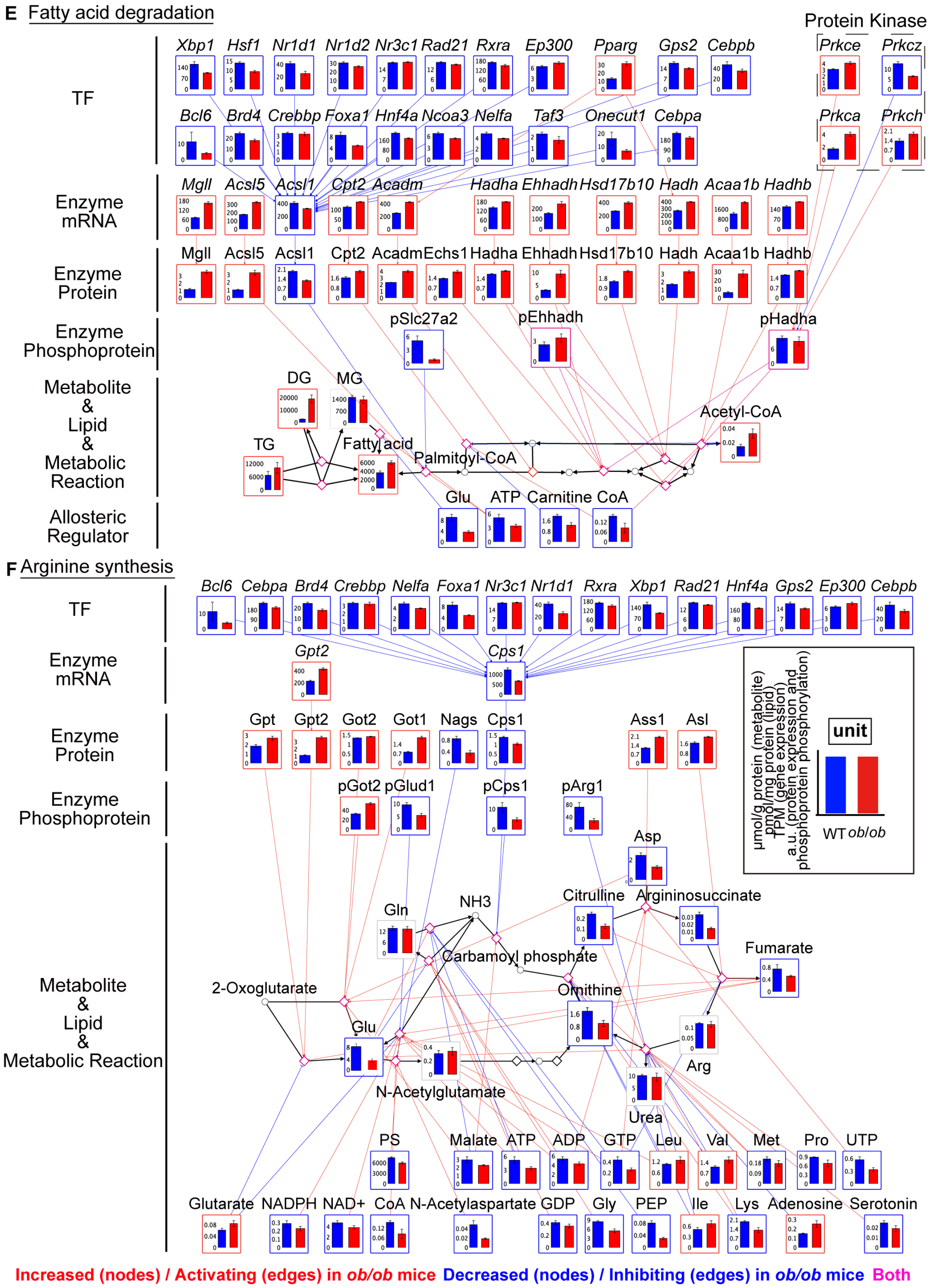
E Fatty acid degradation
TF
Xbp1 Hsf1 Nr1d1 Nr1d2 Nr3c1 Rad21 Rxra Ep300 Pparg Gps2 Cebpb
Bcl6 Brd4 Crebbp Foxa1 Hnf4a Ncoa3 Nelfa Taf3 Onecut1 Cebpa
Protein Kinase
Prkce Prkcz
Prkca Prkch
Enzyme mRNA
Mgll Acsl5 Acsl1 Cpt2 Acadm Hadha Ehhadh Hsd17b10 Hadh Acaa1b Hadhb
Enzyme Protein
Mgll Acsl5 Acsl1 Cpt2 Acadm Echs1 Hadha Ehhadh Hsd17b10 Hadh Acaa1b Hadhb
Enzyme Phosphoprotein
pSlc27a2 pEhhadh pHadha
Metabolite & Lipid & Metabolic Reaction
DG MG TG Fatty acid Palmitoyl-CoA Acetyl-CoA
Allosteric Regulator
Glu ATP Carnitine CoA
F Arginine synthesis
TF
Bcl6 Cebpa Brd4 Crebbp Nelfa Foxa1 Nr3c1 Nr1d1 Rxra Xbp1 Rad21 Hnf4a Gps2 Ep300 Cebpb
Enzyme mRNA
Gpt2 Cps1
Enzyme Protein
Gpt Gpt2 Got2 Got1 Nags Cps1 Ass1 Asl
Enzyme Phosphoprotein
pGot2 pGlud1 pCps1 pArg1
Metabolite & Lipid & Metabolic Reaction
Asp Citrulline Argininosuccinate NH3 Gln Carbamoyl phosphate Fumarate 2-Oxoglutarate Ornithine Glu Arg N-Acetylglutamate Urea
PS Malate ATP ADP GTP Leu Val Met Pro UTP
Glutarate NADPH NAD+ CoA N-Acetylaspartate GDP Gly PEP Ile Lys Adenosine Serotonin
unit
μmol/g protein (metabolite)
pmol/mg protein (lipid)
TPM (gene expression)
a.u. (protein expression and phosphoprotein phosphorylation)
WT ob/ob
Increased (nodes) / Activating (edges) in ob/ob mice Decreased (nodes) / Inhibiting (edges) in ob/ob mice Both

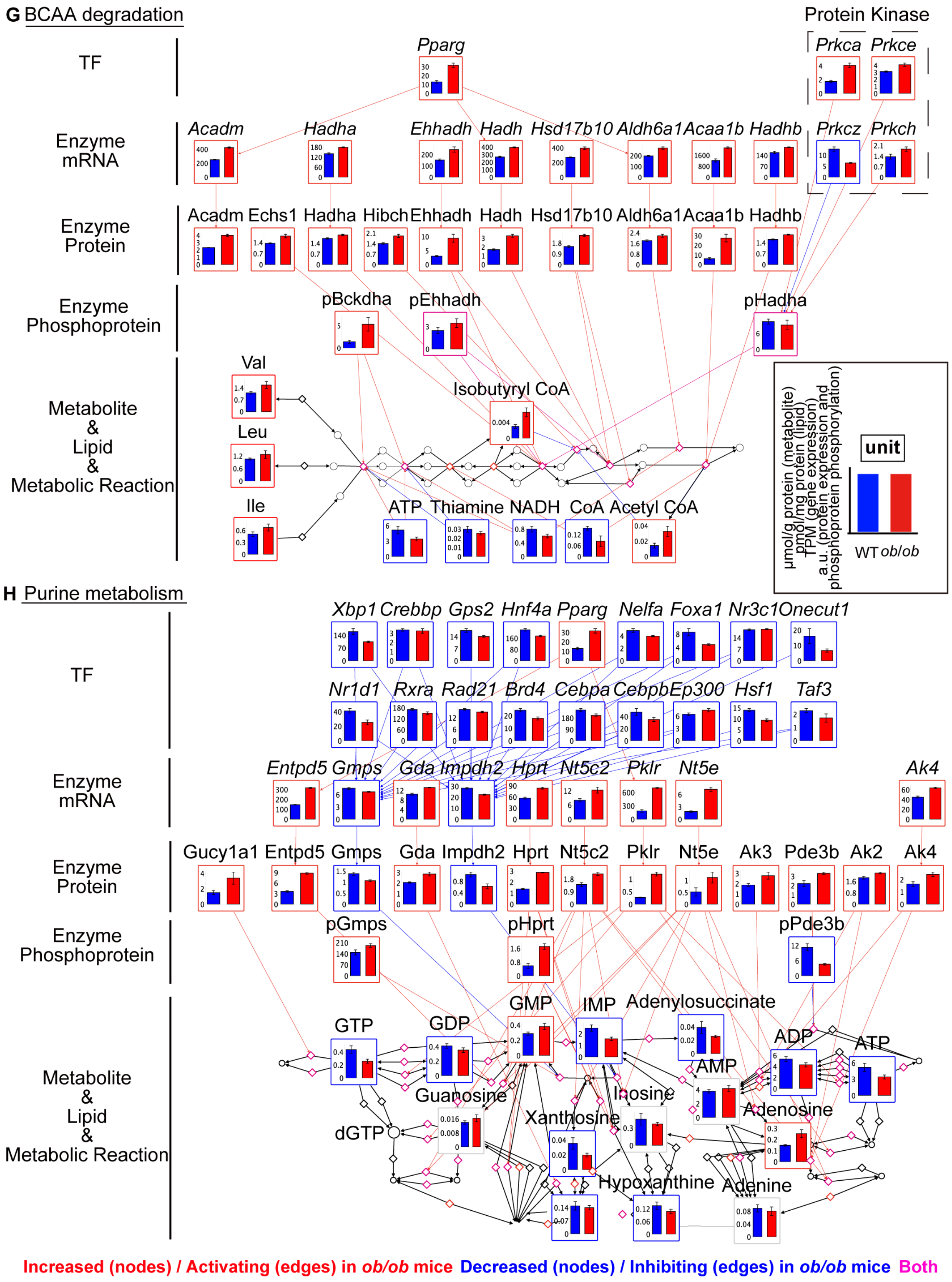
G BCAA degradation
Protein Kinase
TF
Enzyme mRNA
Enzyme Protein
Enzyme Phosphoprotein
Metabolite & Lipid & Metabolic Reaction
Pparg
Prkca Prkce
Acadm Hadha Ehhadh Hadh Hsd17b10 Aldh6a1 Acaa1b Hadhb Prkcz Prkch
Acadm Echs1 Hadha Hibch Ehhadh Hadh Hsd17b10 Aldh6a1 Acaa1b Hadhb
pBckdha pEhhadh pHadha
Val
Leu
Ile
Isobutyryl CoA
ATP Thiamine NADH CoA Acetyl CoA
unit
μmol/g protein (metabolite)
pmol/mg protein (lipid)
TPM (gene expression)
a.u. (protein expression and phosphoprotein phosphorylation)
WT ob/ob
H Purine metabolism
TF
Enzyme mRNA
Enzyme Protein
Enzyme Phosphoprotein
Metabolite & Lipid & Metabolic Reaction
Xbp1 Crebbp Gps2 Hnf4a Pparg Nelfa Foxa1 Nr3c1 Onecut1
Nr1d1 Rxra Rad21 Brd4 Cebpa Cebpb Ep300 Hsf1 Taf3
Entpd5 Gmps Gda Impdh2 Hprt Nt5c2 Pklr Nt5e Ak4
Gucy1a1 Entpd5 Gmps Gda Impdh2 Hprt Nt5c2 Pklr Nt5e Ak3 Pde3b Ak2 Ak4
pGmps pHprt pPde3b
GTP GDP GMP IMP Adenylosuccinate ADP ATP
AMP
Guanosine Inosine Adenosine
Xanthosine
dGTP
Hypoxanthine Adenine
Increased (nodes) / Activating (edges) in ob/ob mice Decreased (nodes) / Inhibiting (edges) in ob/ob mice Both

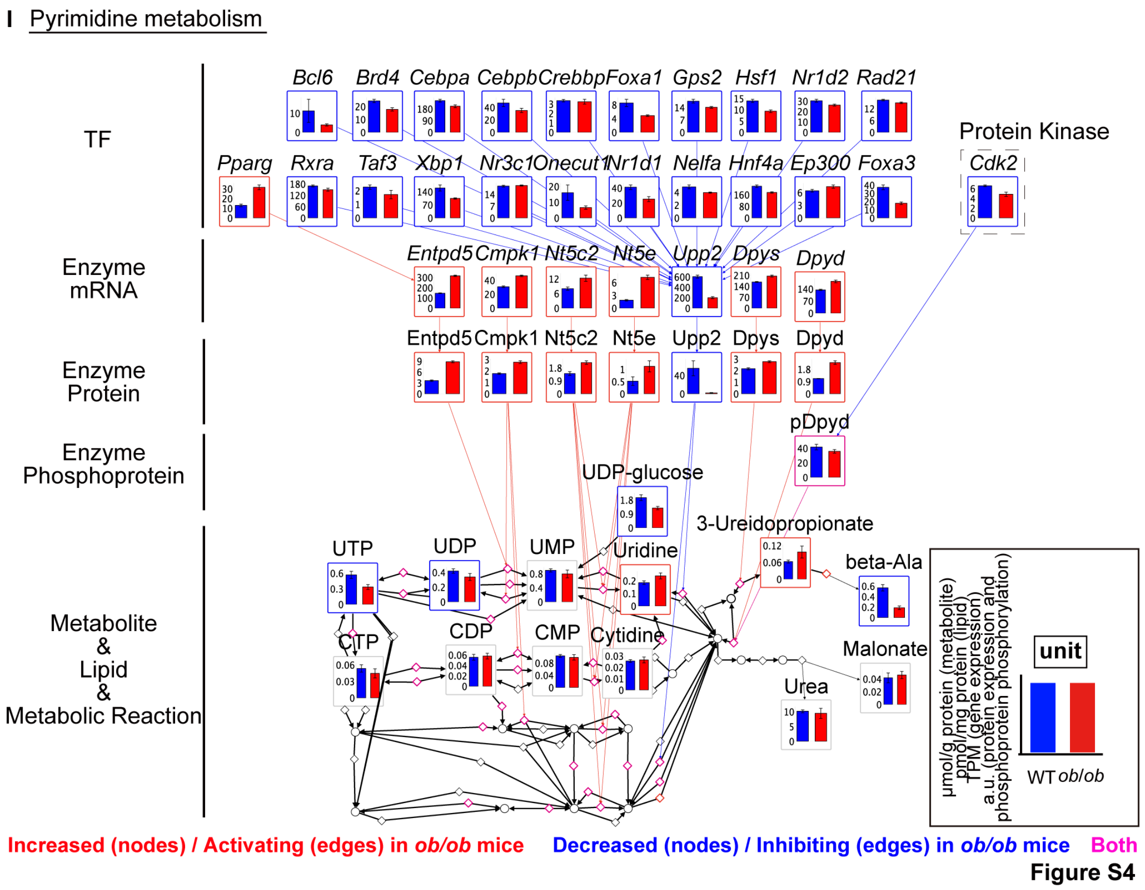


**Figure S4. Metabolic reactions and molecules in metabolic pathways in WT and *ob*/*ob* mice at *ad libitum* feeding. Related to Figure 4.**

(A) Glycolysis/Gluconeogenesis (top). Schematic plot of glycolysis/gluconeogenesis including only rate-limiting enzymes and metabolites information (bottom). The information was obtained from “Glycolysis and Gluconeogenesis” (mmu00010) in the KEGG database. For glycolysis/gluconeogenesis, we found that blood glucose, lactate, and acetyl-CoA increased, whereas the glycolytic intermediate metabolites in intracellular, Fructose 1,6-bisphosphate (F1,6P), 3-Phospho-D-glycerate (3PG), Phosphoenolpyruvate (PEP) decreased in *ob*/*ob* mice. The rate-limiting enzymes of glycolysis Pklr increased, suggesting that glycolysis from PEP to pyruvate is activated. While the rate-limiting enzymes of gluconeogenesis, Fbp1 increased, pFbp1 decreased. Phosphorylation reduced the activity of Fbp1, and such inhibition decreased[11]. Pck2 decreased, and pPck1 increased. Increased Gsk3β mediates Pck1 phosphorylation and promotes Pck1 degradation at high-energy conditions[12]. Many enzyme proteins increased in *ob*/*ob* mice (82%; 14 proteins of total 17 proteins), 4 of which are regulated by the TF *Pparg*. 3 enzyme proteins decreased, which are regulated by 14 TFs, including *Foxa1*, *Cebpb*, and *Xbp1*. Decrease of protein kinase *Prkcz* associated with decreased phosphorylation of pPdha1. The expression or phosphorylation of metabolic enzymes increased for overall glycolysis/gluconeogenesis pathways, including Gpi1, Pgk1, Ldha, Pklr, and Pgam1. The number of activating allosteric regulations (26 regulations) is larger than that of inhibiting allosteric regulations (10 regulations). It is likely that the upstream of gluconeogenesis, from oxaloacetate to PEP, is inhibited, whereas that of the downstream of gluconeogenesis, from F1,6P to F6P, and the downstream of glycolysis, from PEP to pyruvate, is activated by enzymes Fbp1, pFbp1, and Pklr in *ob*/*ob* mice.

(B) TCA cycle. The information was obtained from “Citrate cycle” (mmu00020) in the KEGG database. For TCA cycle, we found that acetyl-CoA increased, while intermediate metabolites, malate, fumarate, and PEP, decreased in *ob*/*ob* mice. Acly, Aco2, Cs, Dhtkd1, Idh2, Aco1, Mdh2, Sdha, Fh1, and Mdh1, involved in TCA cycle and increased. Pck2 regulated by 10 TFs decreased. pPck1 increases phosphorylated, pMdh1 and pAcly decrease phosphorylated in *ob*/*ob* mice. Among enzyme protein regulations, most of them are the activating regulations (17 regulations) by 13 increased enzyme proteins in *ob*/*ob* mice. 15 decreased allosteric regulators, including GTP, UTP, ATP, $NAD^+$, NADH, and glyoxylate, are activating metabolic reactions. These results suggest that metabolic regulations by enzyme proteins and allosteric regulators are likely to activate TCA cycle, whereas intermediate metabolites, PEP, malate, and fumarate, decreased in *ob*/*ob* mice.

(C) Glycogen metabolism. The information was obtained from “starch and sucrose metabolism” (mmu00500) in the KEGG database. For glycogen metabolism, we found that glycogen increased, and UDP-glucose decreased in *ob*/*ob* mice. G6P and G1P did not change. Enzyme proteins, Ugp2, Pygl, and Gbe1 increased, and enzyme phosphoproteins pGys2 increased. 11 decreased allosteric regulators, including ATP, UTP, ADP, and CoA, causing 17 activating allosteric regulations. Glycogen synthesis may decrease with decreased substrate in *ob*/*ob* mice, UDP-glucose.

(D) Fatty acid synthesis. The information was obtained from "fatty acid biosynthesis" (mmu00061), "glycerolipid metabolism" (mmu00561), and "glycerophospholipid metabolism" (mmu00564) in the KEGG database.

(E) Fatty acid degradation. The information was obtained from "fatty acid degradation" (mmu00071) and "glycerolipid metabolism" (mmu00561) in the KEGG database. We identified a significant increase in lipid diacylglycerol (DG) ($\log_2$ FC = 2.8, q = 0.009), TG precursor in *ob*/*ob* mice. TG, fatty acid, and acetyl-CoA also increased. We identified 10 increased enzyme proteins in fatty acid synthesis, and 11 increased enzyme proteins in fatty acid degradation. Enzyme protein Acsl1, which regulated by 20 TFs, decreased in *ob*/*ob* mice. In fatty acid degradation, protein kinases *Prkca*, *Prkch*, *Prkcz*, and *Prkce* modified substrate pHadha by adding phosphates. The enzyme phosphoprotein, pSlc27a2, decreased, and pFasn, pEhhadh, pHadha increased and decreased. These results suggest that fatty acid synthesis is activated by enzyme proteins and allosteric regulators, and fatty acid degradation is activated by enzyme proteins in *ob*/*ob* mice.

(F) Arginine synthesis. The information was obtained from "arginine biosynthesis" (mmu00220) in the KEGG database. Urea cycle is the main step for synthesizing Arg. All the 6 identified DEMs in arginine synthesis pathway, including ornithine, Asp, and argininosuccinate, decreased in *ob*/*ob* mice. Citrulline, a precursor of Arg and limiting factor for de novo Arg production, decreased. We found that the expression of urea cycle related metabolic enzyme, Cps1, regulated by 15 TFs showed decreased expression and phosphorylation, which is consistent with the previous study[13,14]. 6 enzyme proteins, including Gpt, Gpt2, Got1, and Got2 increased. The phosphorylation of metabolic enzymes pCps1, pGlud1, and pArg1 decreased. 22 decreased metabolites leading to 36 activating regulations.

(G) BCAA degradation. The information was obtained from "valine, leucine and isoleucine degradation" (mmu00280) in the KEGG database. We found all 6 identified DEMs, including three BCAAs, valine, leucine, and isoleucine, isobutyryl-CoA, and acetyl-CoA increased in *ob*/*ob* mice. 10 enzyme proteins, 3 enzyme phosphoproteins increased. Most of enzyme proteins are regulated by the responding enzyme mRNAs. Increased protein kinases *Prkca*, *Prkce*, and *Prkch* are associated with increased enzyme phosphoprotein pHadha. 4 allosteric regulators, including ATP, CoA, and NADH, decreased. 7 activating and 5 inhibiting allosteric regulations in *ob*/*ob* mice.

(H) Purine metabolism. The information was obtained from "purine metabolism" (mmu00230) in the KEGG database.

(I) Pyrimidine metabolism. The information was obtained from "pyrimidine metabolism" (mmu00240) in the KEGG database. In purine metabolism, metabolites including ATP, ADP, GTP, GDP, IMP, and adenylosuccinate, decreased in *ob*/*ob* mice. GMP and adenosine increased in *ob*/*ob* mice. Enzyme proteins Entpd5 and Pklr increased, which were regulated by *Pparg*. Gmps and Impdh2, two important enzyme proteins in GMP synthesis, decreased, which were regulated by 17 TFs. Enzyme phosphoproteins, pGmps and pHprt increased, whereas pPde3b decreased. In pyrimidine metabolism, we found decrease in metabolites UTP, UDP, UDP-Glucose, beta-Alanine, and increase in metabolites uridine and 3-Ureidopropionate in *ob*/*ob* mice. 20 decreased TFs regulated decreased expression of Upp2. Decreased protein kinase *Cdk2* regulated enzyme phosphoprotein pDpyd. The expression of enzyme proteins, including Entpd5, Nt5e, Nt5c2, and Dpys increased. In conclusion, we found a dynamic decrease of nucleotide in *ob*/*ob* mice, including triphosphate with three phosphate groups attached to ribose, ATP, GTP, and UTP. The Metabolites and Lipids as Allosteric Regulators were combined into the same region at different heights for better visualization. Bar plots were shown on the corresponding nodes as the means and SEMs of replicates in WT and *ob*/*ob* mice. For metabolites and lipids as substrates or products in metabolic pathway, we showed for all measured molecules. The colors of frames and edges indicate the changed direction.

Diamond nodes indicate metabolic reactions. Black circles indicate the metabolites were not measured. From metabolite to metabolic reaction, only allosteric regulations are colored. Only regulations connected with differentially expressed or phosphorylated molecules remained. Red, increased molecules or activating regulations in *ob*/*ob* mice; blue, decreased molecules or inhibiting regulations in *ob*/*ob* mice; magenta, both increased and decreased molecules, activated and inhibited metabolic reactions, activating and inhibiting regulations in *ob*/*ob* mice; gray, unchanged molecules between WT and *ob*/*ob* mice.

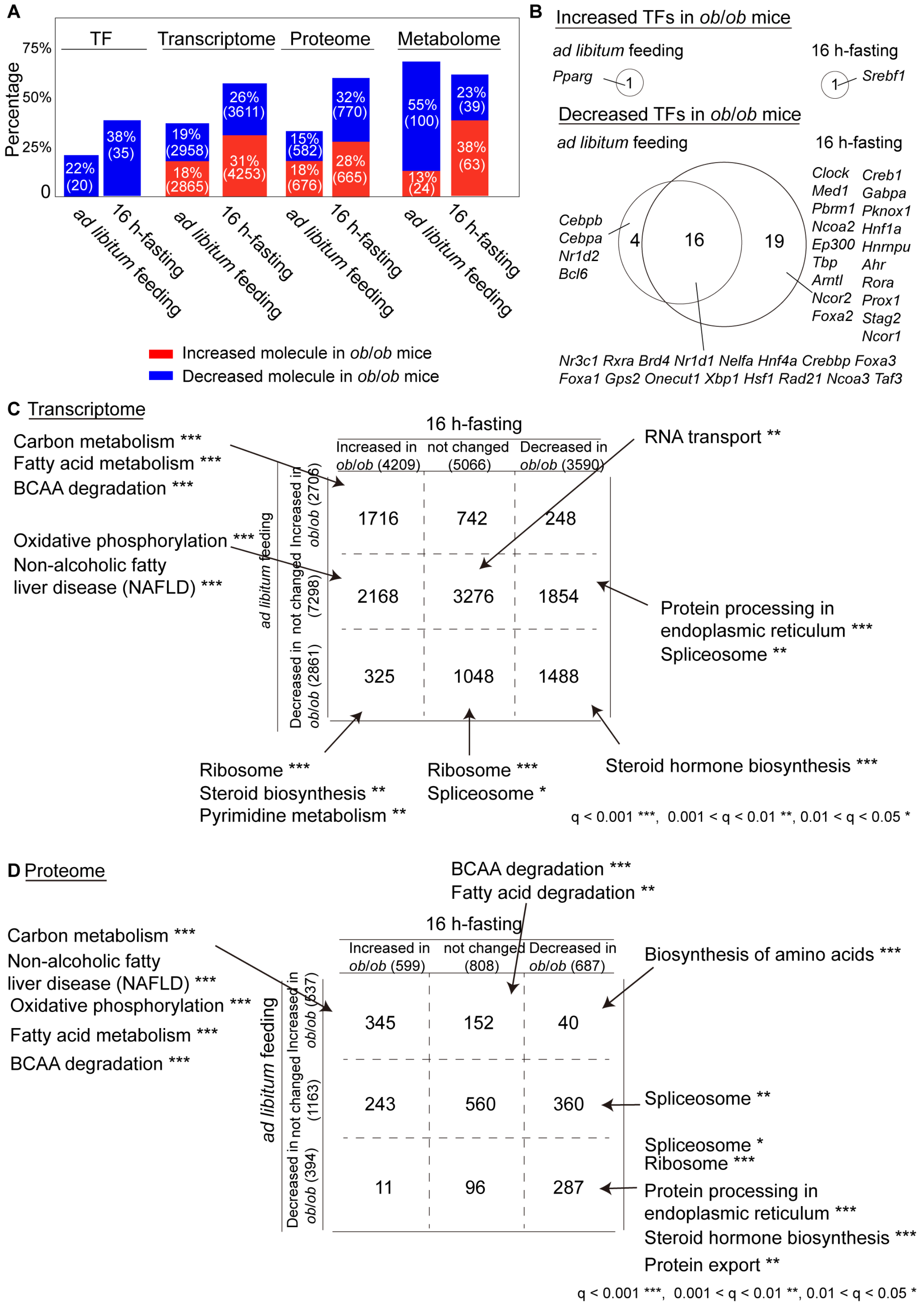
A
TF
Transcriptome
Proteome
Metabolome
Percentage
75%
50%
25%
0
22% (20)
38% (35)
19% (2958)
18% (2865)
26% (3611)
31% (4253)
15% (582)
18% (676)
32% (770)
28% (665)
55% (100)
13% (24)
23% (39)
38% (63)
ad libitum feeding
16 h-fasting
Increased molecule in ob/ob mice
Decreased molecule in ob/ob mice
B
Increased TFs in ob/ob mice
ad libitum feeding
16 h-fasting
Pparg
1
1
Srebf1
Decreased TFs in ob/ob mice
ad libitum feeding
16 h-fasting
Cebpb Cebpa Nr1d2 Bcl6
4
16
19
Clock Med1 Pbrm1 Ncoa2 Ep300 Tbp Arntl Ncor2 Foxa2
Creb1 Gabpa Pknox1 Hnf1a Hnrnpu Ahr Rora Prox1 Stag2 Ncor1
Nr3c1 Rxra Brd4 Nr1d1 Nelfa Hnf4a Crebbp Foxa3
Foxa1 Gps2 Onecut1 Xbp1 Hsf1 Rad21 Ncoa3 Taf3
C Transcriptome
16 h-fasting
Increased in ob/ob (4209)
not changed (5066)
Decreased in ob/ob (3590)
ad libitum feeding
Increased in ob/ob (2706)
not changed (7298)
Decreased in ob/ob (2861)
1716
742
248
2168
3276
1854
325
1048
1488
Carbon metabolism ***
Fatty acid metabolism ***
BCAA degradation ***
Oxidative phosphorylation ***
Non-alcoholic fatty liver disease (NAFLD) ***
RNA transport **
Protein processing in endoplasmic reticulum ***
Spliceosome **
Ribosome ***
Steroid biosynthesis **
Pyrimidine metabolism **
Ribosome ***
Spliceosome *
Steroid hormone biosynthesis ***
q < 0.001 ***, 0.001 < q < 0.01 **, 0.01 < q < 0.05 *
D Proteome
BCAA degradation ***
Fatty acid degradation **
16 h-fasting
Increased in ob/ob (599)
not changed (808)
Decreased in ob/ob (687)
ad libitum feeding
Increased in ob/ob (537)
not changed (1163)
Decreased in ob/ob (394)
345
152
40
243
560
360
11
96
287
Carbon metabolism ***
Non-alcoholic fatty liver disease (NAFLD) ***
Oxidative phosphorylation ***
Fatty acid metabolism ***
BCAA degradation ***
Biosynthesis of amino acids ***
Spliceosome **
Spliceosome *
Ribosome ***
Protein processing in endoplasmic reticulum ***
Steroid hormone biosynthesis ***
Protein export **
q < 0.001 ***, 0.001 < q < 0.01 **, 0.01 < q < 0.05 *

**E** Metabolome

| *ad libitum* feeding \ 16 h-fasting | Increased in *ob/ob* (55) | not changed (61) | Decreased in *ob/ob* (35) |
|---|---|---|---|
| Increased in *ob/ob* (20) | Val, Pipecolate, Ile, Leu, Lactate, 1-Methylnicotinamide, Isethionate, Glycogen, gamma-Guanidinobutyrate, Adenosine 10 | Nicotinamide, Taurine, Uridine, Acetyl CoA, Glutarate, 3-Ureidopropionate, 3-Hydroxybutyrate, 3-Hydroxy-3-methylglutarate 8 | GMP, Isobutyryl CoA 2 |
| not changed (48) | Urea, Diethanolamine, Asn, Succinate, S7P, Adenine, Spermidine, Gln, cis-Aconitate, His, Phe, Arg, SAM+, ADP-ribose, Trp, Cytidine, Guanosine, G6P, F6P, Citrate, 2-Hydroxyglutarate, 6-Phosphogluconate, Mucate, CMP, CDP, Taurocholate, UDP-glucuronate, Glutathione, N1-Acetylspermidine, 29 | Ala, Creatinine, 3-Methylhistidine, 1-Methylhistamine, Glucosamine, Tyr, N6,N6,N6-Trimethyllysine, Azelate, o-Acetylcarnitine, Inosine, Malonate, Threonate, Terephthalate, N-Acetylleucine, Phosphorylcholine 15 | DHAP, N-Acetylglutamate, UMP, AMP 4 |
| Decreased in *ob/ob* (83) | Ser, Homoserine, Thr, Urate, 5-Methylthioadenosine, ADP, F1,6P, 3PG, PEP, Glyoxylate, N-Acetylaspartate, UDP, UTP ATP, GTP, NAD+ 16 | 4-Hydroxymethylimidazole, Spermine, Histamine, IMP, Anserine, Creatine, Asp, Lys, Met, Methionine sulfoxide, Gly-Leu, N-epsilon-Acetyllysine, Cystathionine, Ru5P, Thymidine, Pyridoxamine 5'-phosphate, GDP, Glycerophosphorylcholine, Thiamine, Fumarate, Malate, Thiamine monophosphate, 2,3-DPG, Hexanoate, NADP+, 5-Oxoproline, Ethanolamine phosphate, Glycolate, Glucuronate, Dodecanoate, 2-Deoxyribose 1-phosphate, Pantothenate, Glycerophosphate N-Acetylglucosamine 1-phosphate, FAD, UDP-glucose, Pelargonate, UDP-N-acetylglucosamine, 38 | Gly, beta-Ala, Sarcosine, GABA, Choline, Glu, 3-Aminoisobutyrate, N,N-Dimethylglycine, Hypotaurine, Pro, Betaine, Hydroxyproline, SAH, Ornithine, Hypoxanthine, gamma-Butyrobetaine, alpha-Aminoadipate, Guanidinosuccinate, N-Acetylhistidine, Carnitine, 2-Hydroxybutyrate, Tartrate, Adenylosuccinate, CDP-choline, 2AB, Citrulline, Adenosine 3',5'-diphosphate, Saccharopine, Argininosuccinate, 29 |

**F** Metabolite Class

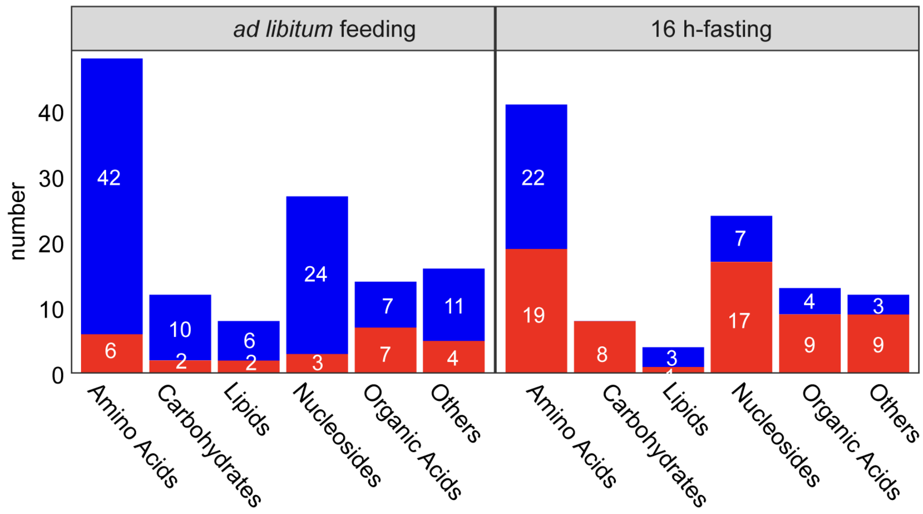


**Figure S5**

**Figure S5. Comparison on single omic data between *ad libitum* feeding and 16 h-**

**fasting in the liver of WT and *ob*/*ob* mice. Related to Figure 6A.**

(A) Percentages of increased and decreased TFs, mRNAs, proteins, and metabolites in *ob*/*ob* mice at *ad libitum* feeding and 16 h-fasting[1]. Number is shown insides parenthesis.

(B) Venn plot of increased (top) and decreased (bottom) TFs in *ob*/*ob* mice between *ad libitum* feeding and 16 h-fasting[1].

(C) KEGG enrichment analysis of mRNAs divided into 9 groups. Both increased, decreased, not changed in *ob*/*ob* mice at *ad libitum* feeding and 16 h-fasting[1]. Opposite changed in *ob*/*ob* mice between *ad libitum* feeding and 16 h-fasting. Changed only in *ob*/*ob* mice at *ad libitum* feeding or 16 h-fasting.

(D) KEGG enrichment analysis of proteins divided into 9 groups. The details are the same as (C).

(E) Metabolites in 9 groups at *ad libitum* feeding and 16 h-fasting[1]. The details are the same as (C).

(F) The DEMs classified by metabolite class at *ad libitum* feeding and 16 h-fasting[1] according to the KEGG database. We compared the differences of molecules in each omic layer (TF, mRNA, protein, metabolite) between *ad libitum* feeding and 16 h-fasting (Figure S5). The percentages of changed TFs, mRNAs, proteins, and metabolites were higher at 16 h-fasting than at *ad libitum* feeding, except for decreased metabolites that are higher at *ad libitum* feeding (Figure S5A). These phenomena suggest that obesity has a larger impact on overall trans-omic networks, except the metabolite layer, in the liver at 16h fasting than at *ad libitum* feeding. We directly compared molecules in each layer (Figure S5B-S5F). For increased TFs, only *Pparg* at *ad libitum* feeding and *Srebf1* at 16 h-fasting were predicted (Figure S5B). For the decreased TFs, most of them (16 TFs) intersected between *ad libitum* feeding and 16 h-fasting, including *Nr3c1*, *Nr1d1*, *Xbp1*. We divided mRNAs and proteins into 9 groups according to their changed direction and performed KEGG enrichment analysis on them (Figure S5C and S5D). For KEGG enrichment analysis on transcriptome (Figure S5C), Carbon metabolism, Fatty acid metabolism, and BCAA degradation increased at both *ad libitum* feeding and 16 h-fasting. Steroid hormone biosynthesis decreased at both *ad libitum* feeding and 16 h-fasting. Oxidative phosphorylation, Non-alcoholic fatty liver disease (NAFLD) increased, and Protein processing in endoplasmic reticulum decreased only at 16 h-fasting. KEGG enrichment analysis on proteome showed that Carbon metabolism and Non-alcoholic fatty liver disease increased at both *ad libitum* feeding and 16 h-fasting (Figure S5D). Ribosome, Spliceosome, Protein processing in endoplasmic reticulum, and Protein export decreased at both *ad libitum* feeding and 16 h-fasting. BCAA degradation and Fatty acid degradation increased only at *ad libitum* feeding. Transcriptome KEGG enrichment analysis results in principle consistent with proteome, suggesting that change of protein amounts is regulated by gene expression. For changed metabolites, F1,6P, 3PG, and PEP in glycolysis/gluconeogenesis, ADP, ATP, UDP, UTP, and GTP in purine and pyrimidine metabolism, increased at 16 h-fasting while decreased at *ad libitum* feeding (Figure S5E). More decreased Amino Acids, Carbohydrates, Lipids, Nucleosides, and Others metabolites than increased of them in *ob*/*ob* mice at *ad libitum* feeding. More increased Carbohydrates, Nucleosides, Organic Acids, and Others metabolites than decreased of them in *ob*/*ob* mice at 16 h-fasting (Figure S5F).

**A** Allosteric regulation (metabolite)

| *ad libitum* feeding \ 16 h-fasting | Inhibitor increased in *ob*/*ob* mice (498) | Inhibitor decreased in *ob*/*ob* mice (139) | Activator increased in *ob*/*ob* mice (164) | Activator decreased in *ob*/*ob* mice (27) |
|---|---|---|---|---|
| Inhibitor increased in *ob*/*ob* mice (85) | 59 | 25 | 1 | 0 |
| Inhibitor decreased in *ob*/*ob* mice (552) | 400 | 111 | 38 | 3 |
| Activator increased in *ob*/*ob* mice (17) | 1 | 0 | 14 | 2 |
| Activator decreased in *ob*/*ob* mice (174) | 38 | 3 | 111 | 22 |

**B** Substrate/Product regulation (metabolite)

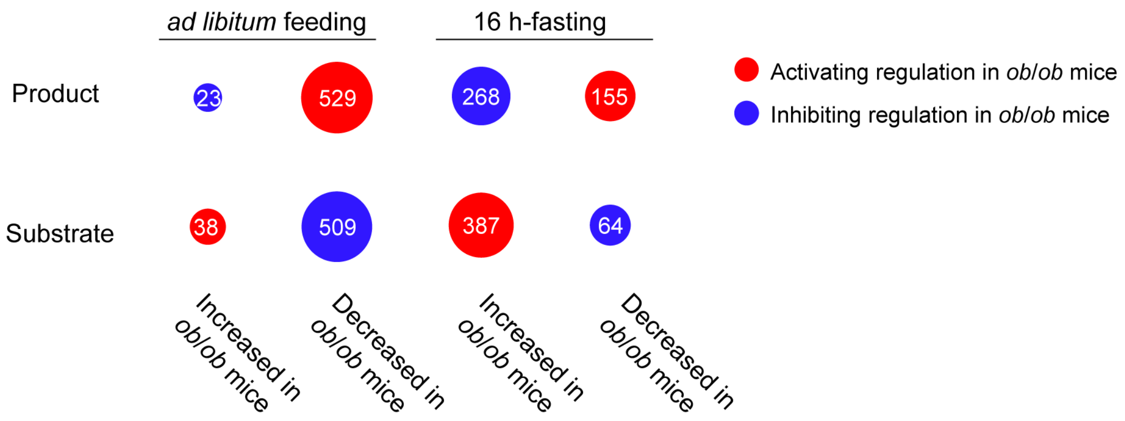

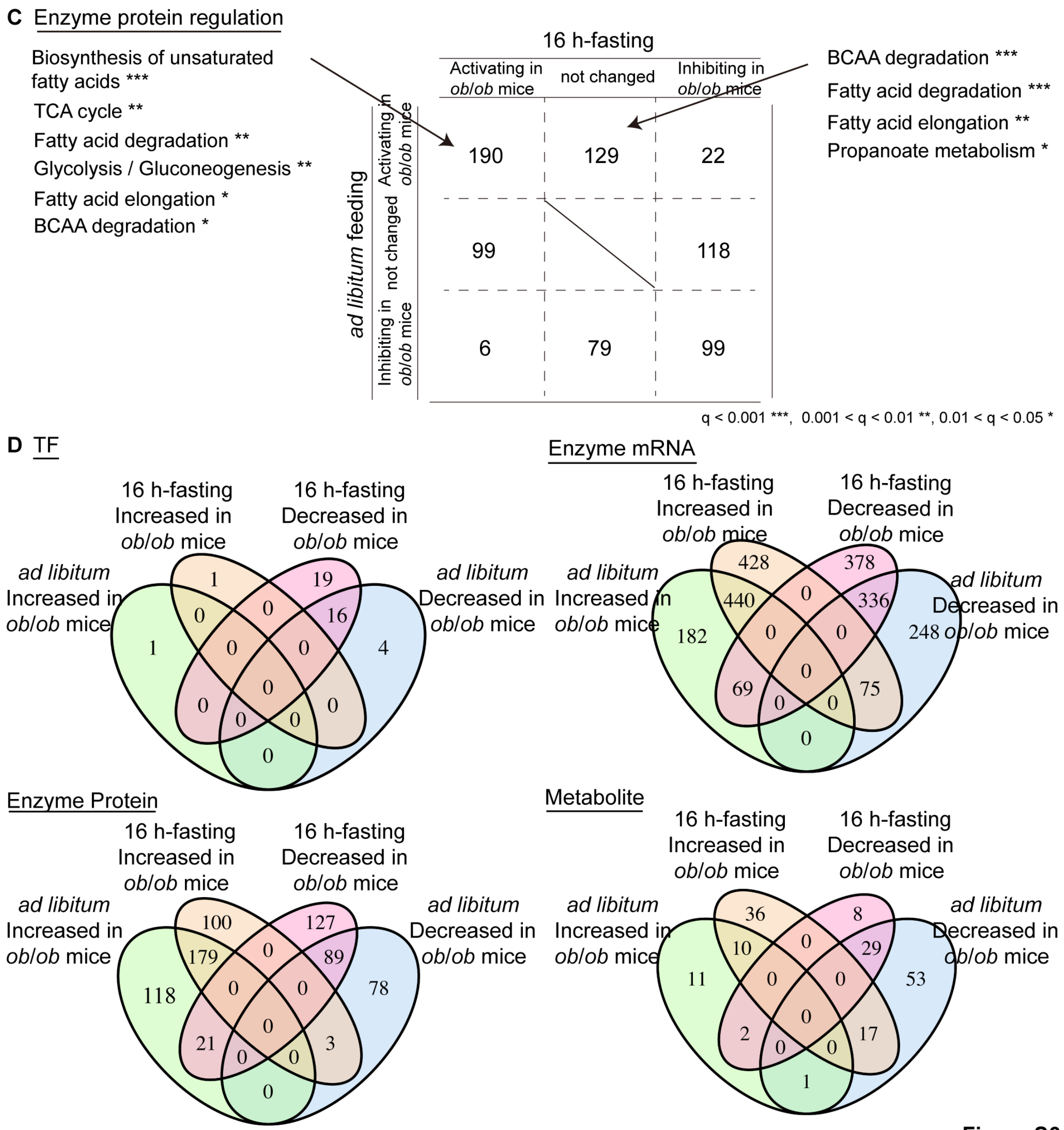


Figure S6

**Figure S6. Comparison on differential regulations and trans-omic networks between *ad libitum* feeding and 16 h-fasting in the liver of WT and *ob*/*ob* mice. Related to Figure 7A.**

(A) The numbers of allosteric regulations by metabolites at *ad libitum* feeding and 16 h-fasting[1] divided into 16 groups by inhibitor or activator increased or decreased in *ob*/*ob* mice.

(B) The numbers of regulations affected by metabolites as substrates and products in *ob*/*ob* mice at *ad libitum* feeding and 16 h-fasting[1].

(C) KEGG enrichment analysis on differential protein regulations divided into 9 groups by *ad libitum* feeding and 16 h-fasting[1].

(D) Venn plots of differential expressed molecules, TF, Enzyme mRNA, Enzyme protein, and Metabolite, in differential regulatory trans-omic networks of metabolic reactions at *ad libitum* feeding and 16 h-fasting[1].

For allosteric regulators, 400 inhibitors increased at 16 h-fasting while decreased at *ad libitum* feeding in *ob*/*ob* mice (Figure S6A). By contrast to allosteric regulation, for substrate and product regulation, decreased metabolites are dominant at *ad libitum* feeding, whereas increased metabolites are the dominant substrates and products in *ob*/*ob* mice at 16 h-fasting (Figure S6B). For enzyme protein regulation, metabolic reactions in metabolic pathways such as Biosynthesis of unsaturated fatty acids, TCA cycle, Fatty acid degradation, Glycolysis/Gluconeogenesis were activated in *ob*/*ob* mice at both *ad libitum* feeding and 16 h-fasting (Figure S6C). Metabolic reactions in BCAA degradation, Fatty acid degradation, Fatty acid elongation, Propanoate metabolism, were activated in *ob*/*ob* mice only at *ad libitum* feeding, indicating more dysregulated enzyme protein regulations at *ad libitum* feeding.

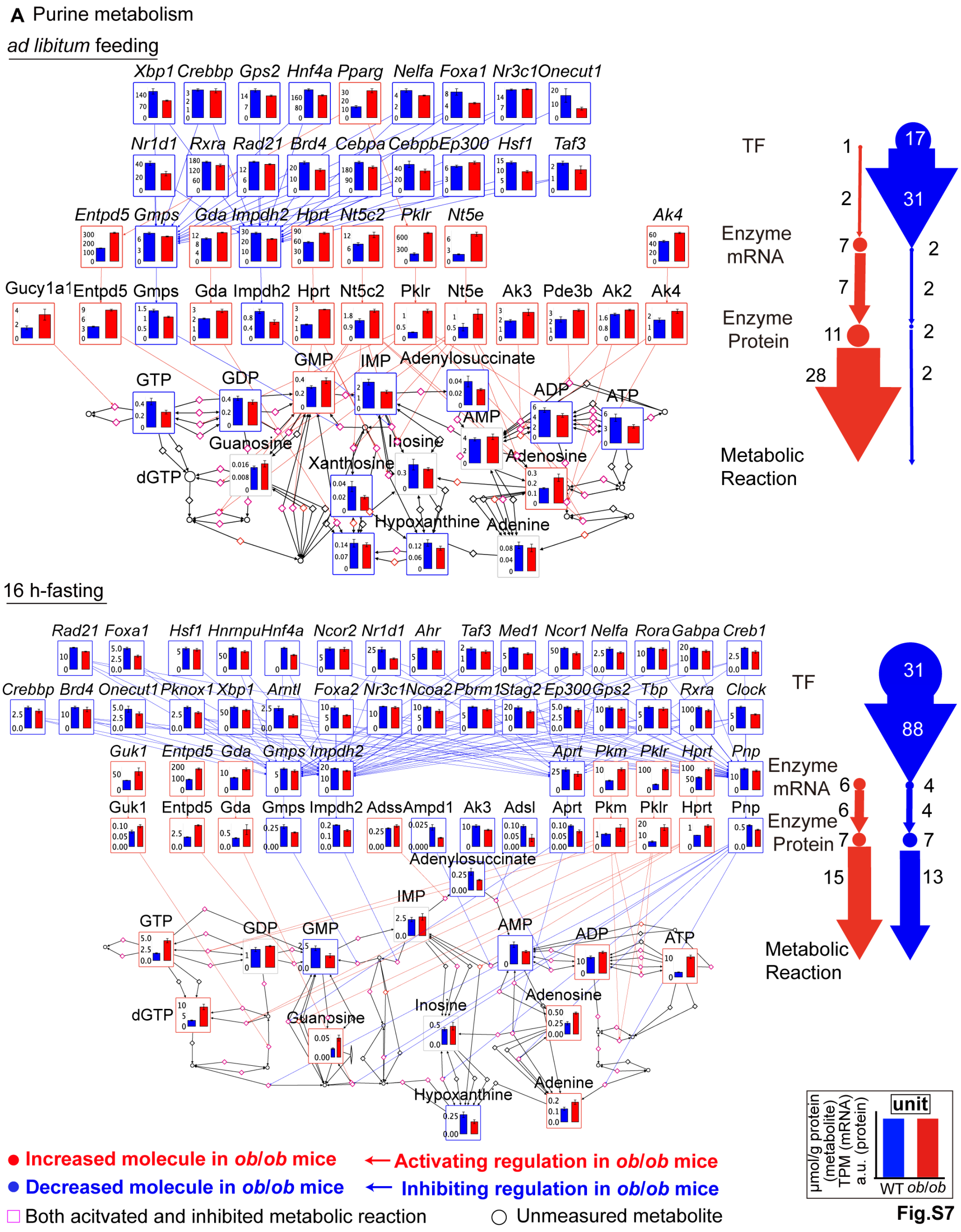

A Purine metabolism
ad libitum feeding
Xbp1 Crebbp Gps2 Hnf4a Pparg Nelfa Foxa1 Nr3c1 Onecut1
Nr1d1 Rxra Rad21 Brd4 Cebpa Cebpb Ep300 Hsf1 Taf3
Entpd5 Gmps Gda Impdh2 Hprt Nt5c2 Pklr Nt5e Ak4
Gucy1a1 Entpd5 Gmps Gda Impdh2 Hprt Nt5c2 Pklr Nt5e Ak3 Pde3b Ak2 Ak4
GTP GDP GMP IMP Adenylosuccinate ADP ATP AMP
Guanosine Inosine Xanthosine Adenosine dGTP Hypoxanthine Adenine
TF
Enzyme mRNA
Enzyme Protein
Metabolic Reaction
1 2 7 7 11 28
17 31 2 2 2 2
16 h-fasting
Rad21 Foxa1 Hsf1 Hnrnpu Hnf4a Ncor2 Nr1d1 Ahr Taf3 Med1 Ncor1 Nelfa Rora Gabpa Creb1
Crebbp Brd4 Onecut1 Pknox1 Xbp1 Arntl Foxa2 Nr3c1 Ncoa2 Pbrm1 Stag2 Ep300 Gps2 Tbp Rxra Clock
Guk1 Entpd5 Gda Gmps Impdh2 Aprt Pkm Pklr Hprt Pnp
Guk1 Entpd5 Gda Gmps Impdh2 Adss Ampd1 Ak3 Adsl Aprt Pkm Pklr Hprt Pnp
Adenylosuccinate IMP GTP GDP GMP AMP ADP ATP
dGTP Guanosine Inosine Adenosine Hypoxanthine Adenine
TF
Enzyme mRNA
Enzyme Protein
Metabolic Reaction
6 6 7 15
31 88 4 4 7 13
● Increased molecule in ob/ob mice
● Decreased molecule in ob/ob mice
□ Both acitvated and inhibited metabolic reaction
← Activating regulation in ob/ob mice
← Inhibiting regulation in ob/ob mice
○ Unmeasured metabolite
unit
μmol/g protein (metabolite)
TPM (mRNA)
a.u. (protein)
WT ob/ob

Fig.S7

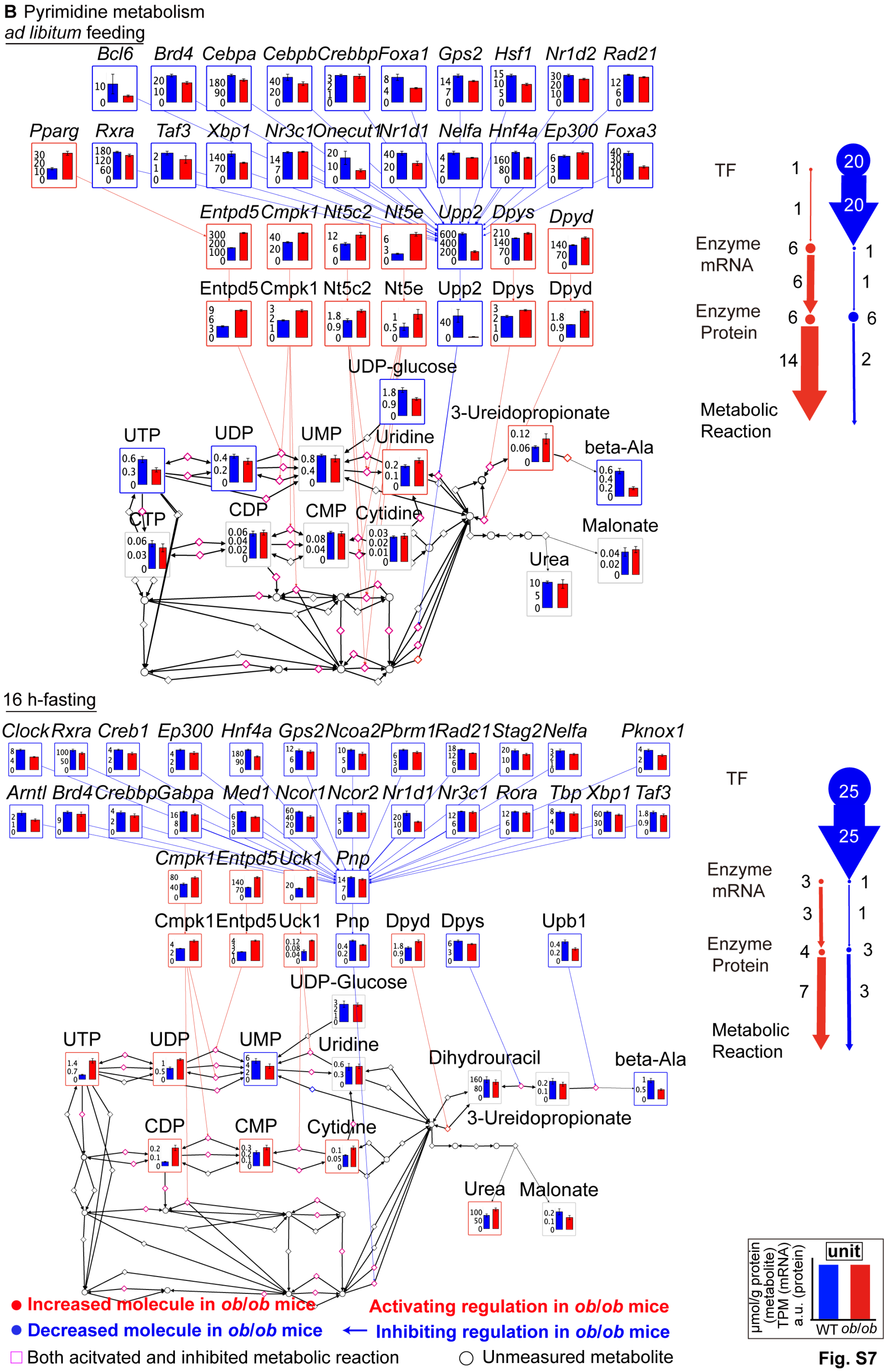
**B** Pyrimidine metabolism
ad libitum feeding
Bcl6 Brd4 Cebpa Cebpb Crebbp Foxa1 Gps2 Hsf1 Nr1d2 Rad21
Pparg Rxra Taf3 Xbp1 Nr3c1 Onecut1 Nr1d1 Nelfa Hnf4a Ep300 Foxa3
Entpd5 Cmpk1 Nt5c2 Nt5e Upp2 Dpys Dpyd
Entpd5 Cmpk1 Nt5c2 Nt5e Upp2 Dpys Dpyd
UDP-glucose
3-Ureidopropionate
UTP UDP UMP Uridine
beta-Ala
CTP CDP CMP Cytidine
Malonate
Urea
TF
Enzyme mRNA
Enzyme Protein
Metabolic Reaction
16 h-fasting
Clock Rxra Creb1 Ep300 Hnf4a Gps2 Ncoa2 Pbrm1 Rad21 Stag2 Nelfa Pknox1
Arntl Brd4 Crebbp Gabpa Med1 Ncor1 Ncor2 Nr1d1 Nr3c1 Rora Tbp Xbp1 Taf3
Cmpk1 Entpd5 Uck1 Pnp
Cmpk1 Entpd5 Uck1 Pnp Dpyd Dpys Upb1
UDP-Glucose
UTP UDP UMP Uridine
Dihydrouracil
beta-Ala
3-Ureidopropionate
CDP CMP Cytidine
Urea Malonate
TF
Enzyme mRNA
Enzyme Protein
Metabolic Reaction
unit
µmol/g protein (metabolite)
TPM (mRNA)
a.u. (protein)
WT ob/ob
● Increased molecule in ob/ob mice
● Decreased molecule in ob/ob mice
□ Both acitvated and inhibited metabolic reaction
Activating regulation in ob/ob mice
← Inhibiting regulation in ob/ob mice
○ Unmeasured metabolite

**Fig. S7**

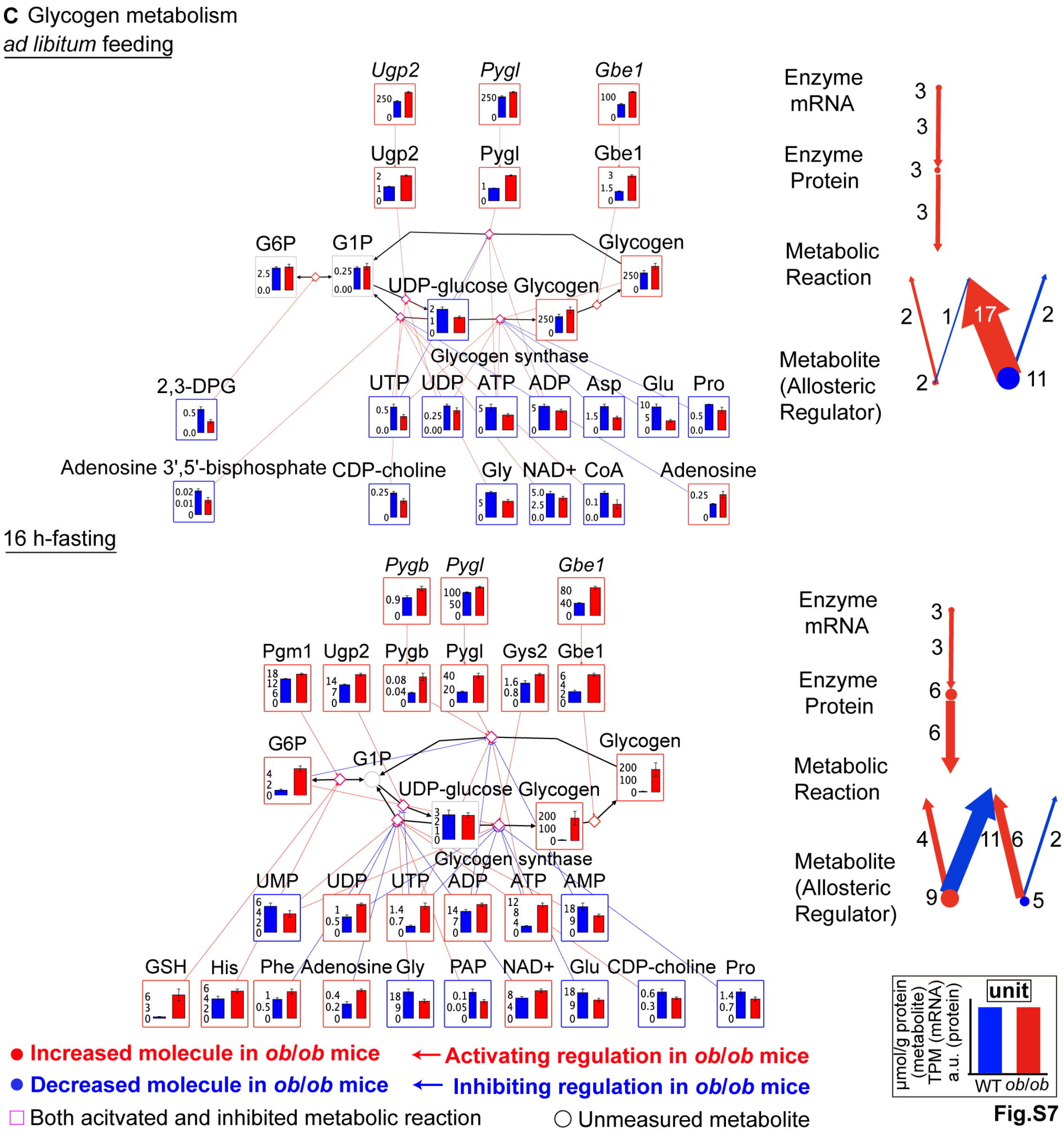
C Glycogen metabolism
ad libitum feeding
Ugp2
Pygl
Gbe1
G6P
G1P
Glycogen
UDP-glucose
Glycogen synthase
2,3-DPG
UTP
UDP
ATP
ADP
Asp
Glu
Pro
Adenosine 3',5'-bisphosphate
CDP-choline
Gly
NAD+
CoA
Adenosine
Enzyme mRNA
Enzyme Protein
Metabolic Reaction
Metabolite (Allosteric Regulator)
16 h-fasting
Pygb
Pgm1
Gys2
UMP
AMP
GSH
His
Phe
PAP
Increased molecule in ob/ob mice
Decreased molecule in ob/ob mice
Both acitvated and inhibited metabolic reaction
Activating regulation in ob/ob mice
Inhibiting regulation in ob/ob mice
Unmeasured metabolite
unit
μmol/g protein (metabolite)
TPM (mRNA)
a.u. (protein)
WT ob/ob

Fig.S7

**D** Fatty acid synthesis

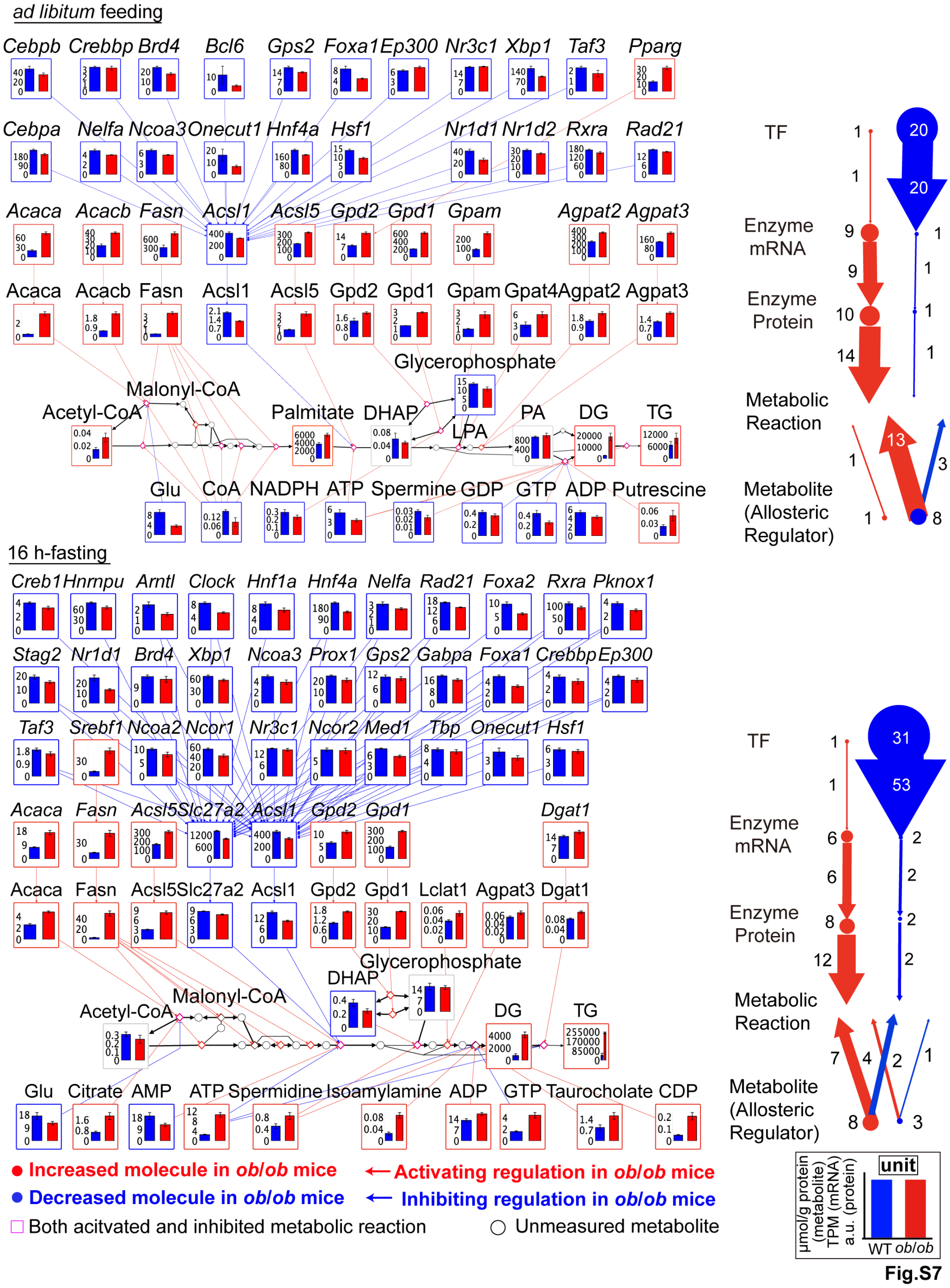


**Fig.S7**

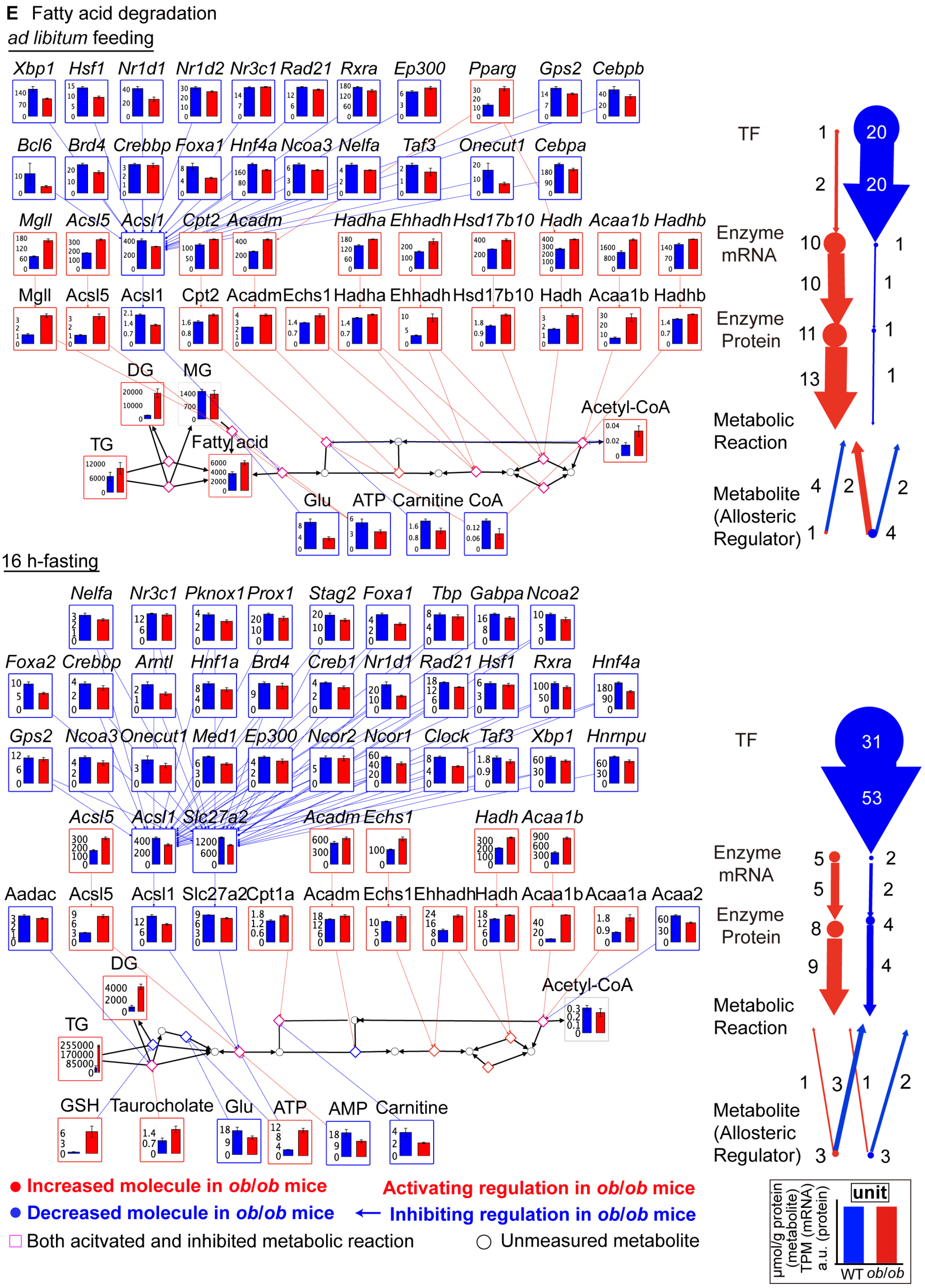

E Fatty acid degradation
ad libitum feeding
Xbp1 Hsf1 Nr1d1 Nr1d2 Nr3c1 Rad21 Rxra Ep300 Pparg Gps2 Cebpb
Bcl6 Brd4 Crebbp Foxa1 Hnf4a Ncoa3 Nelfa Taf3 Onecut1 Cebpa
Mgll Acsl5 Acsl1 Cpt2 Acadm Hadha Ehhadh Hsd17b10 Hadh Acaa1b Hadhb
Mgll Acsl5 Acsl1 Cpt2 Acadm Echs1 Hadha Ehhadh Hsd17b10 Hadh Acaa1b Hadhb
DG
MG
TG
Fatty acid
Acetyl-CoA
Glu
ATP
Carnitine
CoA
TF
Enzyme mRNA
Enzyme Protein
Metabolic Reaction
Metabolite (Allosteric Regulator)
16 h-fasting
Nelfa Nr3c1 Pknox1 Prox1 Stag2 Foxa1 Tbp Gabpa Ncoa2
Foxa2 Crebbp Arntl Hnf1a Brd4 Creb1 Nr1d1 Rad21 Hsf1 Rxra Hnf4a
Gps2 Ncoa3 Onecut1 Med1 Ep300 Ncor2 Ncor1 Clock Taf3 Xbp1 Hnrnpu
Acsl5 Acsl1 Slc27a2 Acadm Echs1 Hadh Acaa1b
Aadac Acsl5 Acsl1 Slc27a2 Cpt1a Acadm Echs1 Ehhadh Hadh Acaa1b Acaa1a Acaa2
DG
TG
Acetyl-CoA
GSH
Taurocholate
Glu
ATP
AMP
Carnitine
● Increased molecule in ob/ob mice
● Decreased molecule in ob/ob mice
□ Both acitvated and inhibited metabolic reaction
Activating regulation in ob/ob mice
← Inhibiting regulation in ob/ob mice
○ Unmeasured metabolite
unit
μmol/g protein (metabolite)
TPM (mRNA)
a.u. (protein)
WT ob/ob


**Fig.S7**

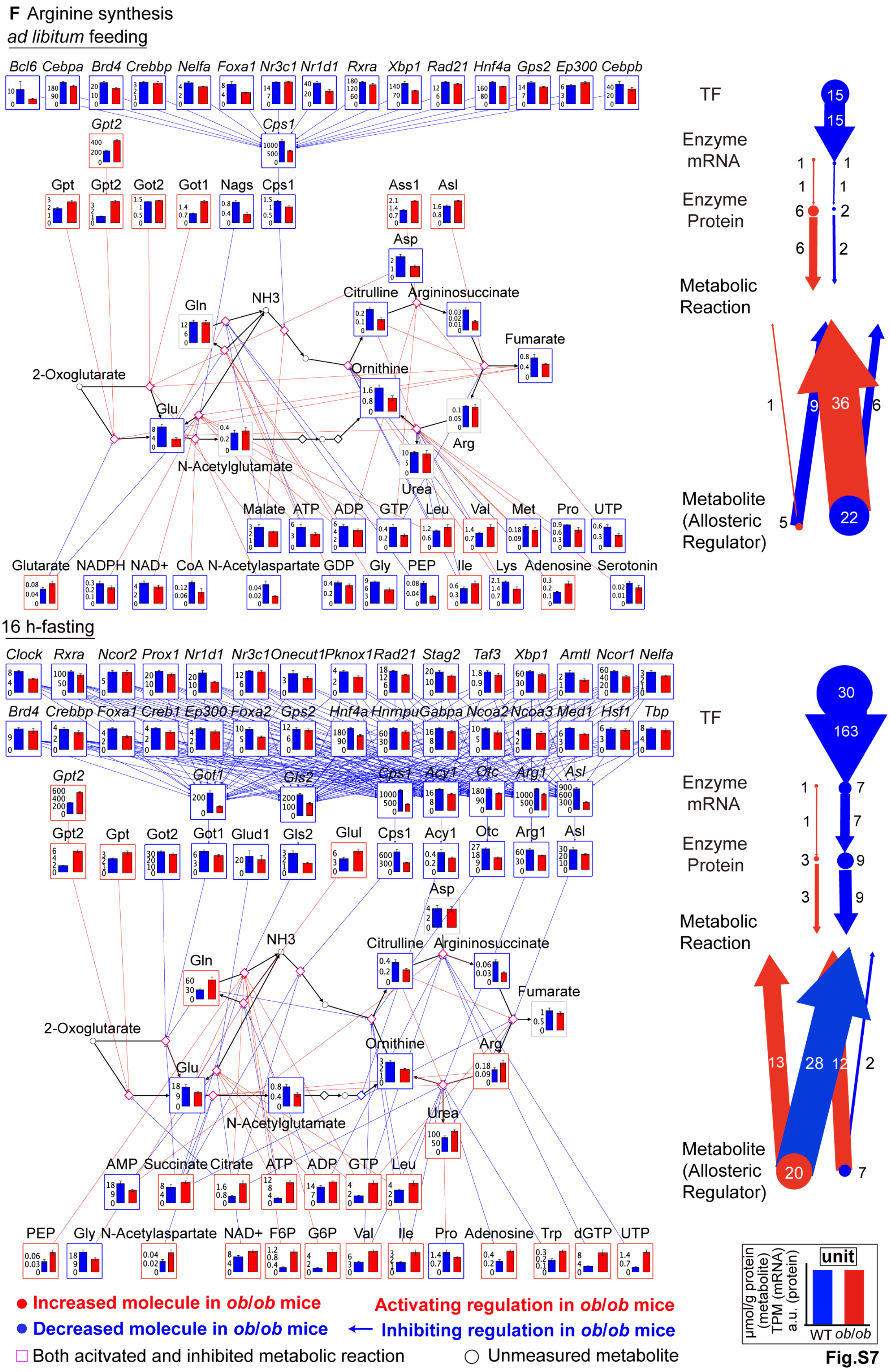
F Arginine synthesis
ad libitum feeding
Bcl6 Cebpa Brd4 Crebbp Nelfa Foxa1 Nr3c1 Nr1d1 Rxra Xbp1 Rad21 Hnf4a Gps2 Ep300 Cebpb
Gpt2
Cps1
Gpt Gpt2 Got2 Got1 Nags Cps1
Ass1 Asl
Asp
NH3
Gln
Citrulline
Argininosuccinate
Fumarate
Ornithine
2-Oxoglutarate
Glu
Arg
N-Acetylglutamate
Urea
Malate ATP ADP GTP Leu Val Met Pro UTP
Glutarate NADPH NAD+ CoA N-Acetylaspartate GDP Gly PEP Ile Lys Adenosine Serotonin
TF
Enzyme mRNA
Enzyme Protein
Metabolic Reaction
Metabolite (Allosteric Regulator)
16 h-fasting
Clock Rxra Ncor2 Prox1 Nr1d1 Nr3c1 Onecut1 Pknox1 Rad21 Stag2 Taf3 Xbp1 Amtl Ncor1 Nelfa
Brd4 Crebbp Foxa1 Creb1 Ep300 Foxa2 Gps2 Hnf4a Hnrnpu Gabpa Ncoa2 Ncoa3 Med1 Hsf1 Tbp
Gpt2 Got1 Gls2 Cps1 Acy1 Otc Arg1 Asl
Gpt2 Gpt Got2 Got1 Glud1 Gls2 Glul Cps1 Acy1 Otc Arg1 Asl
Asp
NH3
Gln
Citrulline
Argininosuccinate
Fumarate
2-Oxoglutarate
Ornithine
Arg
Glu
N-Acetylglutamate
Urea
AMP Succinate Citrate ATP ADP GTP Leu
PEP Gly N-Acetylaspartate NAD+ F6P G6P Val Ile Pro Adenosine Trp dGTP UTP
TF
Enzyme mRNA
Enzyme Protein
Metabolic Reaction
Metabolite (Allosteric Regulator)
Increased molecule in ob/ob mice
Decreased molecule in ob/ob mice
Both acitvated and inhibited metabolic reaction
Activating regulation in ob/ob mice
Inhibiting regulation in ob/ob mice
Unmeasured metabolite
unit
μmol/g protein (metabolite)
TPM (mRNA)
a.u. (protein)
WT ob/ob
Fig.S7

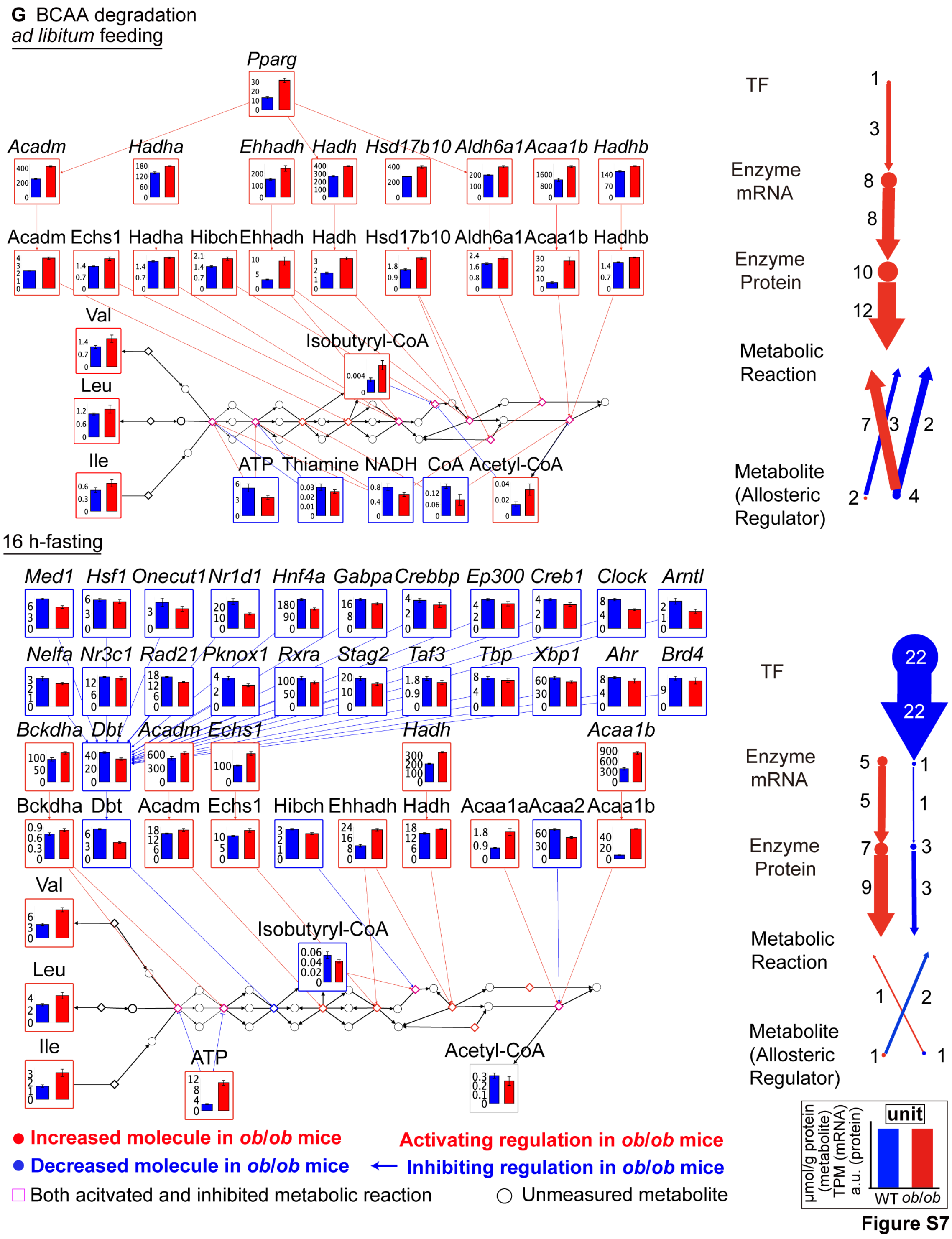


**Figure S7. Comparison of metabolic pathways between *ad libitum* feeding and 16 h-fasting in the liver of WT and *ob*/*ob* mice. Related to Figures 6 and 7.**

Metabolic reactions, molecules, and regulations in (A) Purine metabolism, (B) Pyrimidine metabolism, (C) Glycogen metabolism, (D) Fatty acid synthesis, (E) Fatty acid degradation, (F) Arginine synthesis, (G) BCAA degradation, extracted from the differential regulatory trans-omic network at *ad libitum* feeding (top-left) and 16 h-fasting (bottom-left)[1].

Bar plots are shown on the corresponding nodes as the means and SEMs of replicates in WT and *ob*/*ob* mice. For metabolites as substrates or products, we showed bar plots for all measured molecules. The colors of frame and edge indicate the changed direction. Diamond nodes indicate metabolic reactions. Black edges indicate the reversible metabolic reactions between substrates and products. Black circles indicate the metabolites were not measured. From metabolite to metabolic reaction, only allosteric regulations are colored. We counted the numbers of differentially expressed TF, Enzyme mRNA, Enzyme Protein, Metabolite, Metabolite (Allosteric Regulator) and differential regulations in each metabolic pathway at *ad libitum* feeding (top-right) and 16 h-fasting (bottom-right). We used the size of node and the width of edge to indicate them. For those with more than 30 or less than 2, we used one size respectively (>30, 30, <2, 2) in case too big or too small nodes and edges. Specific numbers were shown around nodes and edges. Red, increased molecules or activating regulations in *ob*/*ob* mice; blue, decreased molecules or inhibiting regulations in *ob*/*ob* mice; gray, unchanged molecules between WT and *ob*/*ob* mice; magenta, metabolic reaction is both activated and inhibited in *ob*/*ob* mice.

For purine metabolism, although there are more activating and less inhibiting enzyme protein regulations at *ad libitum* feeding (28 activating regulations and 2 inhibiting regulations) than 16 h-fasting (15 activating regulations and 13 inhibiting regulations), more decreased metabolites (8 metabolites) were observed at *ad libitum* feeding than at 16 h-fasting (4 metabolites). IMP and GDP decreased in *ob*/*ob* mice only at *ad libitum* feeding (Figure S7A). Guanosine and adenine increased, and AMP decreased only at 16 h-fasting. GMP increased at *ad libitum* feeding while decreased at 16 h-fasting. For the changed enzyme proteins, 11 of 13 enzyme proteins increased at *ad libitum* feeding. 7

increased and 7 decreased enzyme proteins at 16 h-fasting. The number of inhibiting TF regulations and enzyme protein regulations in *ob*/*ob* mice at *ad libitum* feeding is less than 16 h-fasting.

For pyrimidine metabolism, Uridine and 3-Ureidopropionate increased, and UDP-glucose decreased in *ob*/*ob* mice only at *ad libitum* feeding (Figure S7B). Urea, CDP, cytidine, and CMP increased; UMP decreased only at 16 h-fasting. More activating enzyme protein regulations in *ob*/*ob* mice at *ad libitum* feeding (14 regulations) than 16 h-fasting (7 regulations). In general, pyrimidine is likely inhibited in *ob*/*ob* mice at *ad libitum* feeding while activated at 16 h-fasting. These results indicate the opposite dysregulation of pyrimidine metabolism between *ad libitum* feeding and 16 h-fasting.

For glycogen metabolism, Ugp2, Pygl, and Gbe1 increased in *ob*/*ob* mice at both *ad libitum* feeding and 16 h-fasting (Figure S7C). Pgm1, Pygb and Gys2 increased only at 16 h-fasting. No TF regulation and inhibiting enzyme protein regulation at both *ad libitum* feeding and 16 h-fasting. The number of activating enzyme protein regulations at 16 h-fasting (6 regulations) is larger than *ad libitum* feeding (3 regulations). Eleven decreased allosteric regulators caused seventeen activating allosteric regulations in *ob*/*ob* mice at ad libitum feeding. Here, glycogen synthesis was likely to dysregulated with the decreased substrate UDP-glucose in *ob*/*ob* mice at ad libitum feeding.

For fatty acid synthesis, TG and DG increased in *ob*/*ob* mice at both *ad libitum* feeding and 16 h-fasting (Figure S7D). Acetyl-CoA increased, and glycerol-3P decreased only at *ad libitum* feeding. DHAP decreased only at 16 h-fasting. More activating enzyme protein regulations and less inhibiting TF regulations were observed in *ob*/*ob* mice at *ad libitum* feeding (14 activating enzyme protein regulations and 20 inhibiting TF regulations) than 16 h-fasting (12 activating enzyme protein regulations and 53 inhibiting TF regulations). 8 decreased metabolites as allosteric regulators caused 13 activating allosteric regulations at *ad libitum* feeding. 8 increased metabolites as allosteric regulators caused 7 activating allosteric regulations and 4 inhibiting allosteric regulations. Fatty acid synthesis is likely activated by enzyme proteins and allosteric regulators with increased DG and TG in *ob*/*ob* mice at *ad libitum* feeding and 16 h-fasting.

For fatty acid degradation, TG and DG increased in *ob*/*ob* mice at both *ad libitum* feeding and 16 h-fasting (Figure S7E). Acetyl-CoA increased only at *ad libitum* feeding. 11 increased and 1 decreased enzyme proteins at *ad libitum* feeding. 8 increased and 4 decreased enzyme proteins at 16 h-fasting. More activating enzyme proteins regulations, and less inhibiting TF regulations were observed in *ob*/*ob* mice at *ad libitum* feeding (13 activating enzyme protein regulations and 20 inhibiting TF regulations) than 16 h-fasting (9 activating enzyme protein regulations and 53 inhibiting TF regulations). For the changed metabolites, the total number of allosteric regulations in fatty acid degradation (8 allosteric regulations at *ad libitum* feeding and 7 allosteric regulations at 16 h-fasting) is less than in other metabolic pathways. In conclusion, fatty acid degradation is likely activated by enzyme proteins with increased Acetyl-CoA in *ob*/*ob* mice at *ad libitum* feeding.

For arginine synthesis, Arginine, Gln, and Urea increased, N-Acetylglutamate decreased in *ob*/*ob* mice only at 16 h-fasting (Figure S7F). Asp and Fumarate decreased only at *ad libitum* feeding. Cps1 catalyzes the synthesis of carbamoyl phosphate, which is the first step of urea cycle. Cps1 decreased, which may be the reason that all the changed metabolites decreased at *ad libitum* feeding. 22 decreased allosteric regulators caused 36 activating allosteric regulations in *ob*/*ob* mice at *ad libitum* feeding. No activating TF regulation was observed at both *ad libitum* feeding and 16 h-fasting. Less inhibiting TF and enzyme protein regulations, more activating enzyme protein regulations were observed in *ob*/*ob* mice at *ad libitum* feeding (15 inhibiting TF regulations and 6 activating enzyme protein regulations) than 16 h-fasting (163 inhibiting TF regulations and 3 activating enzyme protein regulations). Taken together, it is likely that arginine synthesis is inhibited in *ob*/*ob* mice by enzyme protein, Cps1, at *ad libitum* feeding.

For BCAA degradation (Figure S7G), the total number of allosteric regulations at *ad* *libitum* feeding (12 allosteric regulations) was larger than that at 16 h-fasting (3 allosteric regulations). The changed metabolites, all of 5 increased in *ob*/*ob* mice at *ad libitum* feeding. Isobutyryl-CoA increased at *ad libitum* feeding but decreased at 16 h-fasting. No inhibiting TF regulation and enzyme protein regulation was observed in *ob*/*ob* mice at *ad libitum* feeding. The number of activating TF regulations and enzyme protein regulations in *ob*/*ob* mice at *ad libitum* is larger than 16 h-fasting. BCAA degradation is likely to be activated by enzyme proteins with increased Acetyl-CoA in *ob*/*ob* mice at *ad libitum* feeding.
